# Pathogen genomic surveillance as a scalable framework for precision phage therapy

**DOI:** 10.1101/2024.06.15.599013

**Authors:** Mihály Koncz, Tamás Stirling, Hiba Hadj Mehdi, Orsolya Méhi, Bálint Eszenyi, András Asbóth, Gábor Apjok, Ákos Tóth, László Orosz, Bálint Márk Vásárhelyi, Eszter Ari, Lejla Daruka, Tamás Ferenc Polgár, György Schneider, Sif Aldin Zalokh, Mónika Számel, Gergely Fekete, Balázs Bohár, Karolina Nagy Varga, Ádám Visnyovszki, Edit Székely, Monica-Sorina Licker, Oana Izmendi, Carmen Costache, Ina Gajic, Bojana Lukovic, Szabolcs Molnár, Uzonka Orsolya Szőcs-Gazdi, Csilla Bozai, Marina Indreas, Katalin Kristóf, Charles Van der Henst, Anke Breine, Csaba Pál, Balázs Papp, Bálint Kintses

**Author notes:** These authors contributed equally (Mihály Koncz, Tamás Stirling, Hiba Hadj Mehdi, Orsolya Méhi). Correspondence (B.K.), (B.P.).

## Abstract

Phage therapy is gaining increasing interest in the fight against critically resistant nosocomial pathogens. However, the narrow host range of bacteriophages hampers the development of broadly effective phage therapeutics and demands precision approaches. Here we combine large-scale phylogeographical analysis with high-throughput phage typing to guide the development of precision phage cocktails targeting carbapenem-resistant *Acinetobacter baumannii,* a top-priority pathogen. Our analysis reveals that a few strain types dominate infections in each world region, with their geographical distribution remaining stable within six years. As we demonstrate in Eastern Europe, this spatio-temporal distribution enables preemptive preparation of region-specific phage collections that target most local infections. Finally, we showcase the efficacy of a four-phage cocktail against the most prevalent strain type in both *in vitro* and *in vivo* animal infection models. Ultimately, genomic surveillance identifies patients benefiting from the same phages across geographical scales, thus providing a scalable framework for precision phage therapy.

**Highlights:**

- A few carbapenem-resistant *Acinetobacter baumannii* types dominate infections worldwide
- Phylogeography reveals stable strain composition of individual countries over a six-year period
- This spatio-temporal distribution allows preemptive preparation of region-specific phage collections
- A four-phage cocktail is efficacious against the most prevalent strain type in Europe.

## Introduction

The uncontrollable rise of antibiotic resistance will become the leading cause of human mortality by the year 2050 unless proactive measures are implemented.^1^ Of particular concern is the widespread presence of antibiotic-resistant bacteria in healthcare settings.^2^ Due to the limited number of effective antibiotic options, these infections often produce severe clinical outcomes.^3^ Moreover, as the conventional antibiotic development pipeline is drained, this situation will most likely not improve in the near future.^4,5^ Therefore, it is imperative to find alternative therapeutic approaches, with phage therapy emerging as a prominent candidate.^6^

While there have been numerous compassionate use cases where phages successfully tackled infections that resisted all conventional antibiotic treatments,^7,8,9,10,11,12^ the translation of this potential into positive patient outcomes on a broader scale has been progressing slowly.^13,14^ This limitation stems from the narrow host range of bacteriophages, which complicates their broad therapeutic applications.^15^ Specifically, many resistant bacterial species consist of hundreds of strain types, each characterised by distinct cell surface structures and defence mechanisms that restrict the host range of bacteriophages to a subset of bacterial strain types.^16,17^ In response to this challenge, two primary approaches have emerged.^14^ The first one combines multiple phages into a fixed composition for broader bacterial coverage, followed by the initiation of clinical trials aimed at obtaining marketing authorisation.^18,19^ While several such attempts are currently in progress, as of now, no registered medicinal phage products have reached the market.^20^ The second strategy is personalised treatment, where the patient’s bacterial sample is screened to identify the most effective phages from collections of bacteriophages.^14,15^ While this approach has demonstrated promising clinical efficiency,^21^ it comes with its own set of challenges, including limited throughput due to time-consuming phage production and regulatory approval hurdles that often follow the diagnosis of the infection.^22^ Overall, irrespective of the chosen methodology, further development of the field will largely depend on systematic procedures that identify patients requiring the same bacteriophages on a large scale.^23^

In recent years, genomic surveillance has matured into an efficient tool to track pathogens at an unprecedented scale.^24^ Advances in sequencing technologies have revealed the genomes of hundreds of thousands of bacterial isolates.^25^ This wealth of data now presents a unique opportunity to gain insights into the distribution and transmission patterns of top-priority antibiotic-resistant pathogens on the World Health Organisation’s (WHO) list for which innovative therapies are urgently needed.^26^ In fact, genomic surveillance has established that such infections can originate within healthcare facilities.^27^(ref) This type of transmission raises the prospect of forecasting the likely causative agents of forthcoming infections and preemptively tailoring phage cocktails to combat them. The ability to pre-emptively prepare suitable phages could dramatically reduce the time required for administering personalized therapy following diagnosis. Moreover, knowing the geographic distribution of top-priority pathogens is crucial to facilitate study recruitment to achieve clinical validation and cost-effectiveness for phage therapy. Despite this immense potential, genomic surveillance remains largely untapped in phage therapy.^23^ Our proposition is that genomic surveillance can lay the groundwork for a scalable precision phage therapy (Fig 1). In support of this, we conducted a systematic analysis, with an initial >15,000 *Acinetobacter baumannii* (*A. baumannii)* genome sequences from around the world. We chose to focus on this particular bacterial species as carbapenem-resistant *A. baumannii* (CRAB) is recognized by WHO as one of the critical priority pathogens for which innovative therapies are urgently needed,^26^ (ref) with an alarming mortality rate of 25-60% and more than 100,000 deaths globally in 2019.^28,29^ (ref) Meanwhile, the phage host range of this bacterial species is especially narrow. In fact, a recent estimate suggests that it may require a phage library of more than 300 phages to effectively cover the majority of the clinical isolates.^30^ To investigate this issue, we used large-scale phylogeographic analyses in combination with experimental phage discovery and high-throughput phage typing to demonstrate that 90% of the infections can be attributed to a limited number of strain types in each world region, which can be effectively addressed by a small set of bacteriophages. Furthermore, the causative CRAB strains are predictable within a six-year timeframe for individual countries, and countries with identical CRAB strain types can be rapidly identified. These combined capabilities enable proactive measures, including the development of region-specific phage collections and the formulation of phage cocktails designed to target specific strain types in a highly precise manner.

**Figure 1.**
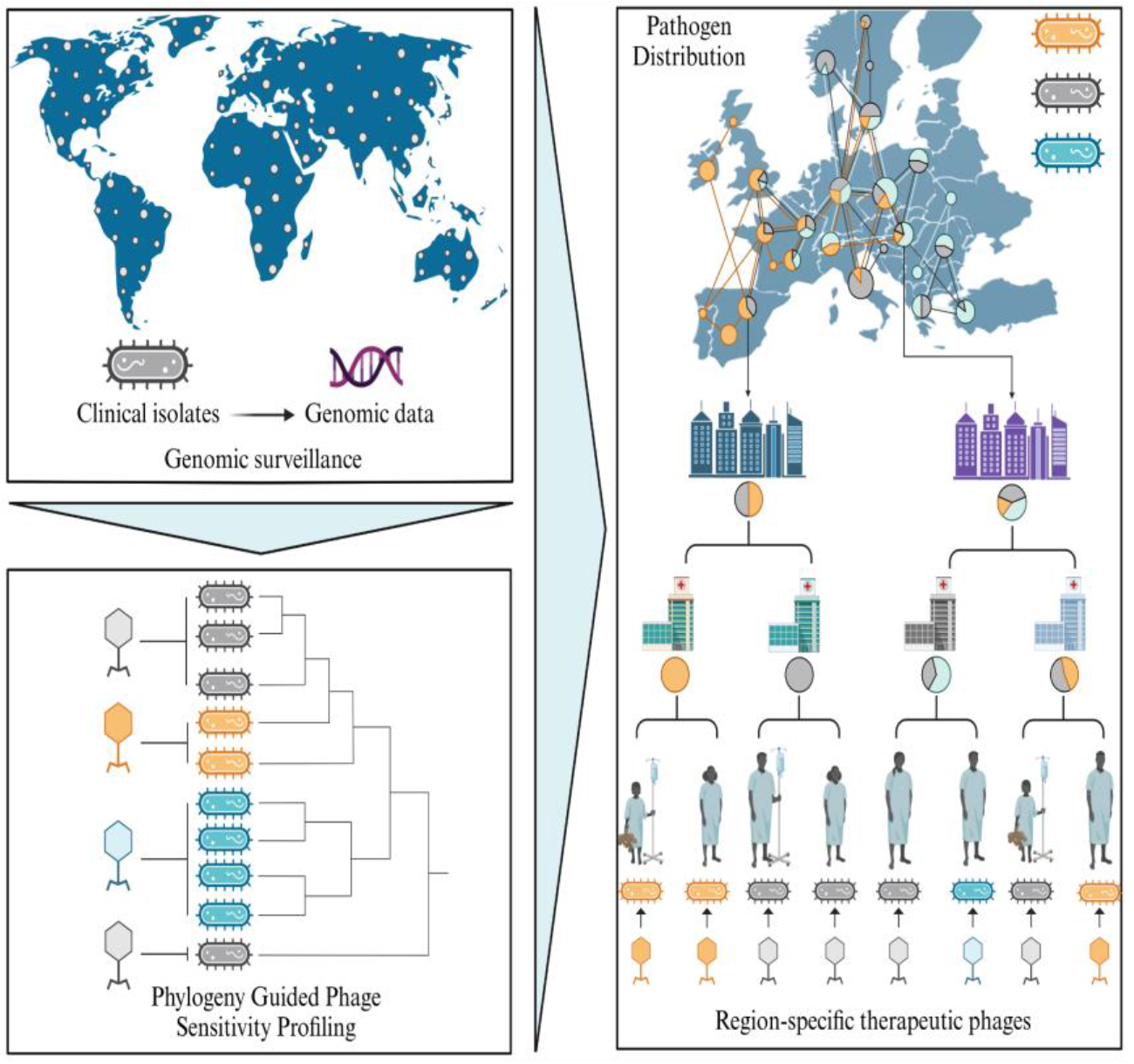
A pathogen genomic surveillance framework to inform precision phage therapy. The process involves: (1) Collecting isolates of the target pathogen from surveillance programmes and laboratory networks; using information on genome sequence, and location and time of isolation; (2) Identifying dominant pathogen strain types requiring distinct bacteriophages based on pathogen genomics and phage sensitivity profiling; (3) Utilising the spatio-temporal distribution of the identified dominant pathogen types to design region-specific phage therapy interventions. These interventions maximise the therapeutic potential for the largest number of patients across geographical scales.

## Results

### The global distribution of CRAB types

Our first objective was to assess the prevalence and geographical distribution of the CRAB strains at both regional and global scales. To accomplish this, we initiated our analysis by retrieving all publicly available *A. baumannii* genomes from the NCBI database (15,410 genomes as of 09/2022), originating from 85 countries across five continents (Supplementary Table 1). Given the limited representation of genomic samples from Eastern and Southern European regions (Supplementary Table 1), we additionally sequenced 419 *A. baumannii* clinical isolates collected between 2011 and 2022 from 44 healthcare facilities located in 34 cities across five neighbouring countries in these regions (Hungary (n=253), Romania (n=120), Serbia (n=28), Montenegro (n=9), and Bosnia and Herzegovina (n=9)). Subsequently, we refined our dataset, narrowing down the initial 15,829 genomes to a final set of 11,129 genomes (Supplementary Table 1). The selection criteria had considerations such as genome quality, human origin, and specified information regarding the place and time of isolation (Methods, Fig S1). Finally, we screened these genomes for genetic determinants that collectively characterise CRAB traits, including resistance genes to carbapenems and other commonly used antibiotic classes (Methods). Our analysis revealed that 80.1% of the 11,129 genomes can be considered CRAB according to these criteria (Fig 2A). As expected, there was a noticeable global increase in the relative proportion of CRAB genomes observed between the years 2000 and 2022 (Fig 2B).

**Figure 2.**
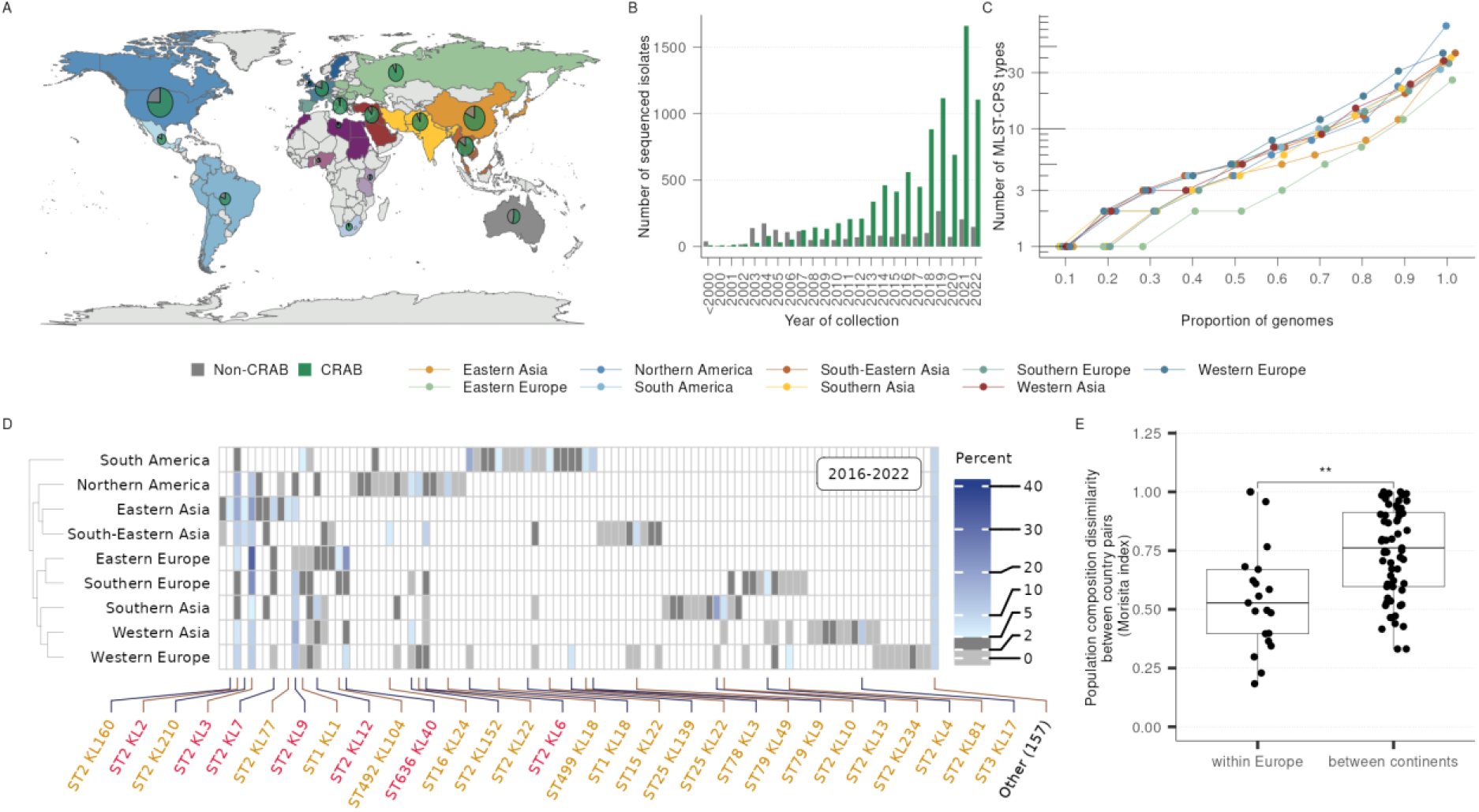
The global geographic distribution of CRAB strains. **A,** Geographical distribution of 11,129 good quality, human-derived *A. baumannii* whole genome sequences (Methods, Supplementary Table 1). The size of each pie chart corresponds to the number of sequenced isolates in the specific world region; colour-coding describes the ratio of CRAB (green) and non-CRAB (grey) sequences as defined by the presence of genetic determinants of resistance (Methods). **B,** Global increase in the relative proportion of CRAB genomes over time between the years 2000 and 2022. **C,** Cumulative proportion of MLST-CPS types among 3,150 concurrent (2016 - 2022) CRAB genomes in the nine studied world regions. Genomes are downsampled to remove sampling bias (Methods). Each point represents the number of most prevalent MLST-CPS types accounting for a certain proportion of isolates collected in a region (Supplementary Table 2). **D,** Relative prevalence of CRAB MLST-CPS types in the nine world regions. The most prevalent strains accounting for 90% of the genomes in a region are shown individually, and the remaining 10% are pooled (Other). Colours indicate the relative prevalence ranges as follows: 0 (white), 0-2% (light grey), 2-5% (dark grey), and 5-40% (shades of blue). MLST-CPS types classified as prevalent or global are marked with orange and red labels, respectively (Methods). **E,** Population composition is more similar between pairs of countries in Europe compared with pairs of countries from different continents (two-sided two-sample Wilcoxon rank-sum test, 7 European countries, 7 non-European countries, *N*_within_ = 21, *N*_between_ = 63, *p* = 0.002). Population compositions are derived from the relative prevalence of MLST-CPS types (Methods). The higher and lower ends of the boxes indicate 75th and 25th percentiles, respectively, the line inside the box indicates the median of the distribution, and the whiskers were drawn using standard 1.5 interquartile ranges. All data are available in Supplementary Tables 1, 2, and 3.

As a next step, we grouped the genomes into CRAB types by employing a dual genetic typing approach, capturing complementary aspects of phage susceptibility. This was achieved by combining multilocus sequence typing (MLST) and cell surface capsular polysaccharide (CPS) typing.^31,32^ MLST clusters evolutionary-related genomes into sequence types (ST) based on the sequence of multiple housekeeping genes.^33^ In contrast, CPS typing identifies specific genomic determinants responsible for synthesising structurally different CPSs, which can function as receptors for different Acinetobacter phages.^34^ It is important to note that the genetic diversity of CPS is shaped by frequent horizontal gene transfers.^16^ Therefore, this trait evolves independently from ST. Finally, we analysed the distribution of currently (2016 - 2022) circulating CRAB types as defined by the combined MLST-CPS typing of the genomes in nine world regions (West-, East- and South-Europe; North- and South-America; West-, East- and South-East-Asia) where our sampling was sufficient to capture the majority of CRAB diversity after correcting for sampling bias (Methods, Fig S2).

In the nine world regions we studied, the contemporary global diversity of CRAB is characterised by a small number of prevalent MLST-CPS types and a large number of rare ones (Fig 2C). Specifically, in a given world region, typically 4 or 5 MLST-CPS types collectively account for 50% of the isolates, while 22 MLST-CPS types make up 90% (Fig 2C). Altogether, we identified 29 “prevalent” MLST-CPS types that have a relative prevalence in the CRAB population of at least 5% in at least one world region (Fig 2D). Nevertheless, there is substantial variation in the composition of CRAB types across different world regions. Specifically, among these 29 prevalent MLST-CPS types, only 7 exhibit a relative prevalence exceeding 2% in three or more world regions, which span across at least two continents (Fig 2D, Fig S3). This makes these 7 types not only prevalent but also globally dispersed. Finally, in large contrast to the populations of the prevalent types, the remaining 10% of the global isolates can be attributed to 157 MLST-CPS types (Fig 2D).

Then, we investigated whether countries in close geographical proximity tend to share a more similar composition of CRAB populations. For this analysis, we focused on seven European, closely situated and seven non-European, geographically distant countries with sufficient genomic sampling (Methods, Fig S4). Our analysis reveals that the MLST-CPS type composition of CRAB populations between pairs of European countries tends to exhibit more similarity compared to pairs of countries located on different continents (Fig 2E). This high similarity is primarily attributed to the presence of 10 prevalent types shared by at least three of the seven European countries, collectively representing 68% of the isolates (Fig S5). Additionally, the set of 40 country-specific MLST-CPS types constitutes only 17% of the isolates in these countries (Fig S5, Supplementary Table 1).

To summarise, the diversity within the CRAB population in each world region is characterised by a few prevalent CRAB types that are responsible for the majority of the infections and a vast number of rare strains. Additionally, while there are substantial differences in CRAB populations across world regions, countries that are geographically close to one another tend to share a more similar CRAB type composition.

### Local CRAB variants drive infections within a 6-year period

As a next step, we investigated the temporal dynamics of CRAB populations. Initially, we analysed temporal changes in the global frequencies of the 29 MLST-CPS types that are currently (2016 - 2022) prevalent (Fig 3A). Statistical modelling revealed that 11 out of 29 prevalent strains showed a significant expansion over the past 12 years on at least one continent (Methods, Fig S6, Supplementary Table 4). The largest expansion was exhibited by the ST2-KL3 type (Fig 3A), which gradually increased from 2009 onwards to become one of the three most prevalent CRAB types in seven out of the nine studied world regions (Fig S3). This expansion across multiple geographical regions implies a potential fitness advantage of ST2-KL3 over other types.

**Figure 3.**
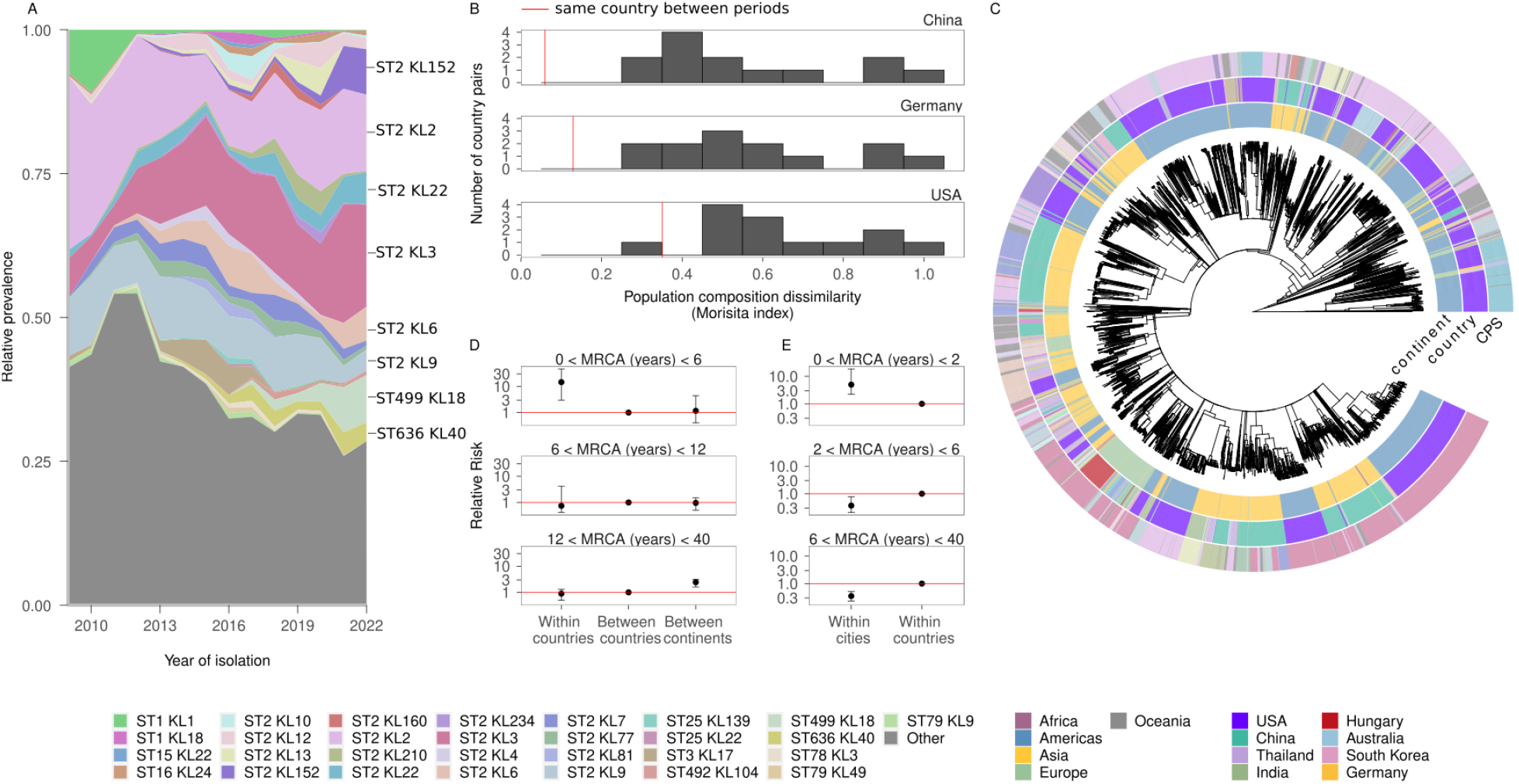
Temporal stability of CRAB populations within geographical regions. **A,** Global relative prevalence of CRAB MLST-CPS types since 2009 (2-year moving average) based on 4,860 CRAB samples following downsampling (Methods). The 29 prevalent MLST-CPS types are colour-coded, and the rest are pooled. MLST-CPS types classified as prevalent since 2016 are listed individually, the rest of the MLST-CPS types are pooled into a single category (grey). **B,** The MLST-CPS type composition of a specific country from the period between 2009 - 2015 is more similar to its contemporary (2016 - 2022) strain composition than to the contemporary composition of most other countries worldwide, with a single exception, which is the USA *vs.* China. The vertical red lines show population composition dissimilarity (Morisita index) comparisons of the same country between the two time periods. The rest of the histogram shows comparisons between the focal country and 13 other countries. **C,** Phylogenetic tree of 7,720 ST2 *A. baumannii* isolates from around the world. Heatmaps indicate CPS type, country and continent of origin. Only countries with at least 100 isolates are colour-coded. **D,** Relative risk that pairs of isolates were collected in a certain geographic location (same country, different country but same continent) compared to the reference category (different country and different continent). Each facet describes pairs of samples within a certain genetic distance range (time to the most recent common ancestor, MRCA, years). Error bars represent 90% confidence intervals, based on 100 downsampled time-calibrated trees. **E,** Higher resolution relative risks that only include isolates from the 5 European countries from which new isolates were collected in the course of this study. Error bars represent 90% confidence intervals, based on 500 downsampled time-calibrated trees. All data are available in Supplementary Tables 1, 6, and 7.

Despite the temporal expansions of specific prevalent MLST-CPS types, we also observed that ∼76% of the prevalent types had already been present in the preceding period (2009 - 2015) with a prevalence exceeding 2% in the majority of world regions where they are currently prevalent (i.e. >5%) (Supplementary Table 5). This observation suggests that CRAB population compositions exhibit local stability over this timeframe. To investigate the temporal stability in more detail, we compared the MLST-CPS type. compositions of specific countries over time. We focused on three countries - China, Germany, and the USA - each of which possessed a sufficient number of sequenced isolates from two distinct periods, namely 2009 - 2015 and 2016 - 2022. In each of these countries, we observed that the previous CRAB-type composition was more similar to its contemporary counterpart than to the contemporary composition of most other countries worldwide, which confirms the temporal stability of CRAB populations in these countries (Fig 3B).

To gain deeper insights into the spatio-temporal dynamics of CRAB types, we proceeded with a phylogeographical analysis of the dominant ST2 clade using established methods.^35^ The analysis covered ∼70% of the available 11,129 *A. baumannii* genomes, providing insights into this pathogen’s transmission dynamics at an unprecedented scale. Specifically, we reconstructed a time-scaled phylogenetic tree using the core genome sequences of 7,720 ST2 isolates collected from across the globe (Fig 3C, Methods). Encouragingly, we found a strong temporal signal within the dataset, affirming the accurate capture of chronological information within the reconstructed time-scaled tree (Fig S7). This was further supported by our estimated root date of 1973 for the tree, which aligns with epidemiological data regarding the emergence of ST2.^36,37^

Then, we used the time-scaled tree to assess the timeframe needed for CRAB lineages to spread across different geographical distances. We observed strong phylogeographic clustering, with genetically closely related isolates primarily circulating within the same country over a short period. Specifically, on average, two contemporary CRAB isolates that diverged very recently (*i.e.* within the past 6 years) exhibited a 19-fold higher likelihood of originating from the same country rather than from different countries (Fig 3D), which is in agreement with our earlier analysis on the stability of the strain compositions of specific countries (Fig 3B). This local spatial clustering was also evident in Europe, which is a region of multiple smaller countries in contrast to other world regions (Fig S8). Nevertheless, over a longer period (6-12 years), this local spatial clustering disappears, indicating substantial transmission of isolates across countries (Fig 3D, Fig S8).

The dominance of local transmissions is also evident on a smaller geographic scale, such as cities. In this analysis, our focus was on Eastern and Southern European regions covered by our CRAB collection due to the availability of fine-scale geographic information (Supplementary Table 1). Specifically, when considering two contemporary CRAB isolates that diverged in the past 2 years, there was a 4-fold higher likelihood that they originated from the same city rather than from different cities within the country (Fig 3E). This pattern indicates that within a 2-year timeframe, locally circulating strains in specific cities are poised to account for the majority of the CRAB infections.

The observation that spatial clustering of isolates only disappears after several years indicates the dominance of local transmissions in the case of this nosocomial pathogen. The calculated value suggests that CRAB strains currently circulating within a specific country are likely to be responsible for the majority of infections within a 6-year period.

### Phylogeny-guided phage hunting and phage sensitivity profiling

We sought to demonstrate the potential of bacterial genomic surveillance in identifying a phage collection that is effective against the CRAB types responsible for the majority of infections within a specific region. To achieve this goal, we focused our efforts on the five neighbouring Eastern and Southern European countries that our genomic sample collection covers: Hungary, Romania, Serbia, Montenegro, and Bosnia and Herzegovina. For phage hunting, we selected eleven MLST-CPS types that are both prevalent in this region, collectively accounting for 90.7% of the 546 samples between 2016 and 2022 (Supplementary Table 1), and also regionally relevant, since ten out of eleven are among the most prevalent MLST-CPS types in entire Eastern Europe (Fig 2D). Subsequently, we conducted our phage hunt using representative clinical isolates from each of the 11 MLST-CPS types (Supplementary Table 13). Phages were isolated from raw communal wastewater collected in five Hungarian cities (Methods). Our efforts resulted in the successful isolation of a total of 15 phages, all of which exhibited potent lytic activity (>10^6^ pfu) against at least one target clinical isolate. Next, we quantified the efficacy of the isolated phages on a larger collection of 92 CRAB clinical isolates from the 11 MLST-CPS types (Supplementary Table 9-10). As anticipated, each MLST-CPS type had a unique phage susceptibility profile (Fig 4A). Remarkably, for eight of these MLST-CPS types, we identified phages that effectively targeted over 95% of the tested clinical isolates (Fig 4A). For example, phage Highwayman potently targeted 28 out of the 29 tested isolates that belong to the most prevalent type (ST2-KL3) in Eastern Europe.

**Figure 4.**
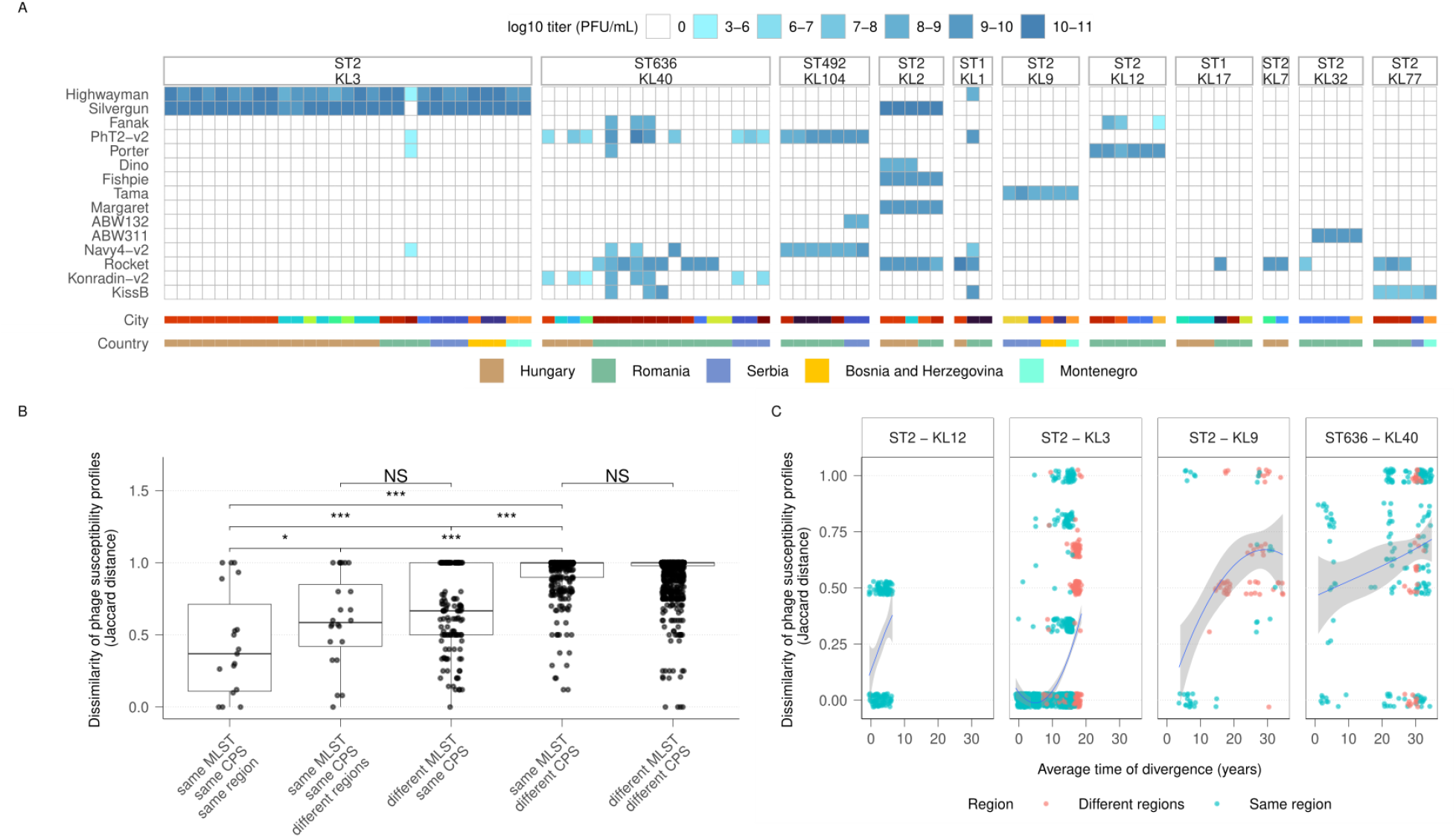
Genomic surveillance-informed phage discovery. **A,** Phage sensitivity profiles of 92 CRAB isolates belonging to the most abundant 11 MLST-CPS types in the studied Eastern and Southern European countries. The assessments were conducted on the 15 isolated phages. The intensity of the blue colour indicates the mean log_10_ phage titer (PFU/ml) of three technical replicates. The geographic origin (country and city) of the CRAB isolates is colour-coded at the bottom of each column. Data is available in Supplementary Table 10. **B,** Pairwise comparisons of binary phage sensitivity profiles of 199 *A. baumannii* isolates. Isolates are classified by MLST, CPS and geographical locations. Jaccard distances of 0 and 1 indicate identical and 100% different profiles, respectively. Each data point represents the average Jaccard distance value of all pairwise comparisons between two isolate categories as specified by MLST, CPS and region categories. Number of pairs from left to right: 23, 28, 150, 340, 1462, one-sided permutation test with 5000 random pair classifications, significance labels are “NS” for not significant, * for *p* < 0.05, and *** for *p* < 0.001). **C,** Pairwise phage sensitivity profile dissimilarities correlate with genetic distances (average time of divergence). *p* and *n* values from the Mantel test (from left to right): 0.008, 0.002, 0.018, 0.009 and 153, 1,431, 91, 231, respectively. The fitted line is smoothed by a loess method. This analysis was done only for those four MLST-CPS types where the phage susceptibility profiles were measured for at least 10 CRAB isolates. Data are available in Supplementary Table 11.

To further evaluate the broader relevance of the discovered phages, we tested them against an additional set of 107 *A. baumannii* clinical isolates using a simplified binary phage susceptibility profiling method (see Methods, Supplementary Table 9-10, Fig S9). These 107 isolates included further clinical isolates of the eleven MLST-CPS types from Eastern and Southern Europe and from distant locations, such as Western Europe and North America (Supplementary Table 9-10, Fig S9), representing an even broader genetic diversity for the eleven MLST-CPS types (Fig S10A-K) and 15 additional ST backgrounds (Supplementary Table 9-10). Subsequently, we conducted a systematic comparison of these binary phage susceptibilities across the entire dataset (i.e. 92+107 *A. baumannii* isolates). The analysis revealed that variations in phage susceptibility profiles are mainly determined by CPS type, but altered ST background and large geographic distance also played a significant, albeit weaker, role. (Fig 4B). The observed impact of geographical distance hints that even within an MLST-CPS type, genetic divergence may also shape phage susceptibility. Indeed, pairs of isolates that diverged earlier tend to show less similar phage susceptibility profiles (Fig 4C). Nevertheless, individual phages differ in their resilience to genetic variations within MLST-CPS types, with several phages retaining a killing effect even across isolates of the same MLST-CPS type that diverged genetically a long time ago. For example, Silvergun is effective against all 23 tested ST2-KL2 isolates, despite their 30+ year genome divergence (Fig S11). The ability of these phages to remain effective over larger genetic distances makes them promising candidates for therapeutic applications.

Finally, specific CRAB types, such as ST636-KL40, exhibit genetic variability that is not well captured by the combined MLST-CPS typing or by the genetic distance alone, but it still markedly influences phage susceptibility (Fig 4A, Fig 4C). We hypothesised that this genetic variation could induce changes in the cell surface without affecting the predicted CPS type.^33^ Therefore, we characterised the surface properties of recently diverged isolates of the ST636-KL40 type using an infrared spectroscopic method (Methods, Fig S12). Importantly, we detected cell surface variations that correlate significantly with phage susceptibility profile differences among the isolates (Fig S12).

In summary, we have successfully constructed a phage library that is highly effective against most clinical isolates of CRAB in Eastern Europe. While bacterial susceptibility to phages is primarily dictated by CPS type, additional genetic variations that arise during the pathogen’s spread across geographical regions also play a role. This underscores the importance of tailoring phage collections to the local genetic diversity of the pathogen.

### Old lytic phages with new targets

As a next step, we sequenced the genomes of the 15 isolated phages. We aimed to determine their suitability for clinical use by defining established safety criteria, including a functionally well-annotated genome, an obligate lytic life cycle, and the absence of harmful genes.^39, 40^ First, we analysed their protein sequence-based relationship with 3,562 known species of tailed dsDNA phages (Caudoviricetes) as done before.^41,42^ This allowed us to define their higher-order taxonomy and genetic relatedness to phages that had been previously characterised as lytic or had been utilised for therapeutic purposes (Supplementary Table 12).

Out of the 15 phages we discovered, which belong to 5 different taxonomic groups, 14 exhibit extensive sequence similarity to previously used therapeutic phages or well-characterised lytic phages (Fig 5, Supplementary Table 12). For example, seven phages from the *Autographoviridae* family share a DNA sequence identity of 80-84% with the therapeutic phage AbTP3phi1 (Fig 5E). Furthermore, two-two myophage morphotype phages in the *Twarogvirinae* subfamily show 87-95% DNA sequence identity with two other previously used therapeutic phages, AB-Navy-4 and AB-Navy-71, respectively (Fig 5E). In all of these cases, despite the high sequence similarities stemming from conserved core gene sets, notable differences exist in critical genetic determinants of host specificity. Specifically, depolymerase-carrying tail proteins responsible for CPS decomposition differ from the corresponding components of previously reported treatment or lytic phages in almost every case (Fig S13, Supplementary Table 17). These findings suggest the existence of closely related Acinetobacter phages across the world, with minor genetic adaptations to accommodate various host strains. The relatively modest genetic variations compared to previously established lytic and therapeutic phages simplify the comprehensive characterization necessary for clinical application.

**Figure 5.**
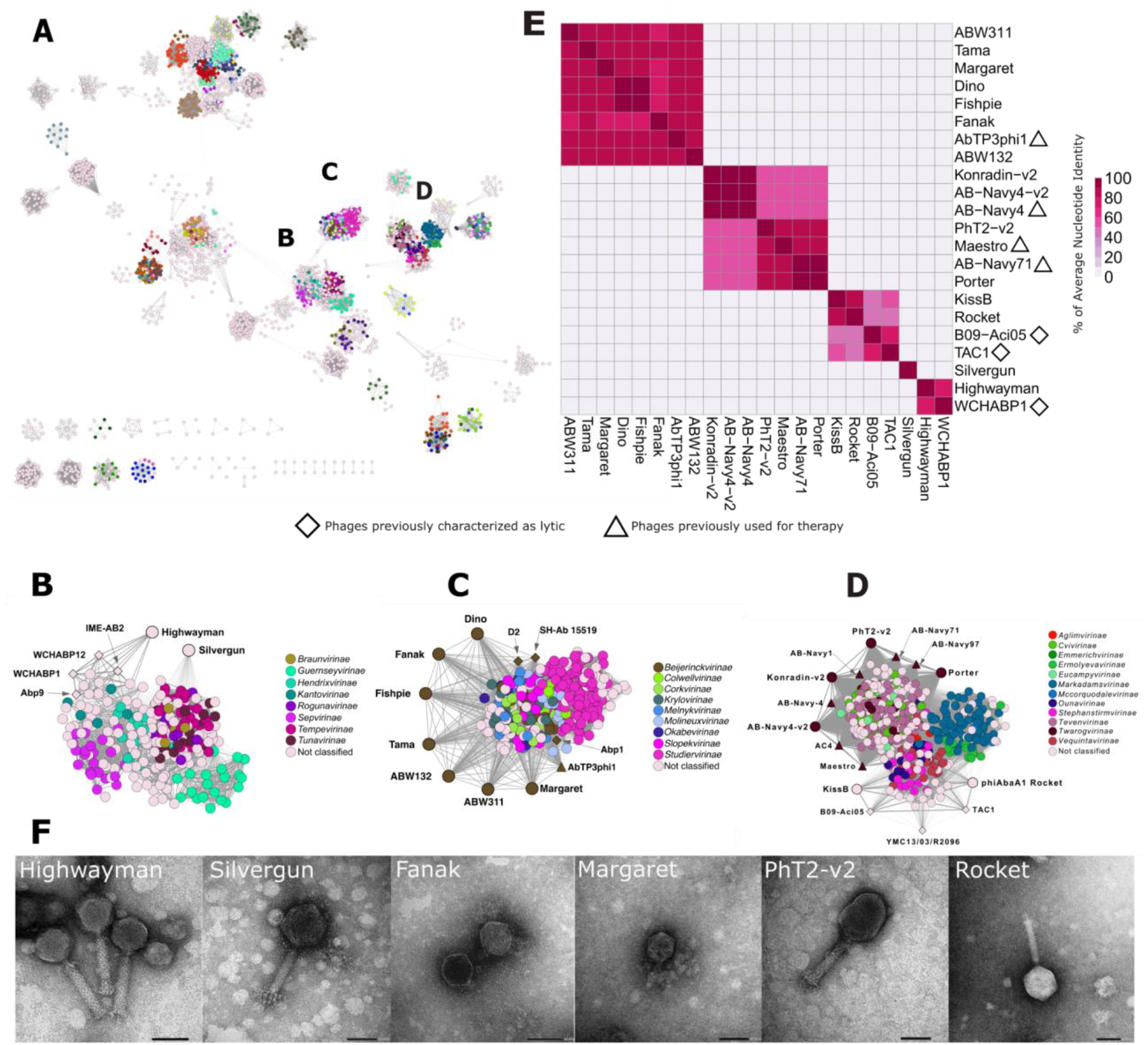
Sequence-based relationship of the 15 discovered phages and known phage species. **A,** Protein orthogroup-based relationship network of 3562 *Caudoviricetes* phages representing all species in the ICTV taxonomy, plus the 15 discovered phages generated using vContact2 (Methods). Nodes are coloured based on their Subfamily membership. Edge thickness is proportional to the number of shared orthogroups between two phages. The three circled clusters are enlarged in figure panel B-D. Data is available in Supplementary Tables 13 and 15. **B-D,** Enlarged protein orthogroup-based clusters containing the 15 phages with their most related phages. The closest previously characterised lytic or therapeutic phages are represented as rhombus or triangle, respectively. Nodes are colour-coded based on their Subfamily membership. Subfamily names are listed. Nodes representing the 15 phages discovered here are represented with enlarged nodes. **E,** Pairwise Average Nucleotide Identities for the 15 discovered phages and for their closest phage relatives that have been used previously for therapy or characterised as lytic. Data is available in Supplementary Tables 12, 14, 15, and 16. **F,** Electron microscopic pictures of two phages from each subnetwork out of the 15 discovered phages. Detailed morphological and genomic analysis are available for each 15 discovered phages in Supplementary Note. Scale bar = 50 nm.

In contrast to the above general patterns, Silvergun exhibits a unique genetic composition that distinguishes it from all other currently known phages. Silvergun shares very few orthogroups with any other known phages, suggesting that Silvergun may be the first characterised member of a new bacteriophage family (Fig 5A-B, Fig S14, Supplementary Table 18). Importantly, Silvergun is able to target the two most prevalent globally spreading strains belonging to ST2-KL3 and ST2-KL2 group (Fig 4A), which collectively account for 31% of the *A. baumannii* genomes isolated in 2022 (Fig 3A). It is possible that the presence of two predicted depolymerase regions in the tail spike and fibre proteins of this phage explains its uniquely relevant host specificity (Supplementary information).

It is important to highlight that none of the 15 isolated phage genomes were found to contain toxin-, antibiotic resistance-genes or temperate markers that would hinder their potential future therapeutic application (Supplementary Table13). All studied phages with similarity to well-characterised lytic phages are predicted to be lytic using the PhageAI tool, with the exception of Highwayman whose lifestyle could not be defined reliably (Methods, Supplementary Table 13). Therefore, we experimentally confirmed the absence of lysogenic behaviour for Highwayman, along with Silvergun and PhT2-v2 phages, on their corresponding hosts using a previously established method (Fig S15.1-3).^43^ Further details on the genomic and morphological analysis of each phage are included in Supplementary Note S1.

### Precision phage cocktail against the ST2-KL3 strain type in Europe

The first clinical case study involving *Acinetobacter* revealed a common challenge of phage therapy: the emergence of bacteriophage resistance, which can lead to treatment failure.^7,44^ To address this issue, a common strategy is the application of phage cocktails (rather than individual phages) that can mitigate the emergence of phage resistance.^45^ Therefore, our goal was to develop a phage cocktail against the European ST2-KL3 CRAB lineage (Fig 3C). We targeted it for two reasons: widespread prevalence across the continent (Fig 2D) and well-represented phylogenetic diversity among our isolates (Fig S10D), aiding the development of a widely applicable phage cocktail.

To start, we focused on Highwayman (H) and Silvergun (S), two phages that together target 98% of a representative set of 56 ST2-KL3 clinical isolates (Fig S9). By co-incubating *in vitro* these two phages and their combination (termed HS) with the ST2-KL3 clinical isolates, we observed the emergence of phage resistance typically after 10-12 hours, irrespective of the application of these phages individually or in combination (Fig 6A, Fig S16). Next, we examined whether Highwayman or Silvergun resistance induces sensitivity to other phages in the collection. We found that all 22 tested Silvergun resistant-lines became sensitive to at least one of four other phages from the collection (Fig S17, Supplementary Table 20). Combining one or two of these four phages with the initial cocktail of Highwayman and Silvergun significantly suppressed the growth of the target cells when measured up to 24 hours, indicating diminished resistance (Fig 6B, Fig S16). Even when resistance was assessed up to 72 hours against one of the best performing four-phage cocktail, termed HSFPh (Highwayman, Silvergun, Fanak and PhT2-v2), we detected only sporadic instances of bacterial resistance (Fig 6C, Supplementary Table 19).

**Figure 6.**
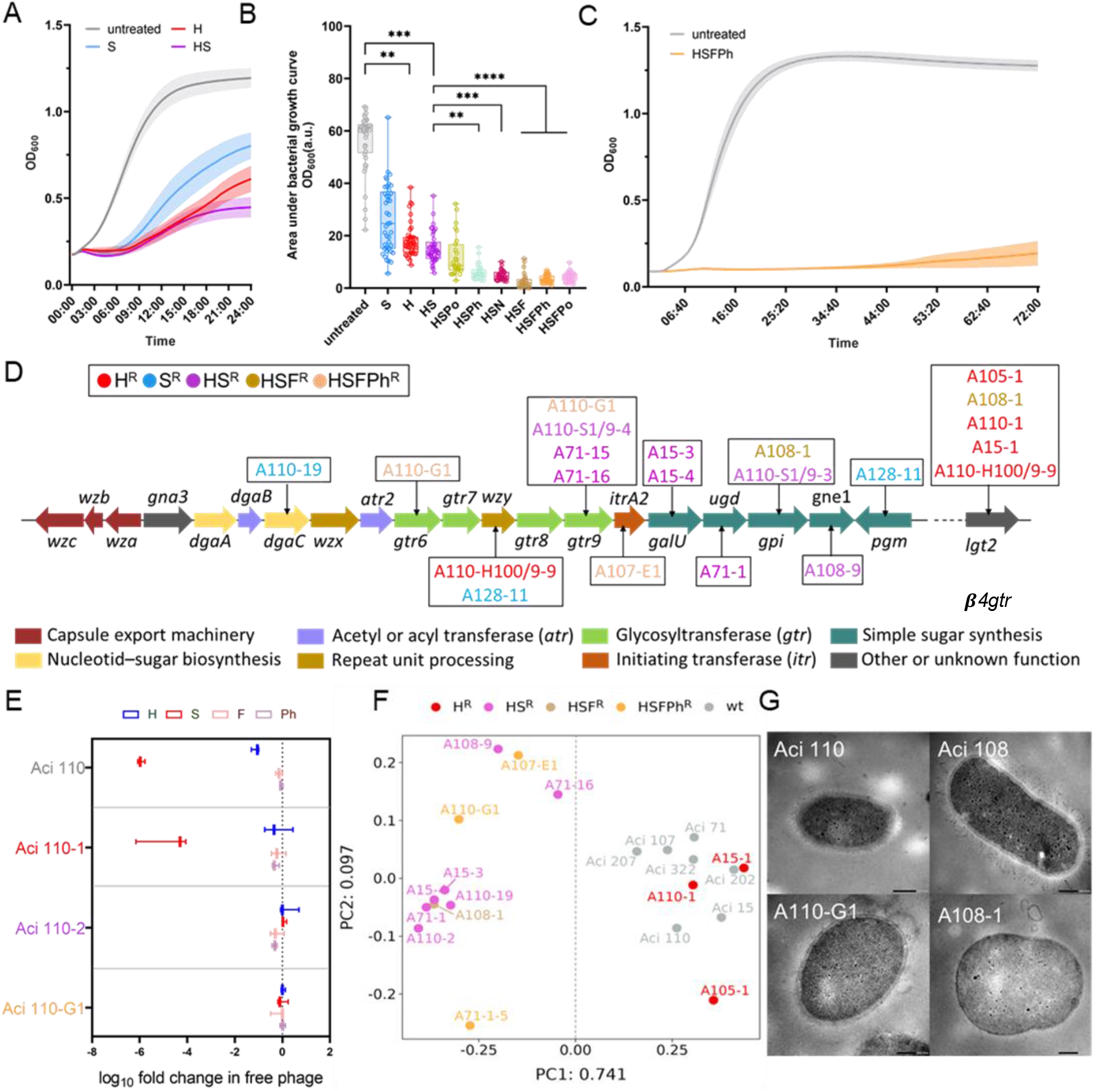
Constructing a precision phage cocktail against the dominant ST2-KL3 strain in Europe. **A**, Growth curves (OD_600_) measured for 24 hours for 41 randomly selected ST2-KL3 European isolates either in the absence (untreated) or in the presence of the Highwayman (H) and Silvergun (S) phages alone, or in combination (HS). Plotted values represent the mean ± CI 95%. **B**, Area under the bacterial growth curves either in the absence (untreated) or in the presence of different phages alone or in combinations. Data points represent the mean area under the bacterial growth curve value of three technical replicates measured with each ST2-KL3 CRAB clinical isolates (*n* = 41 distinct isolates, except for the 3-phage combinations, where *n* = 27). A.u stands for arbitrary unit. ** *p* ⋜ 0.05, *** *p* = 0.0001, **** *p* < 0.0001 from two-sided Kruskal-Wallis test. Abbreviations: H – Highwayman, S – Silvergun, F – Fanak, Po – Porter, N - Navy-v2, Ph – PhT2-v2. **C,** Growth curves (OD_600_) were measured for 72 hours for 25 randomly selected ST2-KL3 isolates in the absence (untreated) or in the presence of the phage cocktail HSFPh. Plotted values represent the mean ± CI 95%. Data for panels A, B, and C are available in Supplementary Table 19. **D**, Schematic representation of the KL3 CPS biosynthetic gene cluster. Functional categories of the gene products are colour-coded. The vertical arrows point to genes which harbour a loss-of-function deletion/insertion or a point mutation in different phage-resistant isolates listed within the frames. Text colours within the frames represent phage resistance categories as follows: red, blue, purple, brown, and orange colours corresponding to H, S, HS, HSF, and HSFPh resistance, respectively. Note that the *lgt2* gene is encoded trans of the KL3 CPS gene cluster. **E**, Phage adsorption assay with the phage H, S, F, and Ph for a wild-type ST2-KL3 isolate (Aci 110) and for its three phage-resistant derivatives: Aci 110-1, Aci 110-2, and Aci 110-G1 which are resistant to H, HS, and HSFPh, respectively. Results show the log_10_ reduction in free phage titres at a maximum adsorption time point in comparison to the t_0_ time point (dashed line) after mixing phages and host bacteria (for details see Methods and Supplementary Table 22). Data are mean ± standard deviation, *n* = 3 technical replicates. **F,** Principle coordinate analysis plot derived from Fourier-transformed infrared measurements (see Methods, Supplementary Table 23) differentiates phage-resistant ST2-KL3 isolates that harbour loss-of-function mutations within the CPS biosynthetic pathway from those that do not (*i.e.* wt and resistant lines harbouring mutation exclusively in the *lgt2* gene). Dashed line separates the two groups. Colour codes represent phage resistance categories as follows: red, purple, brown, and orange corresponding to H, HS, HSF, and HSFPh resistance, respectively. Grey colour represents the wild type. **G**, TEM pictures showing HSFPh-(A110-G1) and HSF-resistant (A108-1) lines that harbour loss-of-function mutations in the CPS biosynthetic genes have altered capsule compared to their wild-type counterparts (Aci-110 and Aci-108, respectively). Scale bar = 200 nm.

To elucidate the genetic basis of the emerging bacterial resistance, we sequenced the genomes of 19 phage-resistant bacterial lines, including lines that emerged against Highwayman, Silvergun, or their corresponding two-phage (HS) or four-phage (HSFPh) cocktails (Supplementary Table 21). Cell lines resistant exclusively to Highwayman contained loss-of-function mutations in the ***β***-4-glucosyltransferase gene (***β****4gtr,* WP_001033817.1). In contrast, mutants resistant to Silvergun or the two other phages in the cocktail (i.e. Fanak and PhT2-v2) typically contained loss-of-function mutations in the genes involved in the capsular polysaccharide (CPS) biosynthesis pathway (Fig 6D, Supplementary Table 21). These patterns suggest that, in contrast to Highwayman, resistance to these three phages requires defective capsule production. This notion is confirmed by the abolished phage adsorptions to the cell surface (Fig 6E) and altered cell surface properties revealed by the combination of infrared spectroscopy and electron microscopy (Methods, Fig 6F-G). Overall, as decapsulation is a known phage-resistance mechanism among other MLST-CPS strain types of CRAB,^34,46^ (ref) these results demonstrate that ST2-KL3 shares this feature with other CRAB types. Notably, however, the sensitivity of the Silvergun resistant lines to Fanak and PhT2-v2 indicate that the decapsulated cell surface is different upon Silvergun resistance as in the case of the HSFPh cocktail resistance, which notion is supported by the difference in the depleted genes (Fig 6D).

Given the capsule’s crucial role in bacterial defence, we hypothesised that the emergence of resistance to the phage cocktails may render the bacterium less virulent or more susceptible to antibiotics.^34,47,48^ To test this, we first measured the *in vivo* virulence of the cocktail-resistant isolates in *Galleria mellonella* (*G. mellonella*) larvae infection model.^49,50^ As expected, CPS-deficient strains exhibited significantly decreased virulence compared to the ancestor wild-type strain (Fig 7A, Fig S18A-E). Importantly, such a decrease in virulence was not observed in the case of Highwayman-resistant lines, which have intact CPS (Fig 7A, Fig S18F). Next, we measured the antibiotic susceptibility profiles of 34 phage-resistant lines against a panel of clinically relevant antibiotics (Fig S19). Remarkably, 71% of the colistin-resistant lines transitioned from resistance to sensitivity against colistin. Moreover, we identified two HSFPh-resistant lines, both of which regained sensitivity to meropenem, shifting from a resistant state to an intermediate state (Fig 7B, see Methods).

**Figure 7.**
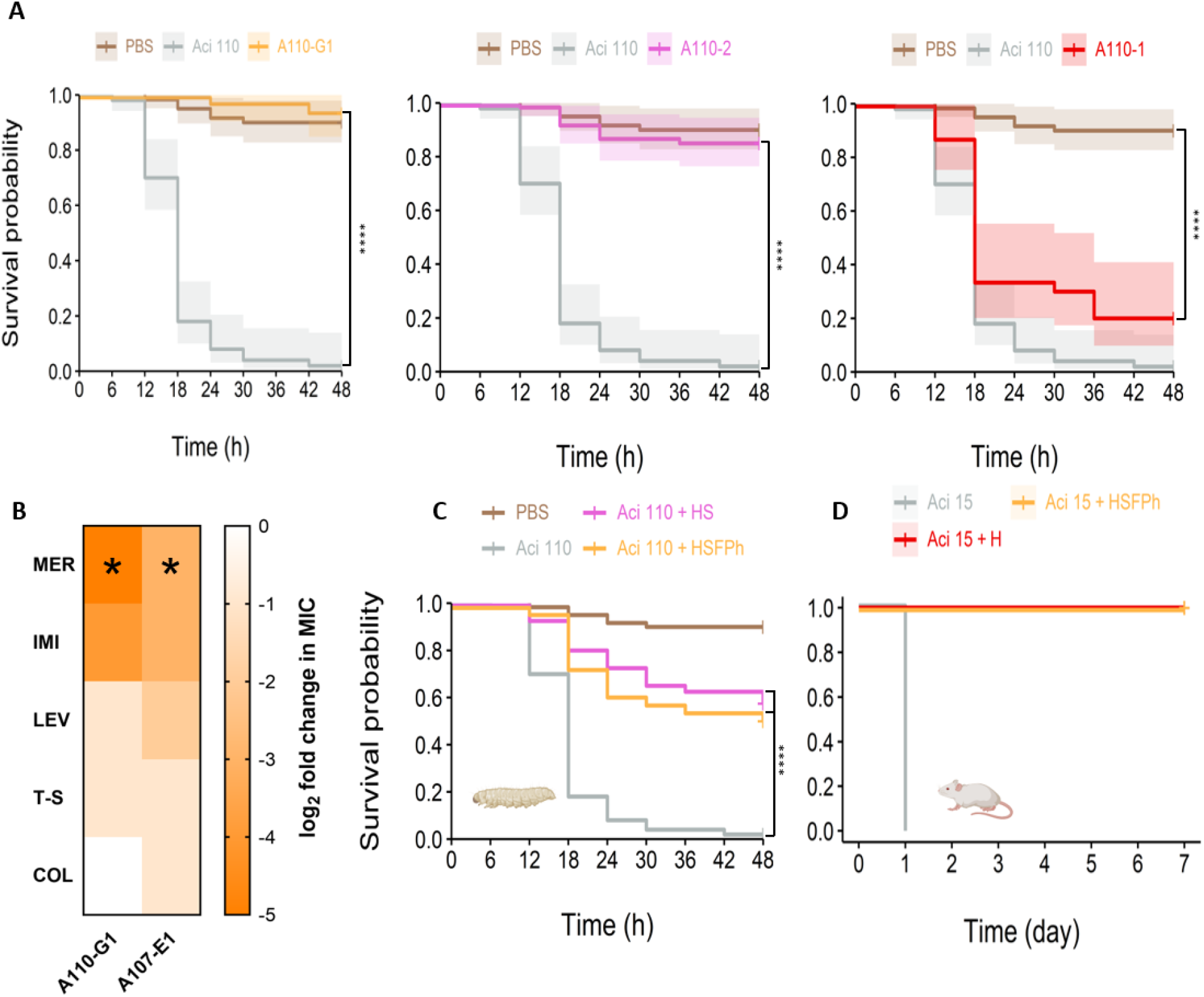
Cost of phage-resistance and effectiveness of the cocktail in *in vivo* infection models. **A,** Kaplan-Meier curves showing the survival of *G. mellonella* larvae after infection with either phage-resistant ST2-KL3 CRAB lines or with wild-type counterparts. HSFPh- and HS-resistant lines (left and middle panel) have decreased virulence compared to their wild-type counterparts, in contrast to the H-resistant line (right panel). **** indicates *p* < 0.0001 from two-sided Log-rank test, *n* = 10 larvae/group, ≥ 3 biological replicates/group, PBS means larvae injected only with PBS, inoculum size = 9*10^6^ CFU. **B,** Antibiotic sensitivity profile of the HSFPh cocktail-resistant lines. The intensity of the orange shading indicates the median (n≥3) log_2_ fold change in the minimum inhibitory concentration (MIC) of the cocktail-resistant lines in comparison to their wild-type counterparts as measured using the microbroth dilution method (see Methods and Supplementary Table 25). The darker the intensity, the higher the reduction in the MIC. The 5 antibiotics with clinical relevance are: MER-meropenem, IMI-Imipenem, COL-colistin, LEV-levofloxacin, T-S-a combination of Trimethoprim and Sulfamethoxazole (1:5 ratio). The star indicates a transition from the resistant state to the intermediate state according to EUCAST clinical breakpoint. **C,** Kaplan-Meier curves showing that both the cocktail HS (MOI 10:10, purple) and the cocktail HSFPh (MOI 10:1:0.1:0.1, orange) significantly improve the survival of *G. mellonella* larvae infected with Aci 110 ST2-KL3 isolate as compared to the untreated larvae (grey) (CFU = 9*10^6^, *n* = 10 larvae/group, ≥ 3 biological replicates/group, PBS – larvae injected only with PBS). **** indicates *p* < 0.0001 from a two-sided Log-rank test. **D,** Kaplan-Meier curves showing that both Highwayman phage alone (MOI 6, red) and the cocktail HSFPh (MOI 6:1:1:1, orange) saved 100% of mice infected with Aci 15 ST2-KL3 isolate as opposed to the untreated animals (grey) (CFU = 10^9^, *n* = 5 mice/group) *** indicates *p* < 0.00027 from the Log-rank test. For A, C, and D data are available in Supplementary Table 24.

Finally, we tested the efficacy of the phages in *in vivo* infection models. First, *G. mellonella* larvae infected with one of the ST2-KL3 isolates were treated with the phage Highwayman or with its cocktails HS and HSFPh 5 min post-infection resulting in a significant improvement in the survival rate of the larvae in all the cases (Fig 7C). The efficiency of the phages in *G. mellonella* was corroborated by an infection model in mice (see Methods). Specifically, after injecting one of the ST2-KL3 CRAB isolates intraperitoneally into mice, we observed that both treatment with Highwayman phage alone and with the HSFPh cocktail resulted in the survival of 100% of the animals, whereas all untreated animals died 1-day post-infection (Fig 7D).

In sum, resistance to HSFPh phage cocktail is relatively rare, and even when it does arise, it leads to decreased virulence and enhanced meropenem sensitivity. Notably, as carbapenem resistance is associated with high mortality in patients infected with Acinetobacter,^51^ (ref) this resensitization by phage resistance can be clinically relevant and supports the use of the cocktail in combination with meropenem.^52,53^

## Discussion

To combat the escalating crisis of antibiotic resistance, especially in the context of hospital-acquired multi-drug resistant infections, this study explores the promise of precision phage therapy guided by genomic surveillance. Despite successful case studies of phage therapy,^7,8,9,10,11,12^ the limited range of bacterial strains targeted by individual phages hinders its widespread application. We aim to overcome this limitation by demonstrating how genomic surveillance can guide phage therapy for carbapenem-resistant A*. baumannii* (CRAB), offering a scalable and cost-effective strategy. Instead of engaging in the time-consuming process of isolating individual phages for each patient’s bacterial strain,^7^ we proactively employ genomic surveillance to assemble region-specific phage collections that target the majority of CRAB infections within a particular geographical area.

Our study introduces several significant novelties. First, we conducted a global analysis of over ten thousand CRAB genomes, identifying 29 prevalent CRAB types responsible for the majority of infections worldwide. Despite inter-regional variation, these prevalent types exhibit homogeneity within specific world regions, suggesting the potential effectiveness of region-specific phage collections for CRAB targeting. Second, our phylogeographical analysis reveals the temporal stability of CRAB populations within countries over a six-year period, providing a crucial time frame for these preemptively prepared region-specific phage collections. Third, in Eastern Europe, a region with high CRAB infection rates, we performed a phylogeny-guided phage hunt, matching the 11 most prevalent CRAB types with 15 new phages with therapeutic potential. Additionally, we systematically characterised the host range of these phages using 199 clinical isolates, allowing us to assess the geographic scope of their applicability. Finally, as a proof-of-concept, we formulated a precision phage cocktail that effectively inhibits a broad range of clinical isolates of the globally most prevalent CRAB type ST2-KL3, demonstrating minimal emergence of resistant variants and efficacy in *in vivo* animal infection models.

Overall, this study pioneers the application of bacterial phylogeography to scale up phage therapy. The application of genomic surveillance has proven pivotal in elucidating the evolutionary history and transmission dynamics of pathogens^24^ and is emerging as a tool to inform vaccine design.^54,55^ The proposed phage therapy framework aims to design phage cocktails that provide benefits to the highest number of patients across geographical scales. Importantly, this will facilitate study recruitment, crucially speeding-up clinical trials and leading to clinical validation. Moreover, a scale-up will lead to enhanced cost-effectiveness for personalised phage therapy.^56^ These milestones are significant for advancing the field and ensuring the precision and practicality of phage therapy in clinical settings. Furthermore, by creating targeted collections of off-the-shelf medicinal phage products against nosocomial pathogens, the study establishes phage therapy as a viable adjunctive treatment for managing acute infections that require immediate intervention. In a similar vein, a systematic approach was recently taken for the development of a four-phage cocktail to selectively kill *Escherichia coli*, an agent of fatal infections in haematological cancer patients.^19^ Although the phage cocktail was designed to cover a substantial fraction of phylogenetically diverse *Escherichia coli* strains sourced from hospitalised patients across 28+ countries, its efficacy might be limited within any given geographical region due to the absence of consideration of the spatio-temporal distribution of strains. Importantly, thanks to the rapidly increasing number of available genome sequences, the genomic surveillance-based framework developed in this study is not limited to CRAB and can be readily applied to target other nosocomial pathogens. This broadens the potential impact of the study, making its findings applicable beyond CRAB and contributing to the development of effective strategies for combating a range of infectious diseases in healthcare settings.

### Limitation of the study

We acknowledge several limitations in our study. Firstly, while we made efforts to control for sampling bias in the analysis of over 10,000 genome sequences, variations in sequencing efforts across different countries may still introduce some bias. Secondly, due to constraints in the availability of worldwide clinical CRAB samples, most laboratory tests, including phage typing, were conducted on CRAB isolates coming from only four world regions. Most importantly, ST2-KL3 CRAB samples were sourced exclusively from three world regions (East-, West, South-Europe), limiting our knowledge on how widely our four-phage cocktail developed for this prevalent CRAB type could be applied. Thirdly, our study faced limitations in demonstrating the superiority of the four-phage HSFPh cocktail over the single phage Highwayman, particularly in overcoming phage resistance, as the animal infection models exhibited fast mortality of the infected animals.

## Acknowledgement

This work was supported by the National Laboratory of Biotechnology Grant 2022-2.1.1-NL-2022-00008 (C.P., B.K., B.P.), the National Laboratory for Health Security Grant RRF-2.3.1-21-2022-00006 (B.P.), the European Union’s Horizon 2020 research and innovation programme under grant agreement no. 739593 (B.K., B.P.), National Research Development and Innovation Office ‘Élvonal’ Programme KKP 126506 (C.P.) and KKP 129814 (B.P.), the National Research, Development and Innovation Office grant FK-135245 (B.K.), FK-124254 (O.M.), PD-131839 (E.A.), János Bolyai Research Fellowship from the Hungarian Academy of Sciences BO/352/20 (B.K.), BO/00303/19/8 (O.M.), New National Excellence Program of the Ministry of Human Capacities (UNKP-20-5-SZTE-654 and UNKP-21-5-SZTE-579) (B.K.), Proof of Concept grant of the Eötvös Loránd Research Network (ELKH-PoC-2022-034) (B.K.), Ministry of Education, Science, and Technological Development of the Republic of Serbia, Serbia [grant No. 451-03-68/2022-14/200110]. We thank Gordana Mijovic and Aleksandra Smitran for providing clinical *A. baumannii* isolates from Montenegro and Bosnia&Herzegovina, Petra Szili for providing help with the larvae experiments and those who provided wastewater samples for phage isolation: Alföldvíz Zrt., Budapest Sewage Works Ltd., Debreceni Vízmű Zrt., Szegedi Vízmű Zrt.

## Author Contributions

Conceptualization and Methodology, B.K., B.P., M.K., T.S., H.H.M., and O.M.; Investigation and validation, M.K., T.S., H.H.M., B.E., O.M., G.A., L.D., T.F.P., G.S., I.G and S.A.Z.; Software, Formal analysis and data curation, T.S., O.M., M.K., A.A., G.A., B.M.V., E.A., M.S., G.F., K.N.V.; Visualisation, T.S., M.K., O.M., H.H.M., A.A., B.B., E.A., G.A., and M.S.; Writing – Original Draft, B.K., B.P., M.K., T.S., H.H.M., O.M., G.A., A.A.; Writing – Review & Editing, B.K., B.P., C.P.; Resources, C.B., C.C., M.I., O.I., B.L., S.M., L.O., E.S., U.O.S.G., A.T., C.V.d.H., A.B., A.V., K.K and M.S.L.; Supervision, B.K and B.P.

## Data Availability

Assembled genomes for this study have been deposited in the European Nucleotide Archive (ENA) at EMBL-EBI under accession number PRJEB64245 (https://www.ebi.ac.uk/ena/browser/view/PRJEB64245). Source data are provided with this paper.

## Code Availability

Scripts and other files needed to reproduce the analysis are available at https://github.com/stitam/Koncz-et-al-Genomic-surveillance. Most analytical pipelines were orchestrated using Nextflow v22.10.1.^57^ Steps in these pipelines were containerised using Docker v24.0.7^58^ and Singularity v3.9.0.^59^

## Methods

### Ethical statement

This research complies with all relevant ethical regulations and was approved by the Scientific and Research Ethics Committee of the Hungarian Health Science Council (BMEÜ/271-3/2022/EKU). Animal experiments were approved by the Animal Experimentation Scientific Ethical Council: Investigation of the Effectiveness of Bacteriophage Strains with Therapeutic Potential in Animal Experimental Model Systems with an authorization number: KA-3429.

### Establishing a collection of *Acinetobacter baumannii* isolates

Due to the underrepresentation of genomic samples from the Eastern and Southern European region, we collected and sequenced 419 Carbapenem-resistant *Acinetobacter baumannii* CRAB clinical isolates from 5 countries from these regions (Hungary (n=253), Romania (n=120), Serbia (n=28), Montenegro (n=9), and Bosnia and Herzegovina (n=9)) between 2011 and 2022 from 44 healthcare facilities in 34 cities. These isolates were categorised as CRAB strains after performing the standard antimicrobial susceptibility tests according to the methods of the European Committee on Antimicrobial Susceptibility Testing (EUCAST). Nine antimicrobial agents were tested regularly including amikacin, gentamycin, tobramycin, ciprofloxacin, levofloxacin, imipenem, meropenem, trimethoprim-sulfamethoxazole, and colistin. The minimum inhibitory concentrations (MICs) were determined by *E*-test for carbapenems and by broth microdilution in the case of colistin. The susceptibility to the remaining antibiotics was determined by the disc diffusion method. The susceptibility test results were interpreted based on the EUCAST breakpoints^1^ (Supplementary Table 26). CRAB isolates were defined with both imipenem (MIC > 4 mg/L) and meropenem resistance (MIC > 8 mg/L). To study the host range of the isolated bacteriophages on an extended set of CRAB isolates, 16 additional CRAB isolates were obtained from BEI Resources (https://www.niaid.nih.gov/research/bei-resources-repository) and 47 from the Belgian Acinetobase collection.^2^ The identity of these strains is available in Supplementary Table 9.

### Genome sequencing

For the isolation of the genomic DNA, all *Acinetobacter* strains were routinely cultured on LB broth (LB, 5 g of tryptone, 2.5 g of yeast extract, 5 g of NaCl, and 500 ml of distilled water) or LB agar (LB plus 1.5% Bacto agar, w/v). For all plaque assays, a 0.5% LB agar overlay (LB and 0.5% Bacto agar w/v) was inoculated with 0.1 ml of a fresh overnight LB culture of the host and poured over LB agar plates.

All strains were grown at 37°C. Genomic DNA of 410 *A. baumannii* isolates was extracted using the GenElute™ Bacterial Genomic DNA Kit (Sigma-Aldrich) and sequenced by Illumina sequencing.

Sequencing libraries were prepared by using either the MiSeq or the NexteraXT library preparation kit from Illumina according to the manufacturer protocol.

When using the MiSeq library preparation kit, pooled sequencing libraries were denatured with 0.1 M NaOH, diluted to 12 pM with HT1 hybridization buffer (Illumina) and mixed with 40% PhiX Control v3 (Illumina) sequencing control library. Denatured sequencing pools were loaded onto MiSeq Reagent kit V2-300 (Illumina) and 2 × 70 bp sequence reads were generated with an Illumina MiSeq instrument with paired-end sequencing and index 1 sequencing primers spiked in the appropriate cartridge positions (12, 14, and 13, respectively) at a final concentration of 0.5 µM.

When using the NexteraXT library preparation kit, genomic DNA has been fragmented to approx. 300 nt fragments and Illumina sequencing adaptors have been added by using tagmentation. Then sample-specific indexes have been added to each sample by PCR. Pooled libraries have been sequenced by Illumina NextSeq 500 using 2×150 PE sequencing chemistry in multiple sequencing runs.

Sequencing ready libraries were quality control checked by BioAnalyzer2100 instrument using High Sensitivity DNA Chip (Agilent Technologies USA, Cat. No. 5067-4626). Sequencing was carried out on NextSeq 500 sequencing system with NextSeq 500/550 Mid Output Kit v2.5 (300 Cycles) chemistry (Illumina, Inc. USA, Cat. No. 20024905).

### Bioinformatic analysis of the *A. baumannii* genomes

#### Genomes from public databases

We downloaded all publicly available 15,410 *A. baumannii* genomes from the NCBI database as of 09/2022.

#### Genome assembly

The raw reads were trimmed using the cutadapt 4.3 program^3^ (min quality 33, max N 0.5, min read length 30). An optimal reference genome for the reference-guided *de novo* assembly was chosen from the 5,271 NCBI genomes that were available on 19/04/2021. The reference genome was selected by comparing *k*-mer counts using the kmc 3.2.1 software.^4,5,6^ We determined the number of shared *k*-mers between the raw reads and the 5,271 available genomes and selected the one with the highest shared k-mer value as a reference genome for the reference-guided *de novo* assembly.

The assemblies were performed with the SPAdes genome assembler 3.15.5^7^ selecting only those contigs with lengths of at least 1,000 and coverage of at least 10x. Assembly fasta files that were at least an order of magnitude larger or smaller than the median file size were dropped from further analysis.

#### *In silico* molecular typing

Whole genome assemblies were subsequently used for molecular typing. Multi Locus Sequence Typing (MLST) was performed according to the Pasteur scheme^8^, using the mlst v2.23.0 software^9^ which incorporates components of the PubMLST database.^10^ In addition, Kaptive v2.0.3 software^11^ was used for identifying polysaccharide capsule and outer lipopolysaccharide loci and for classifying assemblies into CPS and O-types, using the K and OC locus primary reference databases within the software, respectively.

#### Detection of antibiotic resistance determinants

We detected antibiotic resistance determinants in all *A. baumannii* whole-genome sequences as follows. First, we predicted open reading frames (ORFs) using the Prodigal v2.6.3 software^12^ and then identified resistance genes based on sequence similarity to literature-curated resistance genes compiled in the ResFinder database v2.0.0^13^. Specifically, we performed protein sequence similarity searches with DIAMOND v2.0.15^14^ using 10^-5^ *E*-value threshold, 80% identity and coverage thresholds, and keeping the best hit for each ORF. Beta-lactamase genes were assigned to gene families using two relevant publications.^15,16^

Resistance determinants that were outside the scope of ResFinder were examined separately. These included the presence of the insertion sequence ISAba1, which can be required by some genes to confer resistance against carbapenems,^17,18,19,20^ The absence of the carO gene which encodes a porin channel,^21,17^ and three point mutations indicative of fluoroquinolone resistance, gyrAS81L, gyrBA414T, and parCS84L. For ISAba1,^22,23^ we compiled a list of ISAba1 sequences using the European Nucleotide Archive^24^ and a number of publications.^24,25^ Then, we manually filtered this list selecting only the relevant sequences based on their descriptions. Most genomes contained both the insertion sequence and one or more genes or gene fragments, but LC136852 and LC136853 seemed to only contain ISAba1. LC136852 was used to trim the rest of the sequences and acquire variations for the sequence of ISAba1. Finally, a list of 53 non-redundant ISAba1 sequences was used to search ISAba1 in whole genomes using NCBI BLAST v.2.13.0^26^ with a 10^-5^ E-value threshold. For the rest of the resistance determinants, we assembled a list of sequences including carO,^21^ gyrA, gyrB, and parC and searched for these using NCBI BLAST v.2.13.0 with 10^-5^ E-value threshold. For point mutations, we then used a custom script written in R^27^ and R packages Biostrings v2.66.0,^28^ dplyr v1.1.3,^29^ and seqinr v4.2-23,^30^ for identifying genomes which carry the required amino acid substitutions.

#### Identifying CRAB isolates

For the purposes of this study, an *A. baumannii* isolate is considered CRAB if its genome contains resistance determinants against carbapenems as well as against aminoglycosides and fluoroquinolones. Our goal with this strict definition was to identify high-risk isolates that may be resistant towards a wide range of antibiotics in addition to carbapenems.

Aminoglycoside resistance was determined directly from ResFinder results. If at least one aminoglycoside resistance gene was found, the isolate was considered resistant. While ResFinder contains beta-lactam resistance genes, it does not distinguish between carbapenems and non-carbapenem beta-lactams, therefore a custom approach was applied to predict carbapenem resistance. Specifically, we downloaded all available *A. baumannii* antibiotic resistance test results from the Bacterial and Viral Bioinformatics Resource Center^31^ (accessed 2023-02-16) and checked whether any of the beta-lactam resistance genes, ISAba1, or carO could be associated with observed carbapenem resistance. We found 1,283 observations where test results were available for either imipenem, meropenem, ertapenem, or doripenem, and the results could be linked to genomic sequences in our data set. We then tested for statistical association between the experimentally determined phenotypes and the resistance determinants using contingency tables and standard statistics (Supplementary Table 27). An earlier study suggested that beta-lactam resistance genes bla_OXA-23-like_, bla_OXA-24/40-like_, bla_OXA-58-like_, bla_OXA-143-like_, bla_OXA-235-like_, any bla_NDM,_ bla_VIM_, bla_IMP_, bla_KPC_, bla_GES_ were all associated with carbapenem resistance.^17^ Our analysis was consistent with these findings, therefore the presence of these resistance genes formed the basis of our classification. Furthermore, our analysis additionally suggested that bla_OXA-312_ may also be associated with carbapenem resistance (16 true positives and 0 false positives, Fisher test p = 0.0005016, Supplementary Table 27), therefore bla_OXA-312_ was also added to the list of resistance determinants. However, the presence of ISAba1 alone or the absence of carO proved to be poor predictors of carbapenem resistance and were not added to the list. Finally, we considered an isolate fluoroquinolone-resistant if it carried at least one of the three point mutations defined above.

### Analysis of population structure and phylogeography

#### Quality filtering of genomes

For all subsequent analysis, we only kept genome assemblies where a) assembly coverage was at least 25x, b) MLST could be predicted; c) CPS type could be predicted with at least "Good" confidence (i.e. “The locus was found in a single piece or with ≥ 95% coverage, with ≤ 3 truncated/missing genes and ≤ 1 extra gene compared to the reference”); we eliminated genomes where d) GC content was extremely high or extremely low; e) number of contigs was extremely high; f) length of the longest contig, g) N50 (length of the shortest contig which, together with all longer contigs, represent 50% of the nucleotides), or h) N95 (same for 95% of the nucleotides) was extremely low, i) number of ambiguous nucleotides was extremely high, j) BUSCO v5.4.4^32^ complete score was lower than 95, k) taxonomic classification using sequence data with Kraken 2 v2.1.2^34^ suggested a species other than *A. baumannii* or l) the Kraken2 taxon frequency for the species was extremely low (based on Inter-Quartile Ranges, extreme outliers).^33^ We retained only genomes with metadata indicating m) the country of origin for the isolate and n) at least the collection year of the isolate. Biosample metadata from NCBI was extensively curated to o) eliminate samples that were not human-associated or p) to eliminate known duplicates. Following filtering, only genome assemblies of 11,129 *A. baumannii* isolates were analysed further from the 15,829 that were included in the study.

#### Controlling for sampling bias by downsampling CRAB isolates

The set of 11,129 isolates with genome sequences is likely to be inherently biased due to overrepresentation or underrepresentation of certain regions and time periods compared to others. Such biases could reflect regional and temporal differences in reporting, rather than genuine epidemiological differences, and could therefore distort analyses of population structure (e.g. assessment of serotype diversity or regional dynamics of serotype frequencies). To address this, the 11,129 isolates were downsampled by geographical region and time to account for the overrepresentation of samples originating from the same outbreaks. Specifically, we applied the following downsampling procedure: if isolates from the same city, same year, and same month of the year had the same MLST-CPS type, they were considered samples from the same outbreak, and only one randomly chosen isolate was kept for each month for each city. For isolates where the city was not known, we applied the rule as follows. If isolates from the same country, same year, and same week of the year had the same MLST-CPS type, they were considered samples from the same outbreak, and only one randomly chosen isolate was kept per week. Using this downsampling procedure, we reduced the number of isolates from 11,129 to 6,778. This data set was further filtered for isolates that were classified as CRAB and this final dataset of 5,225 (4,860 since 2009, 3,275 since 2016, 3,150 since 2016 in selected regions) downsampled CRAB isolates were used in downstream analyses of population structure and phylogeography (Fig 2C-E, Fig 3 A,B,D,E, Fig S2, Fig S3, Fig S4, Fig S5, Fig S6, Fig S8).

#### Defining prevalent and global MLST-CPS types

An MLST-CPS serotype was defined as prevalent if it accounted for at least 5% of the isolates in at least one of the 23 world regions. The 23 world regions were defined according to the World Bank Development Indicators database (legacy), accessed through the R package countrycode v1.3.0.^35^ Furthermore, a serotype was classified as global if (i) it was prevalent according to the above definition and (ii) it accounted for at least 2% of the isolates in at least 3 world regions spanning at least 2 continents. Prevalent or global serotypes were identified using the 5,225 downsampled CRAB isolates and were identified separately for isolates collected between 2016-2022 (current period) and for isolates collected between 2009-2015 (preceding period) to address temporal dynamics.

#### Temporal dynamics of the relative prevalence of serotypes

We investigated the temporal dynamics of all MLST-CPS types that had been classified as prevalent (see above). For this analysis, we used the dataset with 4,860 downsampled CRAB isolates since 2009. For each MLST-CPS type, we (i) identified the year when the first isolate was documented since 2009, (ii) filtered the dataset to include observations since that date, (iii) classified each isolate as belonging to the particular MLST-CPS type or not, and (iv) applied logistic regression analysis to study trends in relative prevalence over time. We studied temporal trends both globally and at the level of individual continents. To account for potential biases caused by geographical differences within the studied regions, we included the continent as a covariate when inferring temporal trends on a global scale. In a similar vein, we included world regions as a covariate when inferring temporal trends for continents. Note that we only focused on MLST-CPS types with at least 50 isolates available in a given region, e.g. a particular MLST-CPS type on a particular continent was only considered if at least 50 isolates of that MLST-CPS type were collected from that continent since 2009. We applied FDR correction (through the R package stats^36^) to account for multiple comparisons (Supplementary Table 4). Finally, we plotted the 15 statistically significant MLST-CPS trends calculated based on individual continents (Fig S6).

#### Selecting regions with sufficient sample sizes

For analyses that require sufficient representation of local MLST-CPS diversity, we carried out rarefaction analyses. Specifically, we manually included world regions or countries based on rarefaction analysis indicating that sufficient numbers of isolates are available to represent the local diversity of MLST-CPS types. For the rarefaction analysis, we used the 3,275 downsampled CRAB isolates since 2016 and counted the number of isolates belonging to each MLST-CPS type in each world region (Fig S2) or country (Fig S4) and calculated the rarefaction curves from these count matrices using the R package vegan v2.6-4.^37^ After plotting the rarefaction curves, we manually selected 9 out of 16 regions to study the global distribution of CRAB strains (Fig 2 C,D, Fig S3). Similarly, we selected 7 European and 7 non-European countries for comparisons between countries (Fig 2E, Fig 3B, Fig S5). The European countries included France, Germany, Greece, Hungary, Italy, Romania, and Serbia, while the non-European countries included Brazil, China, Israel, Saudi Arabia, South Africa, Thailand, USA.

#### Building time-calibrated phylogenetic trees

The 11,129 filtered genomes were used to build 4 separate time-calibrated phylogenetic trees for 4 sequence types: ST1, ST2, ST492, ST636. These STs cover the most prevalent serotypes worldwide and the most prevalent serotypes in our focal countries (Fig 3C, Fig S10A-K). For each sequence type, a reference genome was selected first. For ST2, the reference genome was selected manually (GCF_003288775.1, a Complete Genome from the NCBI RefSeq database, sequenced using PacBio, 120x coverage, assembled into two contigs, one of which is labelled as chromosome), for the rest of the sequence types we selected the genome with the longest contig. In the case of ties, we selected the earliest genome. Then, each genome belonging to a particular sequence type was mapped to its reference genome using snippy v4.6.0^38^ to produce pseudo-whole genomes. For isolates that were sequenced in the present study, we mapped the reads, while for the rest of the isolates we mapped the assembled genomes. We kept the longest contigs, removed any duplicates (either genuinely duplicate sequences or sequences that only became duplicates after pseudo-whole genome reconstruction) and constructed the phylogenetic tree using gubbins v3.3.0,^39^ (model fitter: raxmlng, tree builder: fasttree, maximum number of iterations: 10) to account for recombinations. Abnormally long branches were subsequently removed with TreeShrink v1.3.9.^40^ The resulting tree was dated in two steps: the tree was first rooted using root-to-tip regression with the lowest sum of the squared residuals and then the rooted tree was dated with treedater^41^ without rerooting, using a strict molecular clock. After dating, tips that had been removed due to sequence duplication were added back to each tree with 0 branch lengths.

#### ST2 global and regional transmission dynamics

To study the phylogeographic patterns of the ST2 sequence type (Fig 3D,E, Fig S8), we built on a previously published method^42^ and analysed genetic similarity between pairs of isolates across different temporal and spatial scales. The general idea behind the analysis is that pairs of isolates may be geographically close or distant and also genetically close or distant, and the distribution of isolate pairs across geographic-genetic distance categories characterises the transmission dynamics of the bacterium. For example, pairs that are both genetically and geographically close indicate local spread, while pairs that are genetically close but geographically distant indicate cross-border transmissions. Therefore, the number of pairs in these categories yields insights into the relative importance of local versus cross-border transmissions.

For the analysis, we used the time-calibrated ST2 tree reconstructed above. However, we only included CRAB isolates and focused on the downsampled dataset to address sampling bias. This procedure resulted in a smaller ST2 tree with 3,537 tips. In the case of global transmission dynamics (Fig 3D), we considered pairs of isolates on the tree that were collected at most 2 years apart and categorised these pairs by geographic location (same country, different country, etc.) and genetic distance (based on the most recent common ancestor, MRCA). Then we counted the number of pairs in each combination of spatial and temporal categories and calculated relative risks separately for each temporal category. Specifically, we calculated the relative likelihood of occurrence in a given spatial category as compared with the likelihood of occurrence in the control spatial category. For example, we found that when focusing on pairs with MRCA below 6 years, the relative risk is approximately 10 for pairs that originate from the same country as compared to those that originate from different countries on the same continent (reference category).

To account for uncertainties associated with this analysis (e.g. sampling of isolates and tree building), we took multiple subsamples from the set of 3,537 isolates constituting the ST2 phylogenetic tree above (100 subsamples with 500 isolates each). For each subsample, we reconstructed a separate phylogenetic tree and analysed it as described above. To reconstruct such a subsampled tree, we selected the respective recombination-free polymorphic sites from the Gubbins output, filtered the subsampled sites using SNP-sites v2.5.1,^43^ built a phylogenetic tree using FastTree v2.1.11 SSE3,^44^ and performed rooting and dating the same way as for the ST1, ST2, ST492, and ST636 trees. Then, we calculated relative risks for each tree and finally, using the 100 subsamples we calculated the mean values and 90% confidence intervals for these risks. Using 90% instead of 95% reflects that the relationships we seek to infer require one-sided tests.

We followed a similar procedure for the phylogeographic analysis of CRAB in Europe (Fig S8), and also for regional transmission dynamics (Fig 4E). For the regional analysis, we looked at pairs that were collected at most 0.5 years apart and drew 500 subsamples with 100 isolates each.

#### Capsule visualisation of *A. baumannii* ST2-KL3 isolates

To visualise the capsule of different *A. baumannii* clinical isolates and induced phage-resistant isolates, we performed Transmission Electron Microscopy (TEM) imaging by following a previously described protocol.^45^ Briefly, 0.5 ml of overnight culture from each bacterial isolate (n = 4) was centrifuged for 2 min at 13000 rpm in a benchtop centrifuge and then fixed overnight at 4 °C with Karnovsky fixative solution (Karnovsky, 1965) (pH 7.4). After fixation samples were briefly rinsed in distilled water (pH 7.4) for 15 min and fixed furthermore in 1% osmium tetroxide in ddH2O (Sigma-Aldrich, St. Louis, MO, USA) for 1 h. After fixation, samples were briefly rinsed in distilled water for 10 min, then dehydrated gradually in ethanol. Afterwards, bacteria were polymerized through propylene oxide (Molar Chemicals) and then embedded in an epoxy-based resin (Durcupan ACM; Sigma-Aldrich) for 48 h at 56°C. From the resin blocks 50 nm thick ultrathin sections were cut using an Ultracut UCT ultramicrotome (Leica; Wetzlar, Germany) and were mounted on a single-hole, formvar-coated copper grid (Electron Microscopy Sciences; Hatfield, PA, USA). The contrast of the samples was enhanced by staining with 2% uranyl acetate in 50% ethanol (Molar Chemicals, Electron Microscopy Sciences) and 2% lead citrate in distilled water (Electron Microscopy Sciences).

#### Isolation of *A. baumannii* bacteriophages

The phage hunt was performed on *A. baumannii* wild-type isolates belonging to different MLST-CPS types (Supplementary Table 13). In all cases, phages were isolated from raw sewage water harvested from waste-water treatment plants from five Hungarian cities (Szeged, Budapest, Hódmezővásárhely, Debrecen, Békéscsaba), using the enrichment procedure described previously^46^ with some modifications. In brief, the sewage water was cleared by centrifugation (4500 rpm for 10 min), followed by filtration through a 0.45-μm-membrane filter and aliquots of wastewater were mixed with an equal volume of 2× LB medium and 0.1% of overnight bacterial culture. Samples were incubated overnight at 37°C with shaking at 140 rpm followed by centrifugation for 10 min at 4500 rpm, and then the supernatant was filtered through a 0.45 μm membrane filter twice to remove the residual bacteria and debris. Plaque assay was performed to screen for the presence of lytic phage activity using the double-layer agar method.^47^ The plates were incubated overnight at 37°C and examined for zones of lysis or plaque formation. Single plaques formed on the lawns of *A. baumannii* strains were picked up and this process was repeated three times, in order to obtain pure phage stock.

#### Phage propagation and purification

Phage propagation was carried out using a liquid culture of the corresponding *A. baumannii* clinical isolate (OD_600_=0.6) at a multiplicity of infection (MOI) of 0.1. 25 ml of phage lysate was concentrated by the addition of 10% polyethylene glycol (PEG) M.W. 8000 and 1M NaCl and incubated overnight at 4 °C. After centrifugation at 4600 rpm for 20 min, the pellet was suspended in 1 ml SM buffer (50 mM Tris-HCl [pH 7.7], 100 mM NaCl, 80 mM MgSO4). The phage titer was determined by the double-layer agar method and the titer was reported as a plaque-forming unit (PFU/ml).

For *in-vivo* experiments, phage stocks were prepared using the Phage-on-Tap protoco^l48^ with some modifications. The corresponding *A. baumannii* culture was grown in 100 ml LB at 37°C to an OD_600_ of 0.6-0.7 and infected with the phage. The phage lysate was concentrated (10X) using centrifugal filters (Amicon Ultra-15, Sigma 100 MDa cut off) and the concentrate was washed with 1X PBS at least 5 times. The phage concentrates were mixed with 0.7 volume of 1-octanol for additional endotoxin removal,^49^ shaken for one hour at room temperature following an incubation at 4°C for 2 hours. The phage-containing phase (bottom phase) was collected with a syringe after centrifugation at 4,000 × g and 4°C for 10 min. The removal of octanol was enhanced by repeating the centrifugation step two additional times. This was followed by a few extra washing steps with PBS using Amicon filter (100 MWCO) devices. To ensure that the endotoxin concentration is low, an additional step was included using Pierce high-capacity endotoxin removal spin 1 ml column (Thermo Scientific) following the manufacturer’s instructions.

### Laboratory characterisation of the isolated phages

#### Phage morphology

In order to obtain transmission electron microscopy images, six µl of the phage lysate was mounted on a TEM copper grid (300-mesh, Electron Microscopy Sciences) with carbon-coated ultrathin formvar film.^50^ The samples were dried using filter paper after 1 minute. To obtain the negatively stained samples, the samples were contrasted with 6 µl uranyl acetate (2% w/v) in 50% ethanol for 2 min (this process was repeated 3 times). After the removal of the excessive staining solution, samples were dried under a Petri dish for 2 h before the electron microscopic evaluation. Negatively stained samples were systematically screened and the grids were examined under a transmission electron microscope (JEM-1400 Flash transmission electron microscope) at 80kV with a 16 MP Matataki Flash scientific complementary metal–oxide– semiconductor (sCMOS) camera (JEOL).

#### Phage adsorption assay

10 ml bacterial culture of wild-type isolates or induced phage-resistant isolates was pre-grown until exponential phase (OD_600_=0.6-0.7) and mixed with the phage at MOI 0,1. The cells were incubated at 37°C with aeration and at each time point 100 μL samples were taken every 5 or 10 min for 30 min. Samples were mixed with 850 μl of LB supplemented with 50 μl of chloroform, vortexed, and then centrifuged for 1 min at 13000 rpm in a benchtop centrifuge. The supernatant was diluted in LB to determine the unadsorbed phages by the plaque assay using the double-layer method. For details, see Supplementary Table 22 and Fig 6E.

#### Determination of phage host specificity

The lytic activity of each phage was screened against multiple *A. baumannii* clinical isolates (Supplementary Table 10) with the standard spot assay as previously described.^51^

For Silvergun phage, we compared the efficiency of infection on different ST2-KL3 and ST2-KL2 isolates respectively, by determining the efficiency of plating (EOP) (supplementary Note S1). Briefly, serial dilution of phage lysate was plated on soft agar overlays of different bacterial isolates. After incubating the plates overnight at 37°C, the individual plaques were counted and the average of 3 technical replicates was calculated and compared to the number of plaques on the original host of isolation. EOP was calculated as follows: average PFU on target bacteria / average PFU on host bacteria (see Supplementary Table 28).

### Bioinformatic analysis of the isolated phages

#### Whole-genome sequencing

Phage genomic DNA was extracted using a commercial phage DNA isolation kit (Norgen Biotek Corp.; 46850) following the manufacturer’s instructions with the sole exception of applying sterile, distilled H_2_O during the elution step instead of the provided elution buffer. Afterwards, samples were sequenced by Illumina sequencing. We used 1 ng of DNA as the input amount recommended in the Nextera XT Sample Preparation Guide. We followed the rest of the protocol as written. Sequencing-ready libraries were quality control checked by BioAnalyzer2100 instrument using High Sensitivity DNA Chip (Agilent Technologies USA, Cat. No. 5067-4626). NGS was carried out on the NextSeq 500 sequencing system with NextSeq 500/550 Mid Output Kit v2.5 (300 Cycles) chemistry (Illumina, Inc. USA, Cat. No. 20024905). Applied read length was in the 150-300 bp range. Processing of the sequencing data was started with the quality control and trimming of reads using Trimmomatic v0.36^52^ with the following command: TrimmomaticPE -threads X [reads_Fw].fastq.gz [reads_R].fastq.gz [Fw]_paired_output.fastq.gz [Rev]_unpaired_output.fastq.gz [Fw].fastq.gz [R]_unpaired_output.fastq.gz ILLUMINACLIP:TruSeq3-PE-2.fa:2:30:10 LEADING:3 TRAILING:3 SLIDINGWINDOW:4:15 MINLEN:3.

Assembly of the thus obtained, filtered reads into contigs was carried out by either MEGAHIT^53^ Li or MetaSPAdes v.3.13.0^54^ or meg on a per-sample basis. Contigs of 10 kb < length were then investigated to identify phages in the samples by BLAST+ v2.12.0 blastn alignment to NCBI RefSeq database,^26^ using the following command: blastn -query [query].fasta -db [database] -outfmt 5 -max_target_seqs 1 -qcov_hsp_perc 5 -out[output].xml-num_threads X

#### Bioinformatic analysis

Bioinformatic prediction of ORFs and annotation of the phage contigs was done by a simple customary python script. In brief, the code translated the whole input nucleotide sequence into amino acid sequence in all six reading frames. Then, regions with a length longer than 50 from start to stop codon, were collected in a list along with the positional and start-end values and the assigned identifier of the candidate gene. The collected amino acid sequences were matched to the NCBI nr database then the name and taxonomic information of the best hits were added to the identifier of the candidate subject (if any). The above data was then collectively processed to generate GenBank files for the individual phage contigs. Further, genes of interest i.e. genes related to therapeutically important functions were listed in separate files. Genes of interest were extracted from the prediction data with regular expressions together with the positional information (Supplementary Note S1).

The therapeutic suitability and the possible life cycle of isolated phages were predicted with PhageAI software using default settings^55^ and the Phageleads web service.^56^ Endonucleases were identified by blastp alignment. The presence of antibiotic or virulence-conferring genes were checked by utilising the RESFINDER v3.0 and VIRULENCEFINDER v1.5 database, respectively, corroborated with PhageLeads results. Insertion sequences were screened using ISfinder.^57^ Depolymerase prediction was done by Depolymerase Predictor. A tail fiber or spike protein was deemed as carrier of a depolymerase region if the value of probability was 0,85<.^58^ The completeness of assembled phage genomes was elucidated via CheckV software^59^ (Supplementary Table 13).

### Phage pangenomic network construction

To visualise the location of our phages in the broad landscape of viruses, we used the classical approach of constructing an orthogroup-based network of viruses, where proteins of all participant phages are clustered, and the phages’ level of connection is determined by the amount of proteins they have in common viral clusters (VCs).The tool we found to best suit this task was vConTACT2^62^, which implements the above methods and outputs results largely adhering to the classifications of the ICTV database. To acquire the data for the network, we retrieved all phage genomes assigned to the Caudovirecetes order from the International Committee on Taxonomy of Viruses (ICTV) database^60^ (version August 22, 2022).

We also conducted literature mining to identify Acinetobacter phages that have been used as therapeutic agents (Supplementary Table 12). In addition, we included Acinetobacter phages from our collection, which were obtained through an extensive phage hunt program described in this paper.

We obtained taxonomy information for the accession IDs originating from the ICTV database, while for our own phages, we conducted a blast search to identify their tail fibers and spikes. Taxonomy information was obtained based on the closest relative’s taxonomic position.

Prodiga^l61^ was used to predict ORFs for all phages, and these ORFs were used as input for vConTACT2’s gene2genome module to generate a network file in Cytoscape format. A custom R script was then used to merge additional metadata information for the nodes visible in the network.

The final network file was then created with vConTACT2’s default settings, no database, the ‘BLASTP’ method, and c1 clustering. To ensure reproducibility and scalability, the entire process was implemented in a Nextflow pipeline. The network subplots in Fig 5 were created in Cytoscape.

### Average Nucleotide Identity (ANI) of phages

We compared the genomes of our phages with those considered therapeutically important or known to have a lytic life cycle (Fig5E). The alignments resulting from this comparison were handled as follows: for a specific section of the genome, if multiple alignments were possible, we chose the one with a higher identity. In cases where an alignment with lower identity covered a longer section, it was used for the entire section. In situations where one genome was longer than the other, we adjusted the Average Nucleotide Identity (ANI) value by subtracting the length difference. For instance, if one genome was 90% the length of the other, even if the genomes exhibited 100% identity, the maximum ANI value achievable was 90%.

### Phage pangenomic visualisations

The genomic synteny of phages was colinearised according to a reference point (an official lytic or therapeutic phage in each case) and then the GenBank files were supplied to Clinker.^63^ The pangenomic comparisons of phages were done with a custom R script that compared each of our phage genome’s genes against the corresponding one from the reference genes using blastn. Then from all compared tail coding sequences, we predicted depolymerase regions using the Phyre2 web platform,^64^ where we selected the best hit among those that had above 0.4 confidence and contained the word ‘hydrolase’ and ‘lyase’, corresponding mostly to the methods described earlier.^65^ For depolymerase comparisons (Supplementary Table 17, Fig S13), we only used the depolymerase regions.

### Isolation of *in vitro* evolved phage-resistant *A. baumannii* mutants

Phage-resistant variants were evolved in a 96-well microtiter plate in 200 µl final volume in LB medium. The OD_600_ values of the wells were measured every 20 min for 24 hrs in an EPOCH 2 (Biontech) instrument, at 37°C with 10 second shaking before each measurement. Each well, except the control wells, contained 100 µl from the bacterial culture with OD_600_ = 0.7 and the phage(s) in 10^6^ PFU/well concentration. Resistance was evolved to a single phage or a combination of 2, 3, or 4 different phages. If phage combinations were used, each phage was added in 10^6^ PFU/well concentration (in 1:1 ratio). Following a 24-hour incubation period, the wells that exhibited cell growth as indicated by OD_600_ values, samples were streaked onto plates containing chromogenic agar and incubated overnight at 37°C for each strain. The resulting individual colonies were restreaked at least 3 times to avoid phage contamination. The phage-resistant phenotypes were confirmed using phage susceptibility tests including the double-layer agar plate method or *in vitro* growth curves.

The phage-resistant genotype was determined by detecting the mutations in the assembled genomes using a custom pipeline. For mutations occurring in the K-locus, we identified the genes of the K antigen biosynthetic pathway using blastn 2.13.0+.^66^ We aligned the blast hit to the reference gene with mafft v7.520 ^67^ and with a custom script identified the nucleotide and the amino acid variants.

### Antimicrobial susceptibility testing of phage-resistant isolates

Minimum inhibitory concentrations (MICs) of five clinically relevant antibiotics: Colistin (Pharma), Meropenem (Bioscience), Imipenem (MedChemExpress), Levofloxacin (MedChemExpress), Trimetoprim: Sulphametoxazole (1:5 ratio) (Sigma, MedChemExpress) were determined using the standard microbroth dilution protocol^68^ and interpreted using the European Committee on Antimicrobial Susceptibility Testing (EUCAST) guidelines.^1^ Cells were grown in Mueller Hinton Broth 2 (MHB2) (Millipore) and the inoculum size was set to 5 x 10^5^ bacteria per ml. Among the tested antibiotics were two carbapenems (Meropenem and Imipenem) and the last-resort antibiotic Colistin. In order to maximise reproducibility and accuracy, we used a robotic liquid-handling system (Hamilton) to automatically prepare 7-step, two-fold serial dilutions in 384-well microtiter plates. After 18 h of incubation at 37°C, raw OD_600_ nm values were measured in a Biotek Synergy microplate reader. MIC was defined by a cutoff OD_600_ nm value (mean +2 s.d. of A600 nm values of bacteria-free wells containing only growth medium). MIC fold change was calculated as MIC_wild-type_ strain / MIC_phage-resistant_ strain.

### Efficacy of phages in *Galleria mellonella* larvae model

We used *Galleria mellonella* larvae as one animal model to evaluate the effectiveness of Highwayman and Silvergun phages alone and in combination with other phages against *A. baumannii* clinical isolates. For each experimental condition, 10 larvae were used and experiments were repeated independently at least 3 times. Only larvae exhibiting a uniform cream colour were used for the experiments. Survival of the larvae was monitored for 48 hr, every 6 hr.

As a first step, we determined the proper inoculum size to be used in further experiments. 6 different *A. baumannii* isolates and 5 different inoculum sizes for each isolate (CFU = 9×10^6^, 9×10^7^, 9×10^8^, 9×10^9^, 9×10^10^ per ml) were tested. Based on these preliminary results, the inoculum size of 9×10^8^ CFU/ml was chosen as the mortality rate of the larvae at this inoculum size was 50% at 18-hrs post-infection for half of the tested isolates.

Prior to the infection of the larvae, the bacterial strain was grown overnight in MHB2 medium at 37°C with aeration. To remove the growth media, the bacterial culture was washed twice with 1X Phosphate-Buffered Saline (PBS) and the density was set to 9×10^8^ CFU/ml. From this, 10 μl was injected into the first proleg of the larvae. Phage treatment was administered in the same volume in PBS, in the opposite proleg, 5 minutes after the bacterial infection. This time point was chosen based on the results of preliminary experiments where we compared the survival rate of the larvae that received the same phage treatment 5 minutes and 60 minutes post-infection and found that the survival rate of the larvae receiving the treatment 5 minutes post-infection was significantly higher.

### Isolation of *in vivo* evolved phage-resistant *A. baumannii* mutants

We isolated phage-resistant *A. baumannii* mutants from larvae 48 hours post-infection, which marked the endpoint of the survival experiment, or earlier if larvae died before this time point. On the larva’s ventral side, an incision was made with a sterile blade, and then we streaked from the body lumen onto Acinetobacter-specific chrome agar (CHROMagarTM Acinetobacter, CheBio). The resulting colonies were re-streaked at least 3 times to obtain a pure bacterial culture. The resistance phenotype was confirmed by phage typing against the phage(s) used in the treatment of the larvae and, in addition, we checked the susceptibility of the isolate against the other cocktail components, PhT2-v1 and Fanak as well. If an altered phage infection pattern was detected compared to the wild-type, we sent the isolate for sequencing.

### Efficacy of phages in intraperitoneal mouse model

We created a mouse intraperitoneal model to study the infectivity of Aci 15 *A. baumannii* isolate. Female BALB/c mice, 6-7 weeks old and weighing 17-18 g, were obtained from Envigo in the Netherlands. All animal care and handling procedures adhered to the European Federation for Laboratory Animal Science Association (FELASA) guidelines, and the EnivoInvest Co., Hungary’s Animal Welfare committee approved the protocols (permit number: BA02/2000-12/2022).

As a first step, for the Aci 15 isolate, we determined the proper inoculum size to be used in further experiments. For this, 5 different inoculum sizes (CFU = 10^5^, 10^6^, 10^7^, 10^8^ and 10^9^) were tested. Prior to the experiment, one colony of the Aci 15 isolate from a one-day-old LB agar plate was transferred into 5 mL LB liquid medium and was cultivated for 14 hours in a shaker thermostat (120 rpm, 37 °C). On the next day, the liquid culture was centrifuged (2 minutes, 10.000 g) and washed once with PBS. The gained bacterium suspension was serially diluted (10x) to obtain different inoculum sizes. From these dilutions, 200 μl was administered intraperitoneally to each mouse. Mice were separated into groups (each containing 5-5 animals) based on the infection doses. Weight and death rates of the infected and control animals were recorded in the subsequent 7 days. Based on these results, the inoculum size of 10^9^ CFU/mice was chosen for phage rescue experiments.

Therapeutic efficacies of single (phage Highwayman alone) and combined (HSFPh phage cocktail) purified bacteriophages against Aci 15 isolate were tested as described before,^69^ with slight modifications. For this purpose, mice (20 animals) were divided into 4 groups (G1-G4). Members of the first three groups (G1-G3) were infected with 10^9^ CFU/mice, while mice in group G4 received only from the phage suspension mix and served as phage controls (see Fig S18H). Mice in G1 served as bacterium controls without phage administration. In contrast, members of G2 received phage Highwayman (MOI 6); while members of G3 received the HSFPh phage cocktail (MOI 6:1:1:1). Phages in the case of G2 and G3 were administered 10 minutes after the injection of bacteria. Death rates and body weights of all animals were daily recorded for seven days after the treatments.

### Comparing the cell surface properties of wild-type and phage-resistant *A. baumannii* isolates with Fourier-transformed infrared spectroscopy

We adapted the protocol described before^70^ with minor changes. Briefly, according to the manufacturer’s instructions, bacterial biomass was collected from LB agar plates. The isolates were incubated for 24 h at 37°C. After incubation, bacterial suspensions were prepared by adding a full inoculation loop (1 μl) of bacteria in 1.5 ml Eppendorf tubes containing 50 μl of 70% ethanol and 2 mm metal beads provided by the supplier. The bacterial suspension was thoroughly vortexed after which 50μl of sterile H_2_O was added. Then, 15μl of bacterial suspension was pipetted on a silicon plate (provided by Burker IR Biotyper equipment) in three technical replicates per isolate (Supplementary Table 23). We prepared two standard suspensions (provided by the supplier) and 10μl were added as controls in two technical replicates. The plate was then dried for 20 minutes in the laminar hood at room temperature. Results were evaluated using Opus Software V7.5.18 and IR Biotyper Software V2.1.0.195 with the default settings (32 scans per technical replicate; spectral resolution, 6 cm^−1^; apodization function, Blackman-Harris 3-term; zero-filling factor 4). Following the manufacturer’s instructions, the pre-processing steps of the acquired data were as follows: the FTIR spectra measurements of individual technical replicates that did not meet the default quality criteria (0.4 < absorption < 2, signal-to-noise-ratio < 40, technical replicate of an isolate was not clustering with the other replicates of that isolate) were manually removed from further analysis to prevent inclusion of wrongful data. The distance matrix provided by the IR Biotyper software was used for further analyses.

### Screening for lysogen activity of the isolated phages with therapeutic potential

As the absence of lysogenic activity is a requirement for phages to be used in the traditional phage therapy,^71^ we tested some of the isolated phages: Silvergun, Highwayman, and PhT2-v2 with therapeutic potential for lysogenic activity by adapting the protocol described before.^72^ Briefly, after performing a spot assay with the phage in question (PFU scaling from 10^5^ to 10^8^) bacterial cells were isolated from mesas, zones of confluent bacterial or phage-resistant cell growth in the centre of the lysis spots. After applying the phage in two different concentrations, twenty colonies were isolated and tested in patch plate screens to see if the phages were capable to lysogenize the cells. This step was repeated twice to eliminate the chance of carry-over of phage particles between the steps. To ensure the absence of the lysogen activity of the phages tested, we performed the spontaneous phage release test and the immunity assay as well. Plates were incubated overnight and phage infection was identified by lysis zones.

### Validation of the wide host range of the Rocket phage

To validate the wide host range of the Rocket phage, we performed plaque PCR to detect the presence of the phage from single plaques formed by the phage on bacterial cultures of different *A. baumanii* isolates belonging to different MLST-CPS strains. As a negative control, we used cells from the same plate from the vicinity of the plaques.

Primers used for the plaque PCR

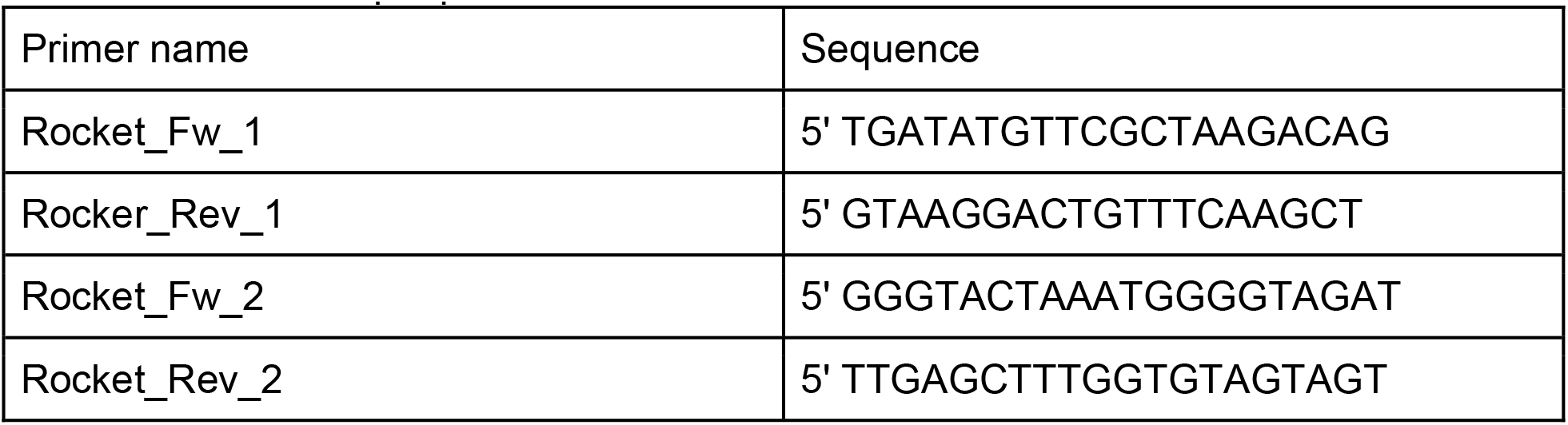

