## Supplemental information for "Pathogen genomic surveillance as a scalable framework for precision phage therapy"

#### Supplementary information

#### Supplementary figures

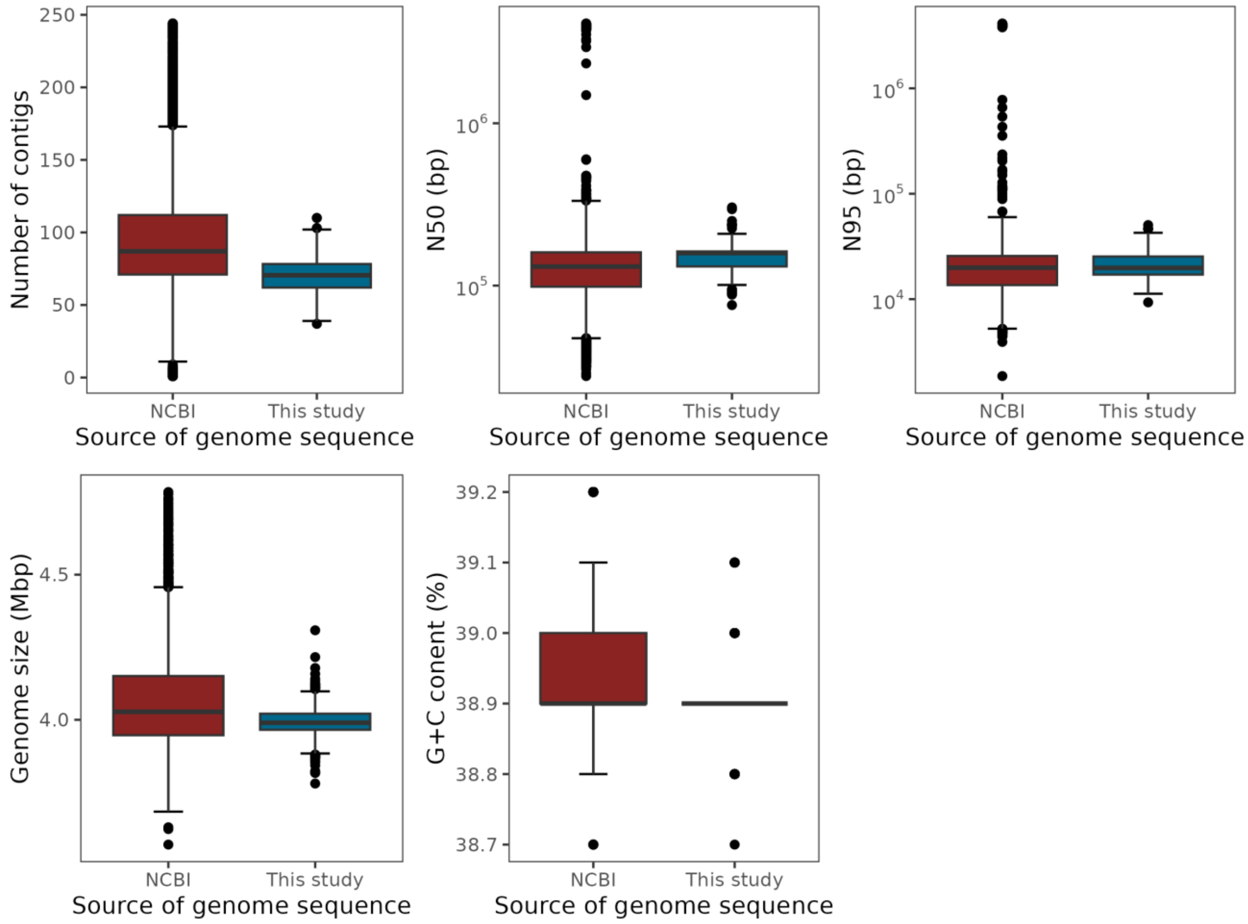


**Fig S1. Assembly quality control metrics for genome assemblies included in this study.** Genomes sourced from NCBI or sequenced as part of this study are shown in red and blue, respectively. The figure describes 8,915 assemblies that passed all quality filters and were classified as CRAB (Methods). N50 indicates the length of the shortest contig which, together with all longer contigs, represent 50% of the nucleotides. N95 indicates the same for 95% of the nucleotides. The whiskers indicate thresholds for 1.5 interquartile range (IQR) outliers. For quality filtering, we used more permissive 3 IQR outlier thresholds. The figure thus shows the isolates that were included in our analysis.

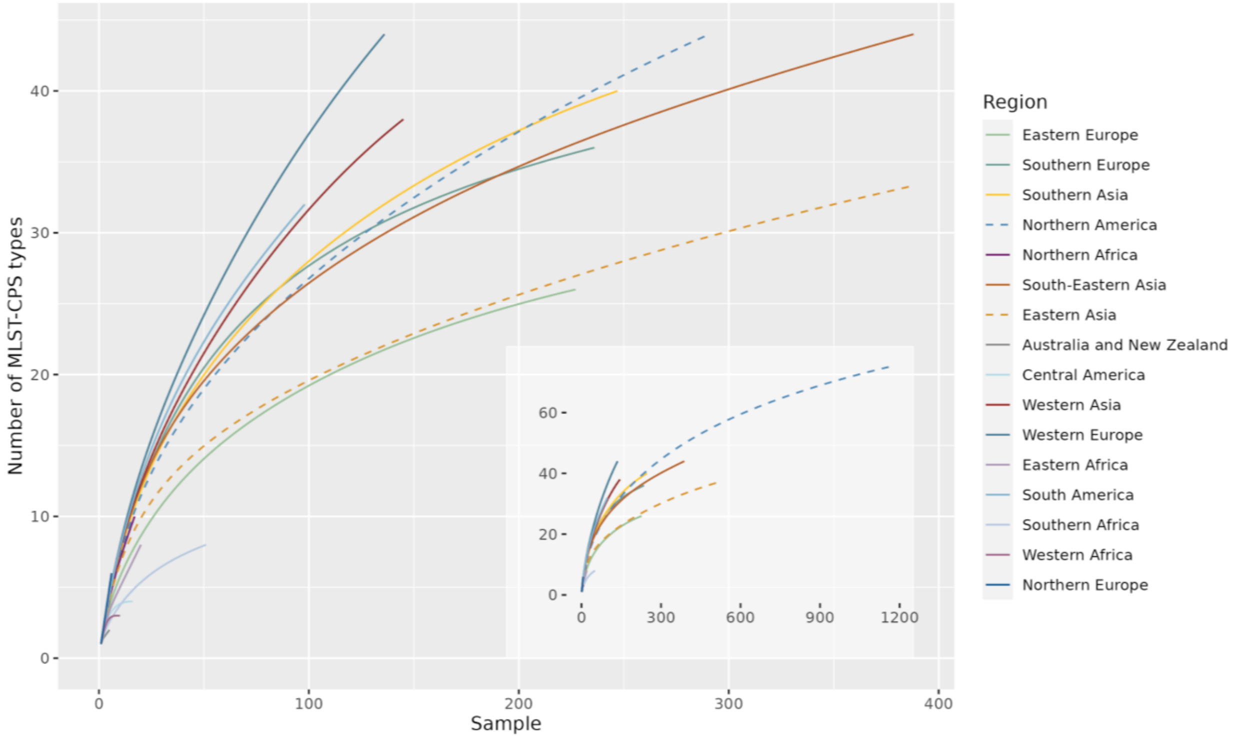


**Fig S2. Rarefaction analysis of 3,275 downsampled CRAB isolates for all geographical world regions where CRAB genomes were available in the period between 2016 and 2022.** Genomes were downsampled to remove sampling bias (Methods). World regions were defined according to the World Bank Development Indicators database (legacy), accessed through the R package countrycode v1.3.0.[1] Axis boundaries on the main plot were truncated to eliminate the distortion caused by Northern America and Eastern Asia from where a large number of isolates were collected. The embedded figure shows the same plot with default axis boundaries. For details of the rarefaction analysis see Methods, selecting regions and countries for comparative analysis.

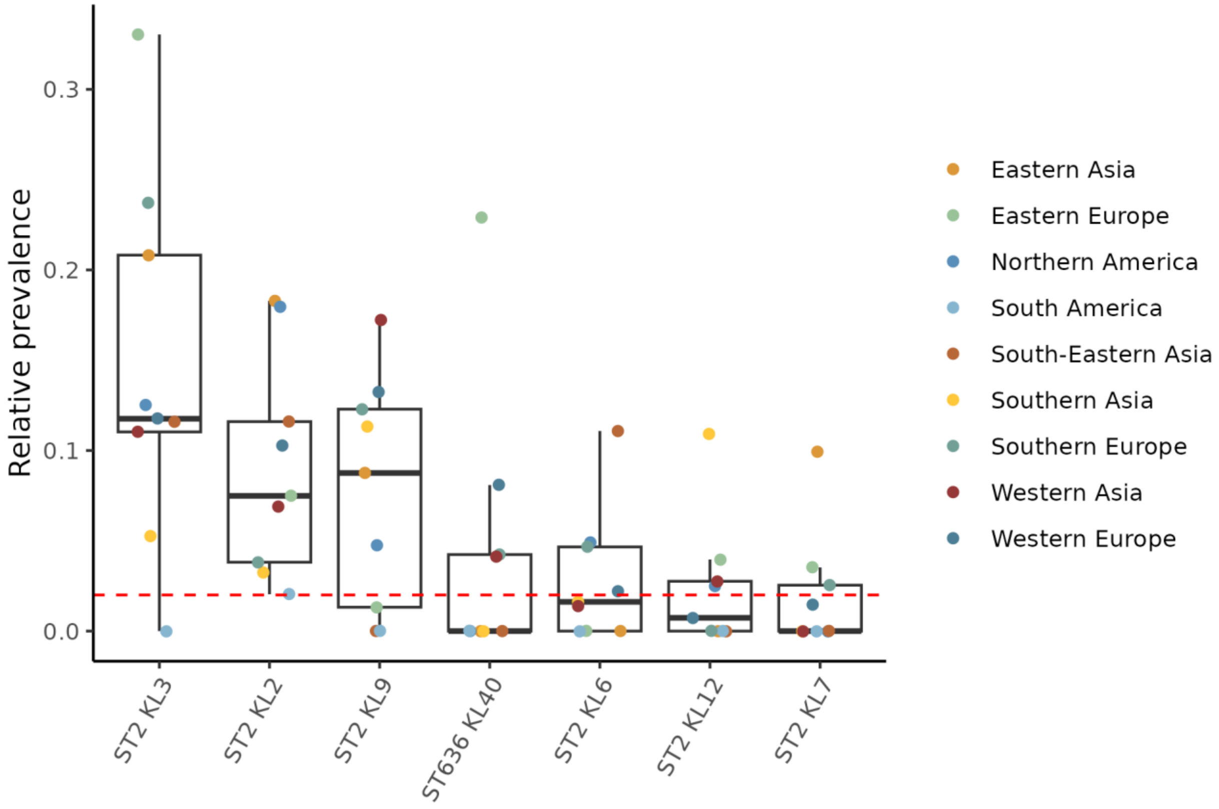


**Fig S3. Relative prevalence of the 7 global MLST-CPS types in 9 selected regions of the world.** Analysis was carried out based on 3,150 downsampled CRAB MLST-CPS types in these regions between 2016 - 2022. Each point represents the relative prevalence of isolates belonging to a certain MLST-CPS type in a certain region. The boxplots describe the distribution of the 9 relative prevalence values for each MLST-CPS type. The higher and lower ends of the boxes indicate 75th and 25th percentiles, respectively, the solid bold black line inside each box indicates the median of the distribution, the whiskers were drawn using standard 1.5 interquartile ranges. The horizontal dashed red line marks the 2% threshold, which is used in classifying an MLST-CPS type as global (at least one world region with 5% and two additional with 2% prevalence, spanning at least two continents). See Methods for more information on downsampling, CRAB filtering, selecting regions or identification of prevalent and global serotypes.

A

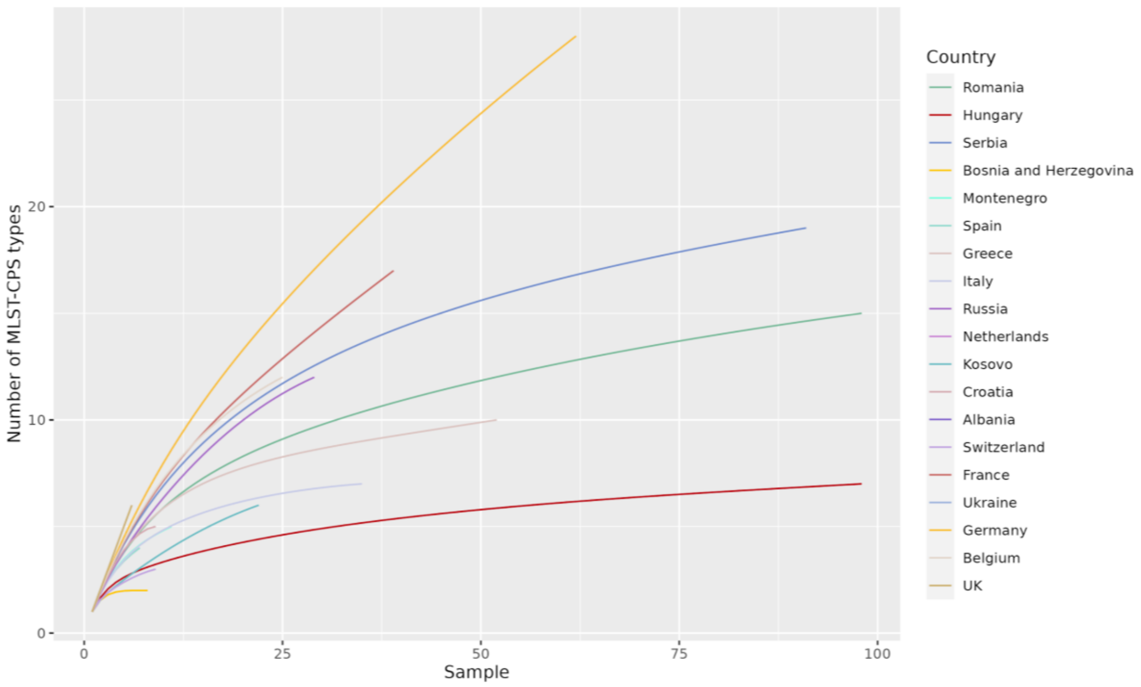


B

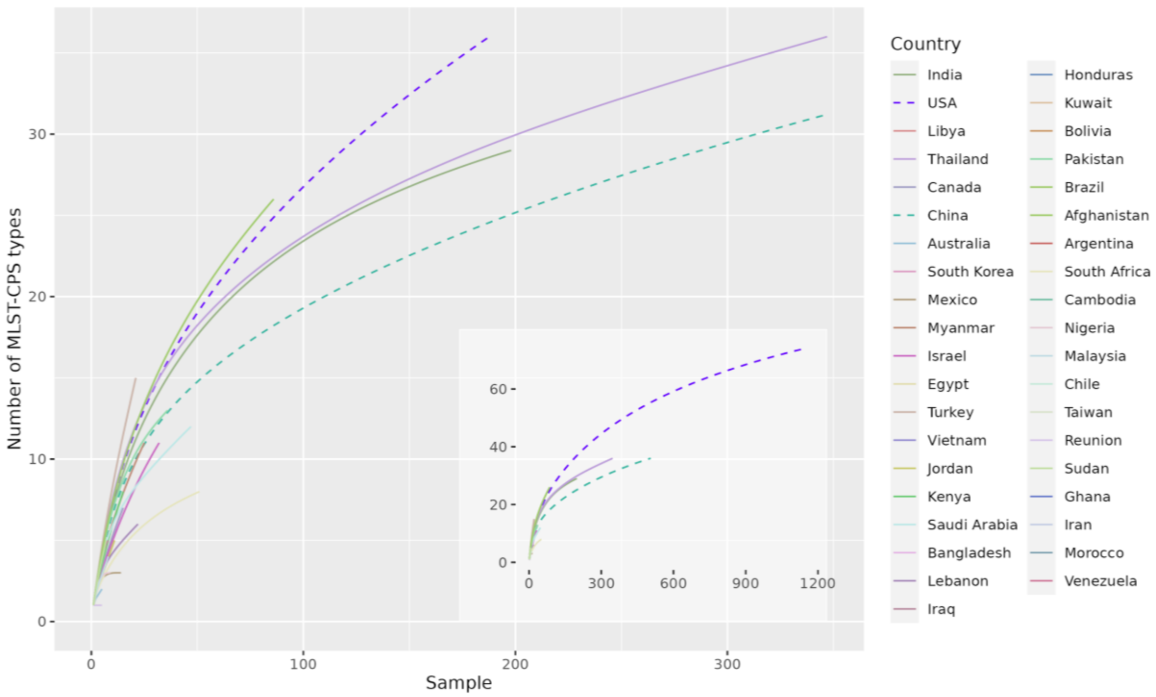


**Fig S4. Rarefaction analyses for countries. A,** Rarefaction analysis for European countries based on 605 downsampled European CRAB isolates from the period between the years 2016 and 2022. For details of the rarefaction analysis see Methods, selecting regions and countries for comparative analysis. **B,** Rarefaction analysis for non-European countries based on 2,670 downsampled non-European CRAB isolates from the period between 2016 and 2022. Axis boundaries on the main plot were truncated to eliminate the distortion caused by China and USA from where a large number of isolates were collected. The embedded figure shows the same plot with default axis boundaries. For details of the rarefaction analysis see Methods, selecting regions and countries for comparative analysis.

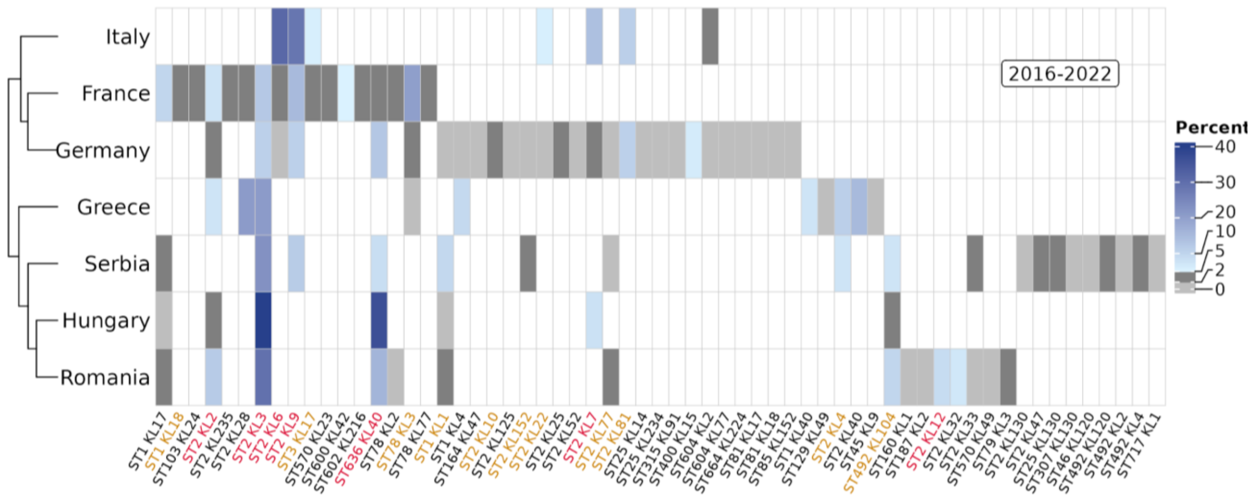


**Fig S5. Relative prevalence of CRAB MLST-CPS types in 7 European countries between 2016 - 2022.** The colour-coding indicates the prevalence of an MLST-CPS type in a world region: 0 (white), 0-2% (light grey), 2-5% (dark grey), 5-40% (shades of blue). Orange and red labels mark MLST-CPS types that were classified as prevalent or global, respectively. The analysis is based on 475 downsampled genomes. See Methods for more information on downsampling, CRAB filtering, or identification of prevalent or global serotypes.

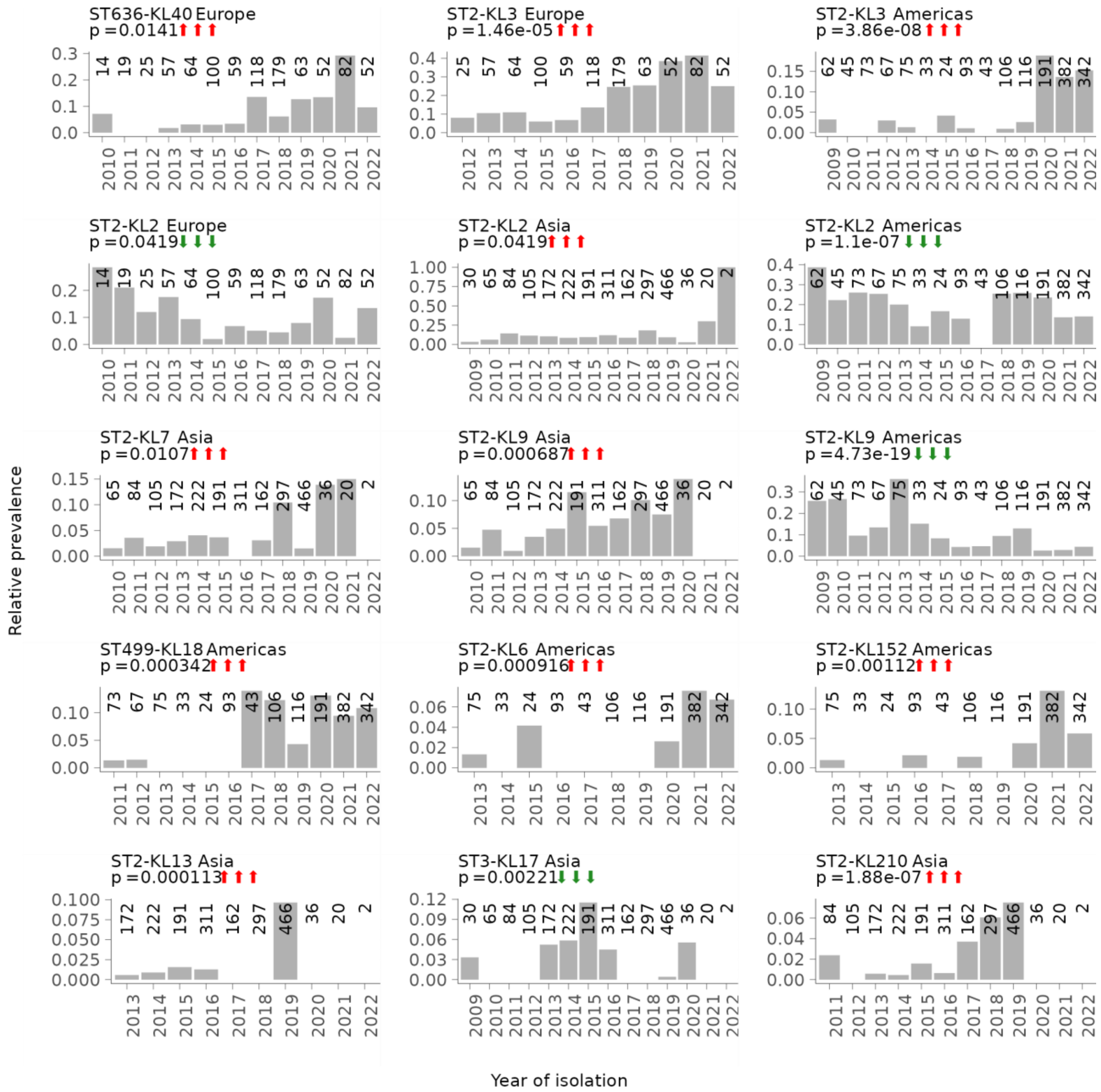


**Fig S6. Relative prevalence of CRAB MLST-CPS types over time.** Relative prevalence was calculated by dividing the number of isolates from a specific MLST-CPS type by the total number of isolates in a given year. Each plot shows a single MLST-CPS type on a single continent. The numbers above each column indicate the overall number of CRAB isolates after downsampling in each collection year (see Methods). Only those MLST-CPS type - continent combinations are shown where we observed a statistically significant increase or decrease over time (red and green arrows, respectively) by statistical modelling (Methods). The p-values were calculated using logistic regression, refer to the statistical significance of the collection date variable and were corrected for multiple comparisons using FDR correction. Data is available in Supplementary Table 4.

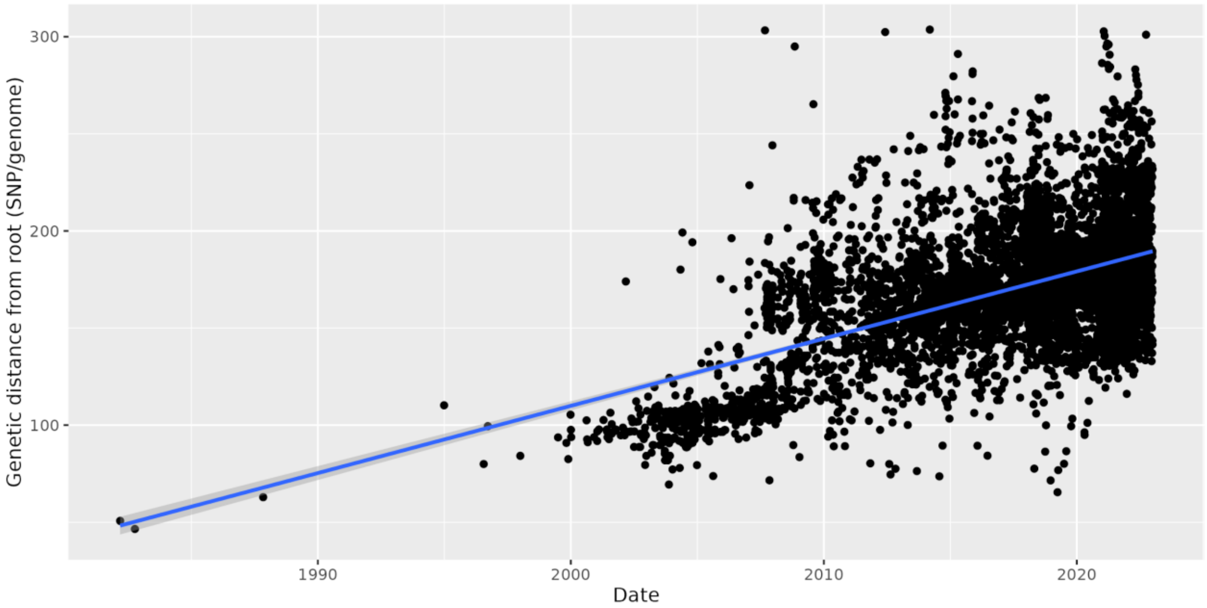


**Fig S7. Root-to-tip regression plot for the global dated ST2 KL3 tree with 7,720 tips**. Time of most recent common ancestor: 1973, root-to-tip mean rate: 3.47 SNP/genome/year for root-to-tip mean rate, root-to-tip $R^2$: 0.279, root-to-tip p-value: 0. The root-to-tip mean rate is in agreement with an earlier study which estimated a mutation rate of ~5 SNP/genome/year for GC1 clones.[2] For more details, see “Constructing phylogenetic trees for relevant sequence types” section of Methods.

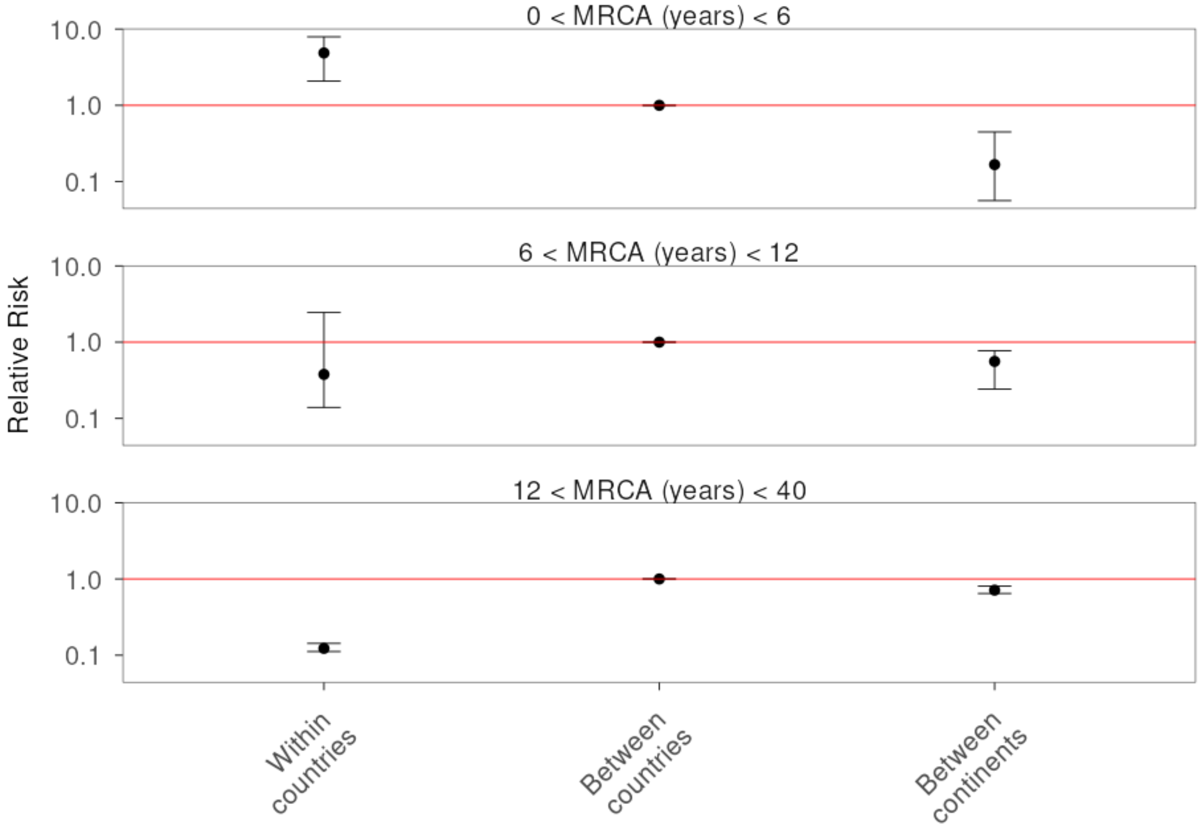


**Fig S8. Phylogeographic analysis of CRAB in Europe.** Relative risk that pairs of isolates were collected in a certain geographic location (same European country, different European countries, different continents where one of them is from Europe) compared to the reference (different European countries). Each facet describes pairs of samples within a certain genetic distance range (time to most recent common ancestor, MRCA, years). Error bars represent 90% CIs based on 100 downsampled time-calibrated trees. In each subsample, 75% of the isolates were drawn from European genomes and 25% from non-European genomes. Some comparisons (e.g. two isolates from the same non-European country) were masked so that risks reflect the situation in Europe but non-European countries are still kept for the between continents geographic category. Data is available in Supplementary Table 8.

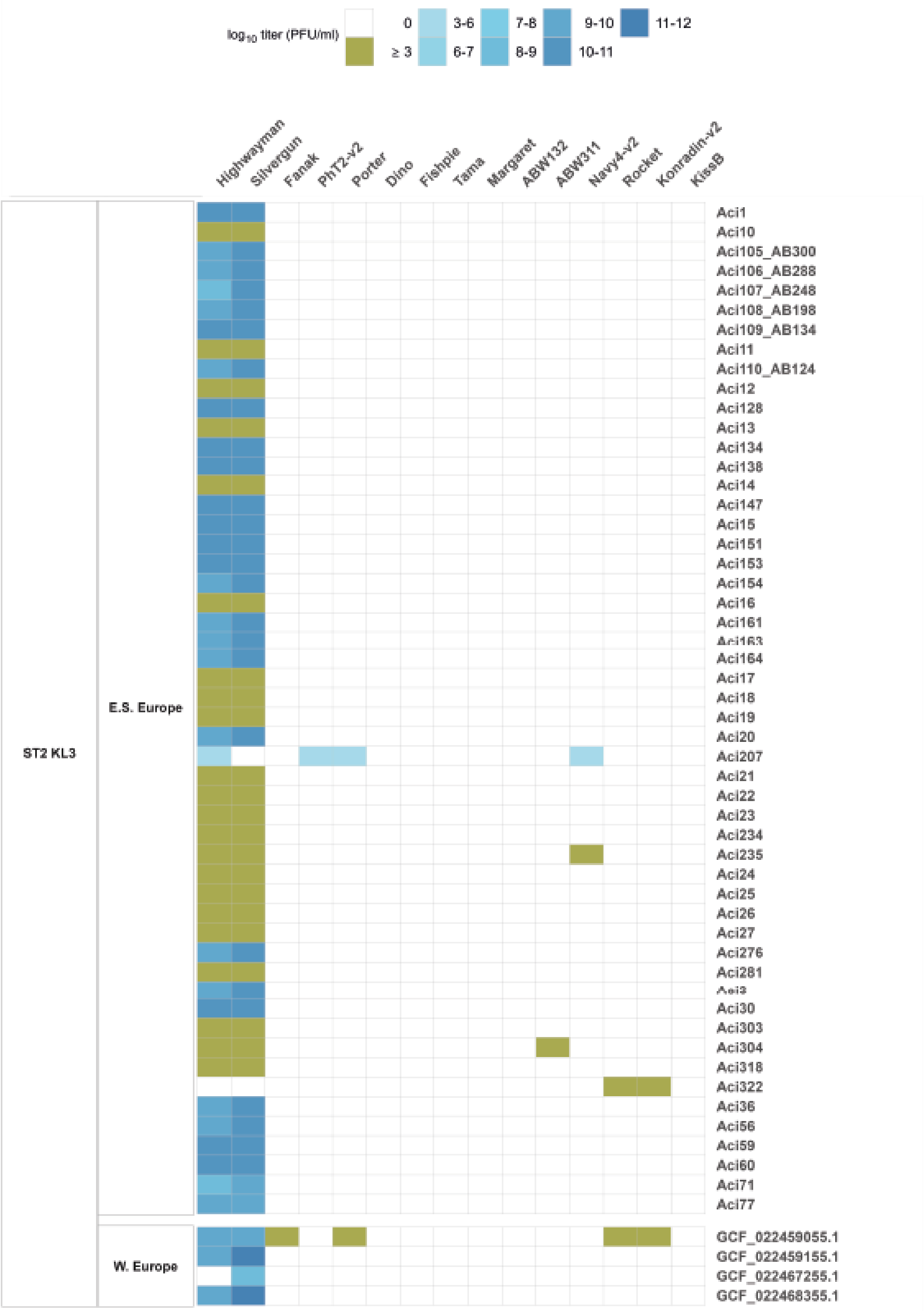

log10 titer (PFU/ml)
0
≥ 3
3-6
6-7
7-8
8-9
9-10
10-11
11-12
Highwayman
Silvergun
Fanak
PhT2-v2
Porter
Dino
Fishpie
Tama
Margaret
ABW132
ABW311
Navy4-v2
Rocket
Konradin-v2
KissB
ST2 KL3
E.S. Europe
W. Europe
Aci1
Aci10
Aci105_AB300
Aci106_AB288
Aci107_AB248
Aci108_AB198
Aci109_AB134
Aci11
Aci110_AB124
Aci12
Aci128
Aci13
Aci134
Aci138
Aci14
Aci147
Aci15
Aci151
Aci153
Aci154
Aci16
Aci161
Aci164
Aci17
Aci18
Aci19
Aci20
Aci207
Aci21
Aci22
Aci23
Aci234
Aci235
Aci24
Aci25
Aci26
Aci27
Aci276
Aci281
Aci30
Aci303
Aci304
Aci318
Aci322
Aci36
Aci56
Aci59
Aci60
Aci71
Aci77
GCF_022459055.1
GCF_022459155.1
GCF_022467255.1
GCF_022468355.1

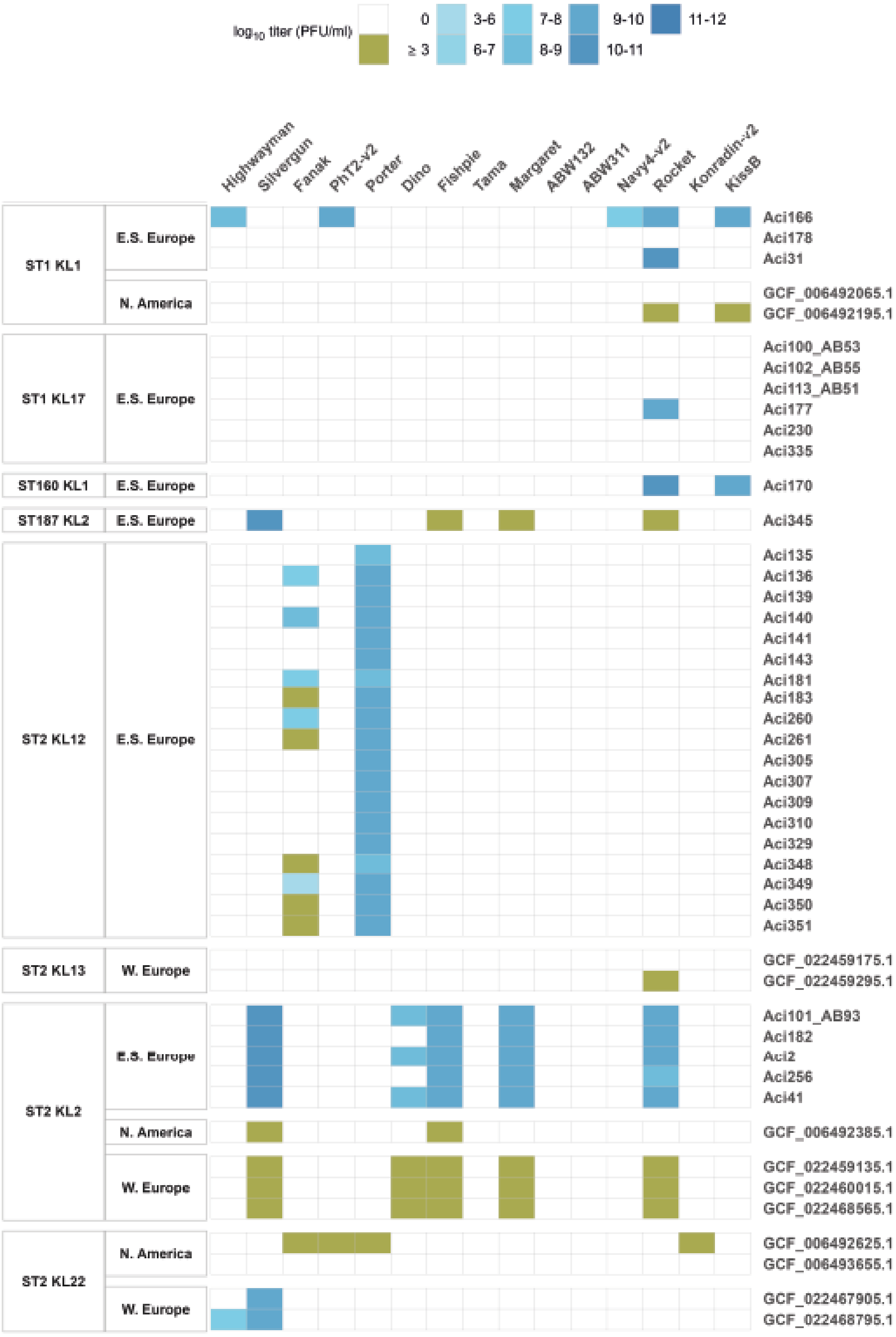
log10 titer (PFU/ml)
0
≥ 3
3-6
6-7
7-8
8-9
9-10
10-11
11-12
Highwayman
Silvergun
Fanak
PhT2-v2
Porter
Dino
Fishpie
Tama
Margaret
ABW132
ABW311
Navy4-v2
Rocket
Konradin-v2
KissB
ST1 KL1
E.S. Europe
N. America
ST1 KL17
ST160 KL1
ST187 KL2
ST2 KL12
ST2 KL13
W. Europe
ST2 KL2
ST2 KL22
Aci166
Aci178
Aci31
GCF_006492065.1
GCF_006492195.1
Aci100_AB53
Aci102_AB55
Aci113_AB51
Aci177
Aci230
Aci335
Aci170
Aci345
Aci135
Aci136
Aci139
Aci140
Aci141
Aci143
Aci181
Aci183
Aci260
Aci261
Aci305
Aci307
Aci309
Aci310
Aci329
Aci348
Aci349
Aci350
Aci351
GCF_022459175.1
GCF_022459295.1
Aci101_AB93
Aci182
Aci2
Aci256
Aci41
GCF_006492385.1
GCF_022459135.1
GCF_022460015.1
GCF_022468565.1
GCF_006492625.1
GCF_006493655.1
GCF_022467905.1
GCF_022468795.1

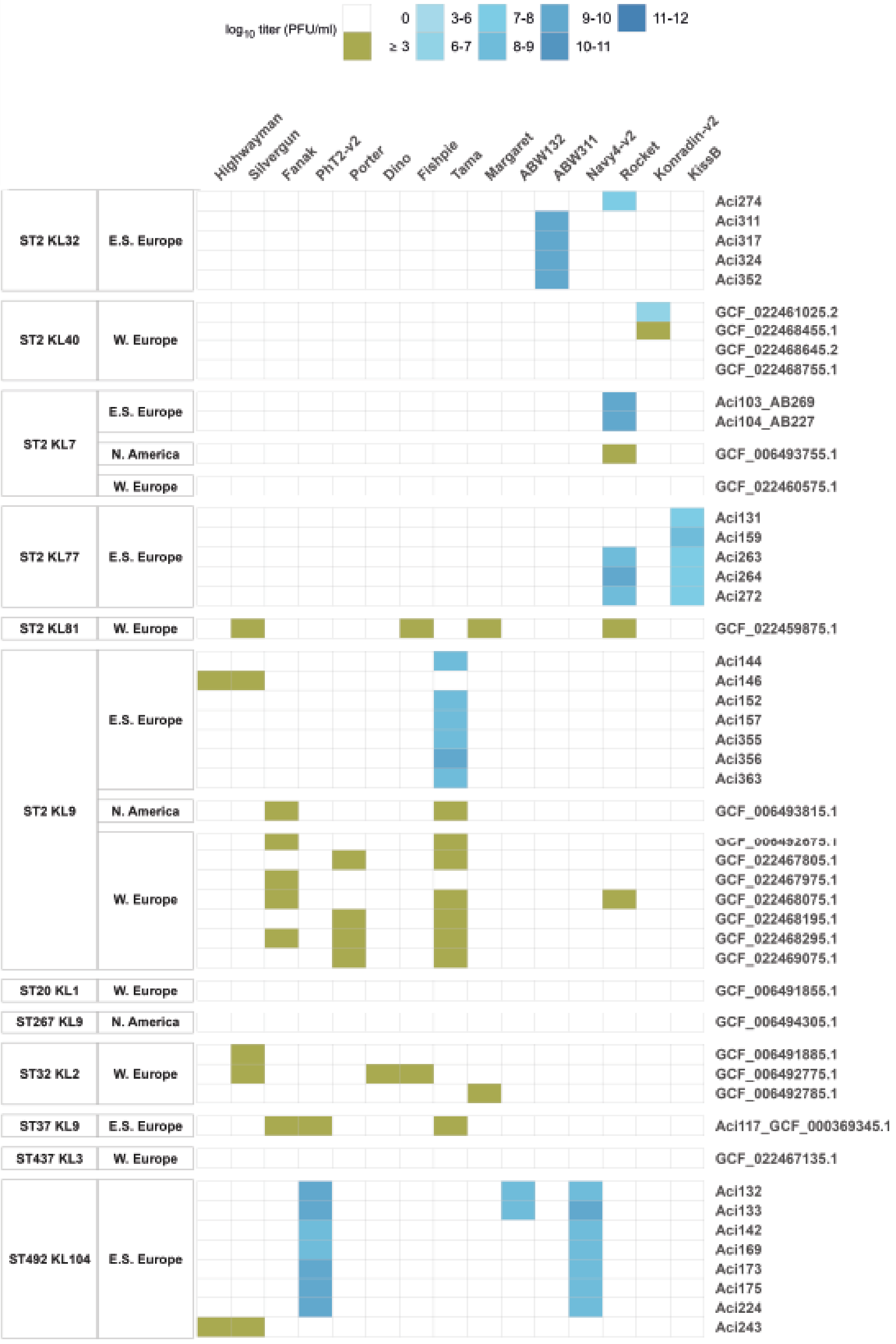

log10 titer (PFU/ml)
0
≥ 3
3-6
6-7
7-8
8-9
9-10
10-11
11-12
Highwayman
Silvergun
Fanak
PhT2-v2
Porter
Dino
Fishpie
Tama
Margaret
ABW132
ABW311
Navy4-v2
Rocket
Konradin-v2
KissB
ST2 KL32
E.S. Europe
Aci274
Aci311
Aci317
Aci324
Aci352
ST2 KL40
W. Europe
GCF_022461025.2
GCF_022468455.1
GCF_022468645.2
GCF_022468755.1
ST2 KL7
E.S. Europe
Aci103_AB269
Aci104_AB227
N. America
GCF_006493755.1
W. Europe
GCF_022460575.1
ST2 KL77
E.S. Europe
Aci131
Aci159
Aci263
Aci264
Aci272
ST2 KL81
W. Europe
GCF_022459875.1
ST2 KL9
E.S. Europe
Aci144
Aci146
Aci152
Aci157
Aci355
Aci356
Aci363
N. America
GCF_006493815.1
W. Europe
GCF_022467805.1
GCF_022467975.1
GCF_022468075.1
GCF_022468195.1
GCF_022468295.1
GCF_022469075.1
ST20 KL1
W. Europe
GCF_006491855.1
ST267 KL9
N. America
GCF_006494305.1
ST32 KL2
W. Europe
GCF_006491885.1
GCF_006492775.1
GCF_006492785.1
ST37 KL9
E.S. Europe
Aci117_GCF_000369345.1
ST437 KL3
W. Europe
GCF_022467135.1
ST492 KL104
E.S. Europe
Aci132
Aci133
Aci142
Aci169
Aci173
Aci175
Aci224
Aci243

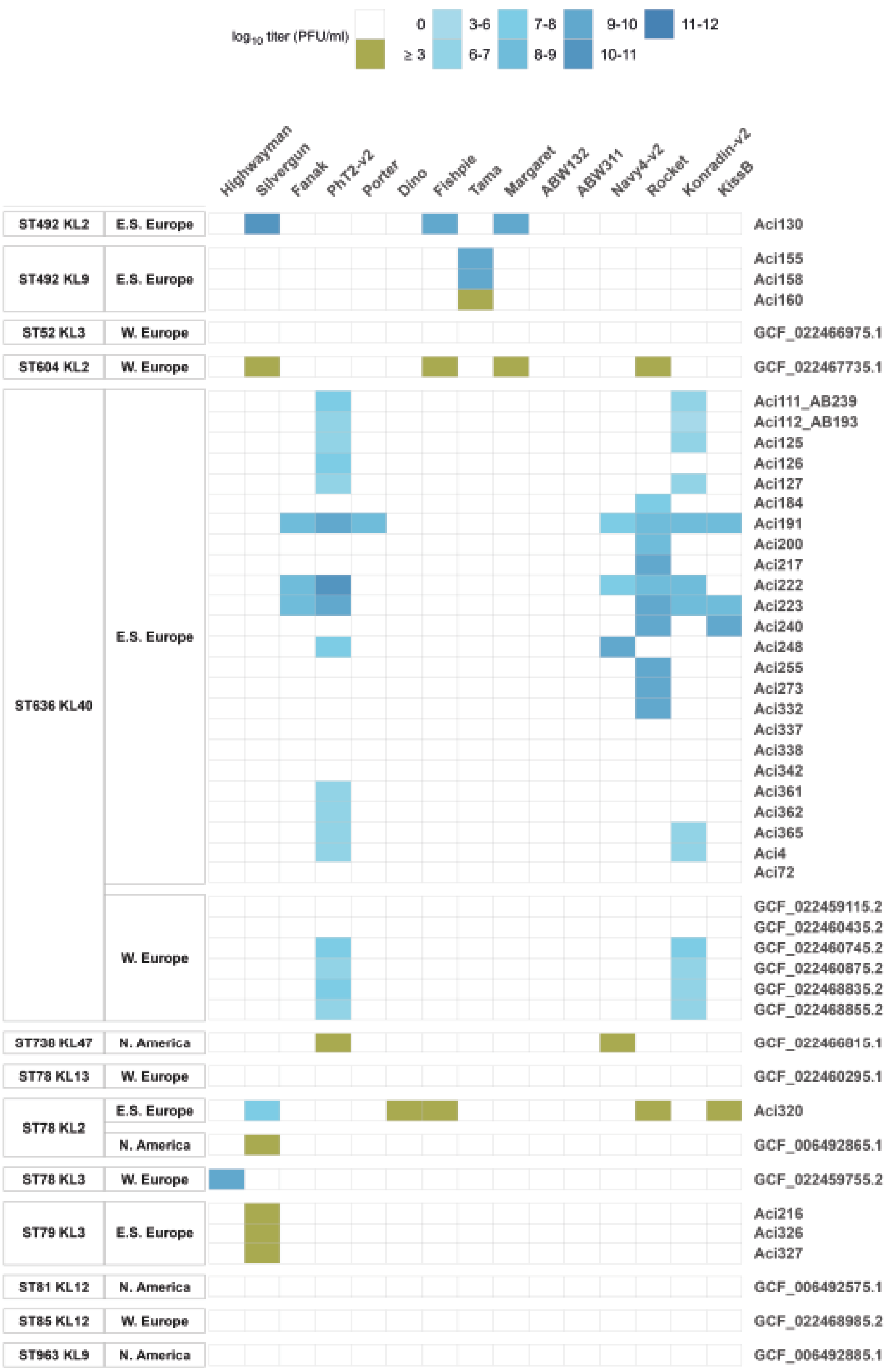
log10 titer (PFU/ml)
0
≥ 3
3-6
6-7
7-8
8-9
9-10
10-11
11-12
Highwayman
Silvergun
Fanak
PhT2-v2
Porter
Dino
Fishpie
Tama
Margaret
ABW132
ABW311
Navy4-v2
Rocket
Konradin-v2
KissB
ST492 KL2
E.S. Europe
Aci130
ST492 KL9
E.S. Europe
Aci155
Aci158
Aci160
ST52 KL3
W. Europe
GCF_022466975.1
ST604 KL2
W. Europe
GCF_022467735.1
ST636 KL40
E.S. Europe
Aci111_AB239
Aci112_AB193
Aci125
Aci126
Aci127
Aci184
Aci191
Aci200
Aci217
Aci222
Aci223
Aci240
Aci248
Aci255
Aci273
Aci332
Aci337
Aci338
Aci342
Aci361
Aci362
Aci365
Aci4
Aci72
W. Europe
GCF_022459115.2
GCF_022460435.2
GCF_022460745.2
GCF_022460875.2
GCF_022468835.2
GCF_022468855.2
ST730 KL47
N. America
GCF_022466815.1
ST78 KL13
W. Europe
GCF_022460295.1
ST78 KL2
E.S. Europe
Aci320
N. America
GCF_006492865.1
ST78 KL3
W. Europe
GCF_022459755.2
ST79 KL3
E.S. Europe
Aci216
Aci326
Aci327
ST81 KL12
N. America
GCF_006492575.1
ST85 KL12
W. Europe
GCF_022468985.2
ST963 KL9
N. America
GCF_006492885.1

**Fig S9. Sensitivity profile of 199 *A. baumannii* isolates (y axis) against the 15 isolated phages (x axis).** Isolates are grouped based on the MLST-CPS type they belong to and their geographic origin. The intensity of the colour blue indicates the mean (n = 3) of the $\log_{10}$ phage titer (PFU/ml) (the higher the titer, the darker the intensity). Green colour (≥ 3) indicates that the exact titer was not measured, but it was determined with spot assay that the given phage can infect the given isolate. Data is available in Supplementary Table 10. Abbreviations: E.S. Europe - Eastern and Southern Europe, W. Europe - Western Europe, N. America - North America.

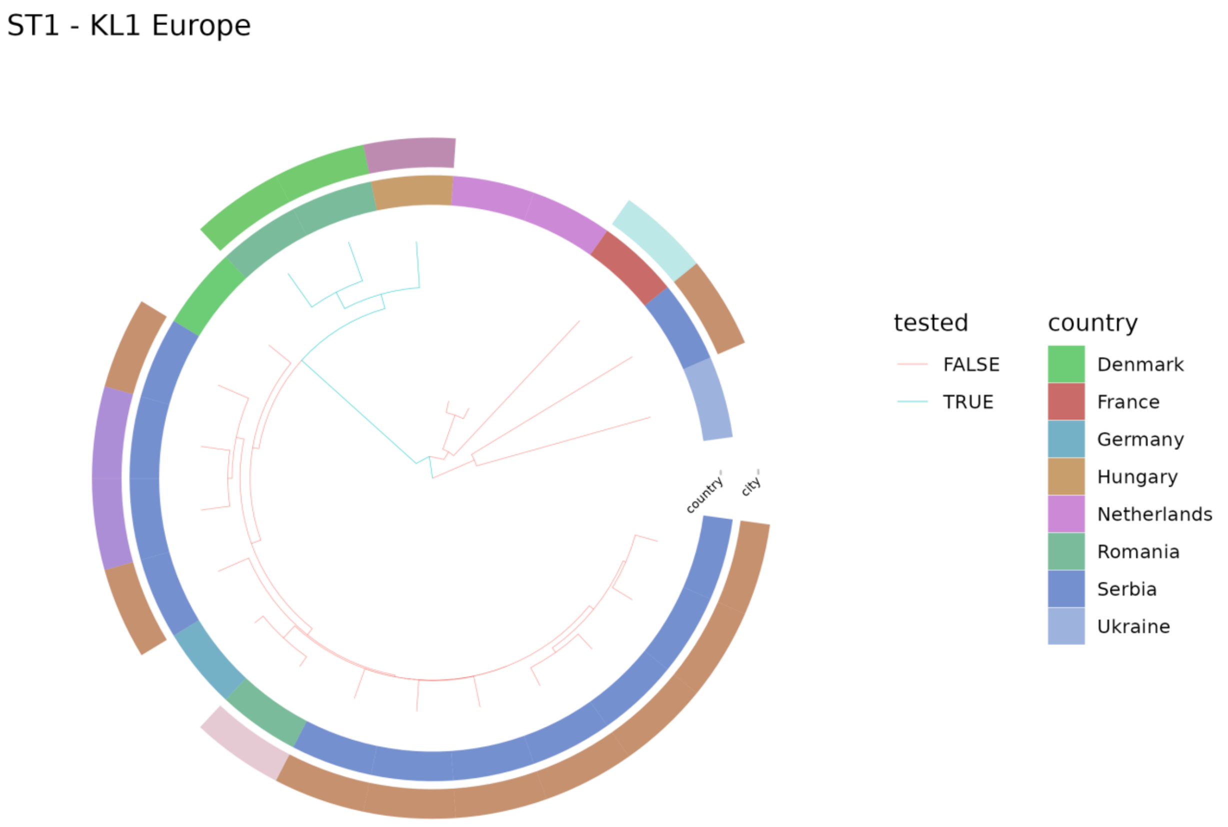


**Fig S10A. Phylogenetic tree of 22 European ST1-KL1 strains.** The phylogenetic tree was created using all 566 ST1 genomes from the 11,129 good quality (Methods, Filtering assemblies) *Acinetobacter baumannii* whole genome sequences included in this study. The ST1 tree was then filtered to ST1-KL1 European genomes for plotting.

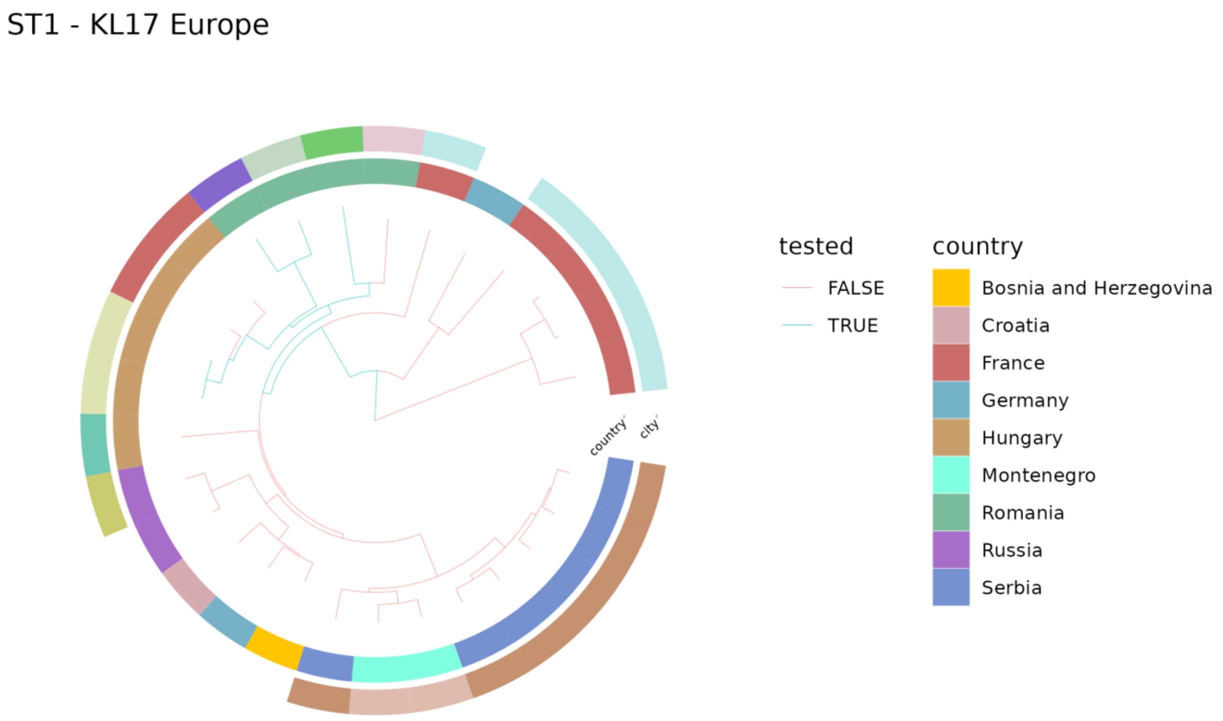


**Fig S10B. Phylogenetic tree of 28 European ST1-KL17 strains.** The phylogenetic tree was created using all 566 ST1 genomes from the 11,129 good quality (Methods, Filtering assemblies) *Acinetobacter baumannii* whole genome sequences included in this study. The ST1 tree was then filtered to ST1-KL17 European genomes for plotting.

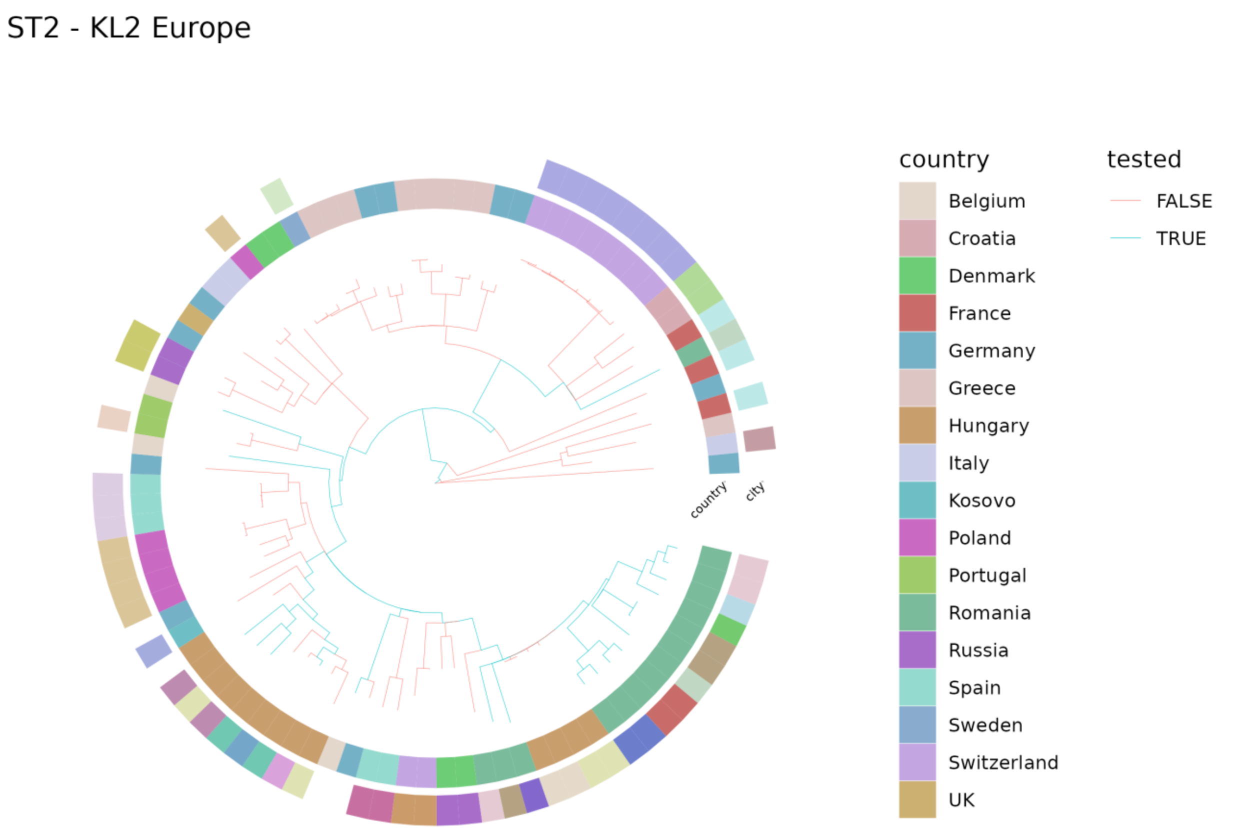


**Fig S10C. Phylogenetic tree of 90 European ST2-KL2 strains.** The phylogenetic tree was created using all 7,735 ST2 genomes from the 11,129 good quality (Methods, Filtering assemblies) *Acinetobacter baumannii* whole genome sequences included in this study. The ST2 tree was then filtered to ST2-KL2 European genomes for plotting.

#### ST2 - KL3 Europe

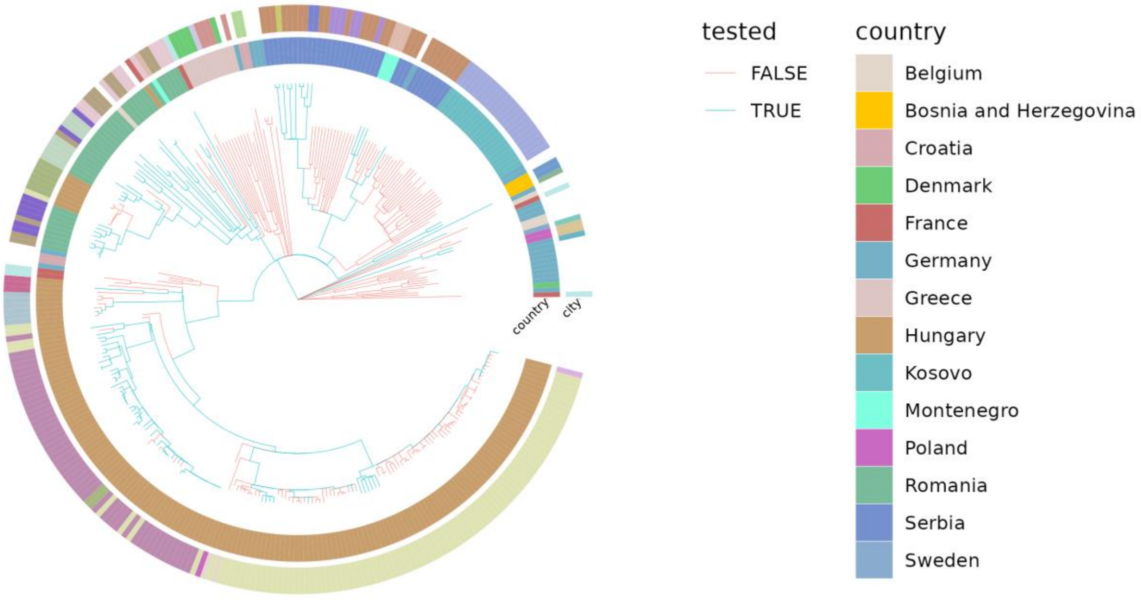


**Fig S10D. Phylogenetic tree of 316 European ST2-KL3 strains.** The phylogenetic tree was created using all 7,735 ST2 genomes from the 11,129 good quality (Methods, Filtering assemblies) *Acinetobacter baumannii* whole genome sequences included in this study. The ST2 tree was then filtered to ST2-KL3 European genomes for plotting.

ST2 - KL7 Europe

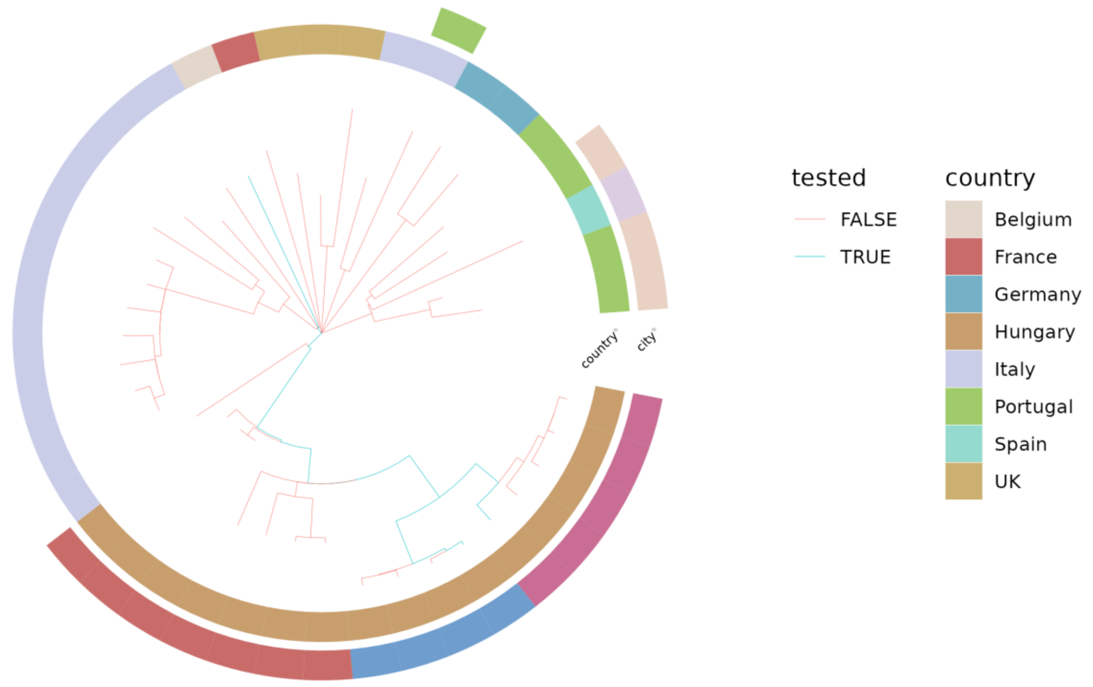


**Fig S10E. Phylogenetic tree of 42 European ST2-KL7 strains.** The phylogenetic tree was created using all 7,735 ST2 genomes from the 11,129 good quality (Methods, Filtering assemblies) *Acinetobacter baumannii* whole genome sequences included in this study. The ST2 tree was then filtered to ST2-KL7 European genomes for plotting.

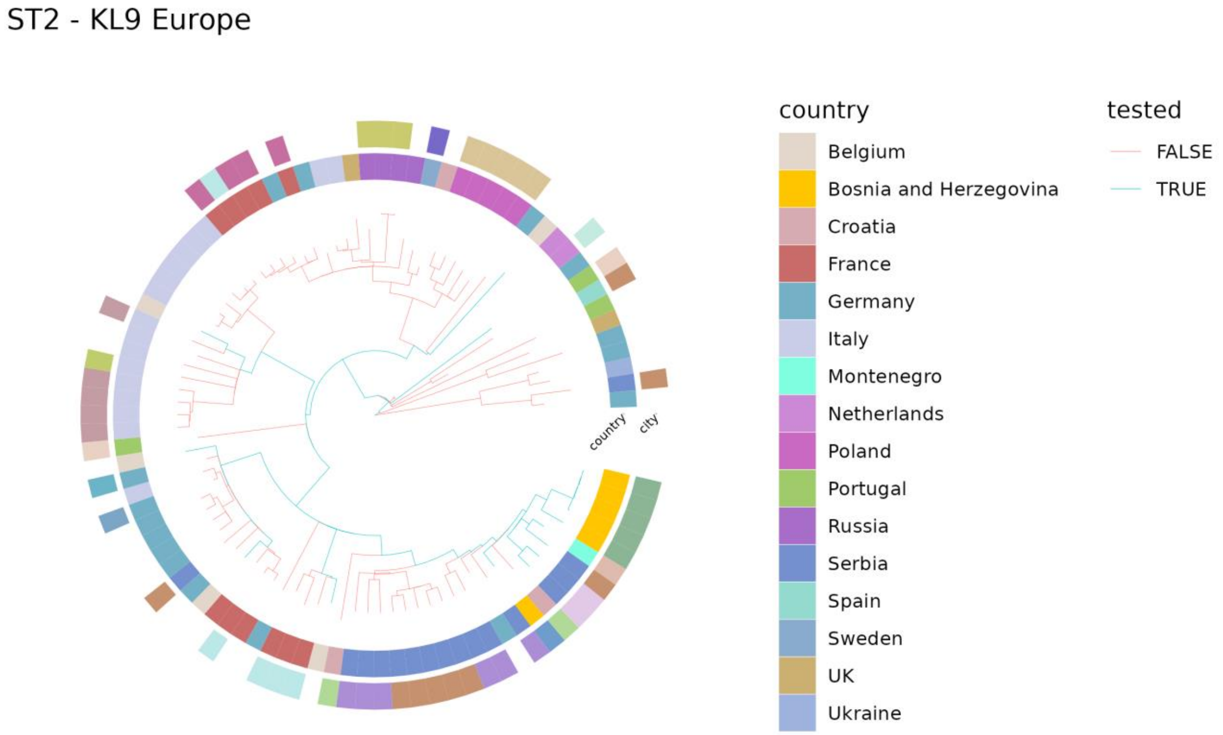


**Fig S10F. Phylogenetic tree of 94 European ST2-KL9 strains.** The phylogenetic tree was created using all 7,735 ST2 genomes from the 11,129 good quality (Methods, Filtering assemblies) *Acinetobacter baumannii* whole genome sequences included in this study. The ST2 tree was then filtered to ST2-KL9 European genomes for plotting.

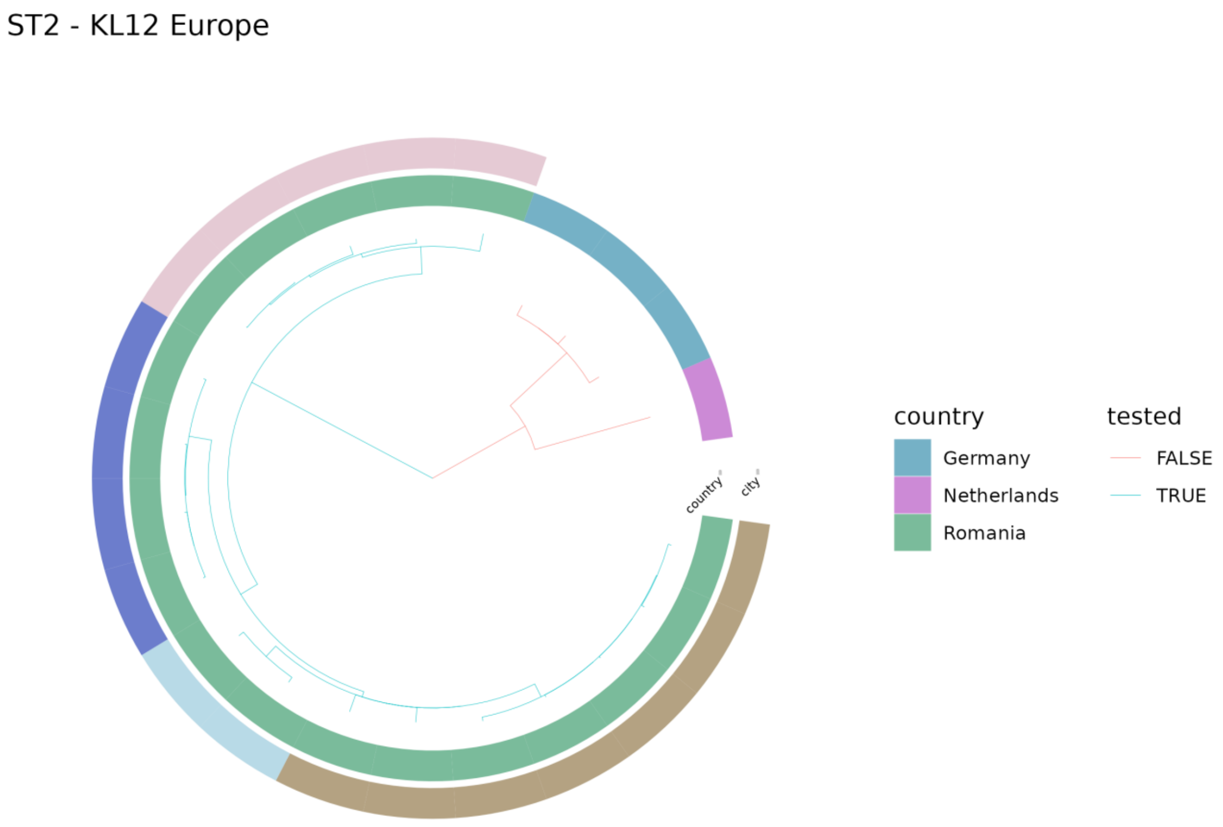


**Fig S10G. Phylogenetic tree of 22 European ST2-KL12 strains.** The phylogenetic tree was created using all 7,735 ST2 genomes from the 11,129 good quality (Methods, Filtering assemblies) *Acinetobacter baumannii* whole genome sequences included in this study. The ST2 tree was then filtered to ST2-KL12 European genomes for plotting.

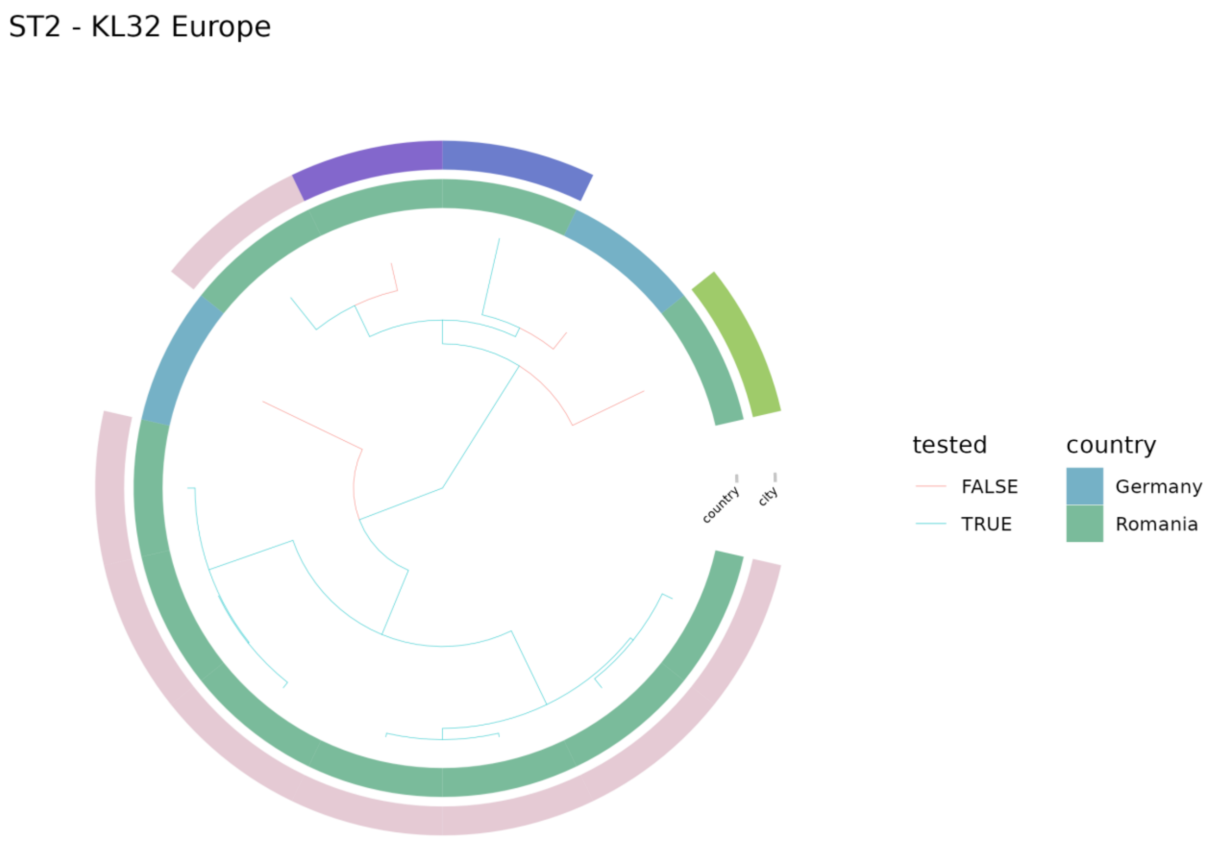


**Fig S10H. Phylogenetic tree of 13 European ST2-KL32 strains.** The phylogenetic tree was created using all 7,735 ST2 genomes from the 11,129 good quality (Methods, Filtering assemblies) *Acinetobacter baumannii* whole genome sequences included in this study. The ST2 tree was then filtered to ST2-KL32 European genomes for plotting.

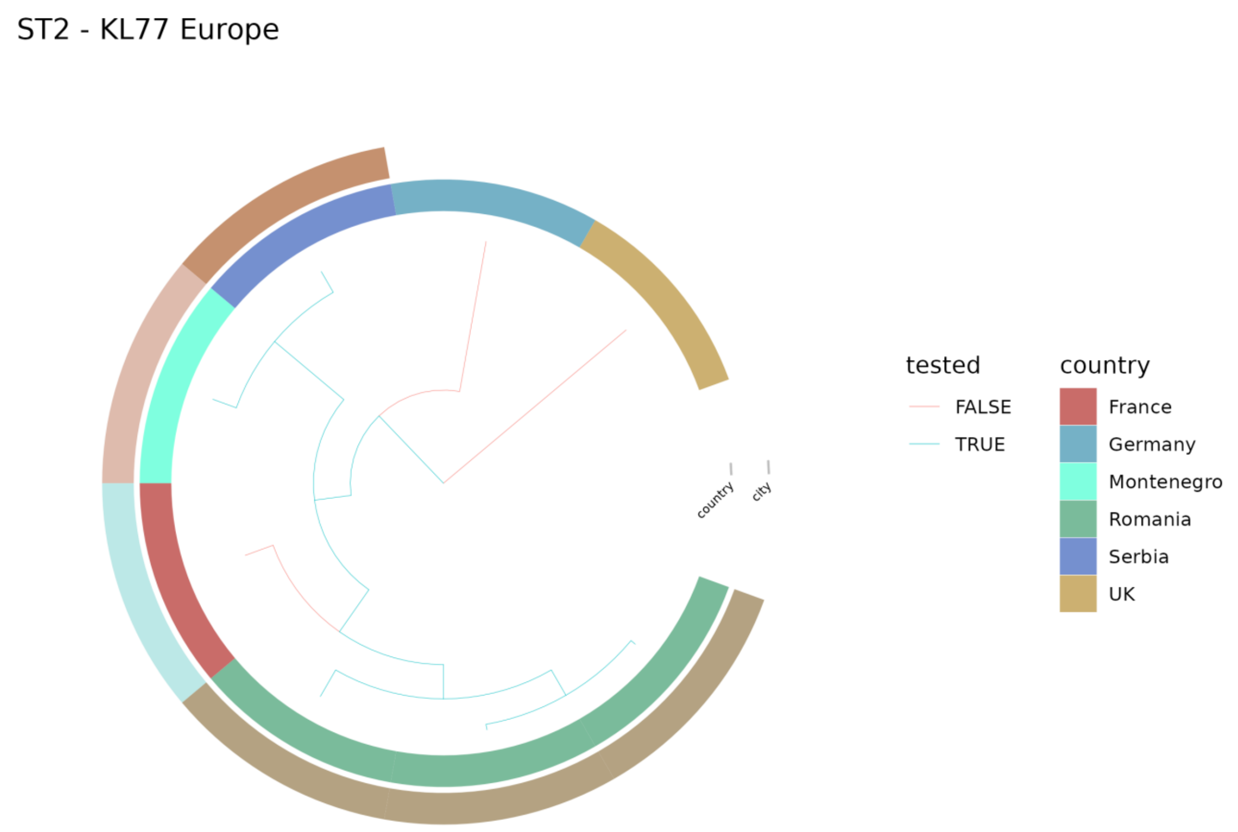


**Fig S10I. Phylogenetic tree of 8 European ST2-KL77 strains.** The phylogenetic tree was created using all 7,735 ST2 genomes from the 11,129 good quality (Methods, Filtering assemblies) *Acinetobacter baumannii* whole genome sequences included in this study. The ST2 tree was then filtered to ST2-KL77 European genomes for plotting.

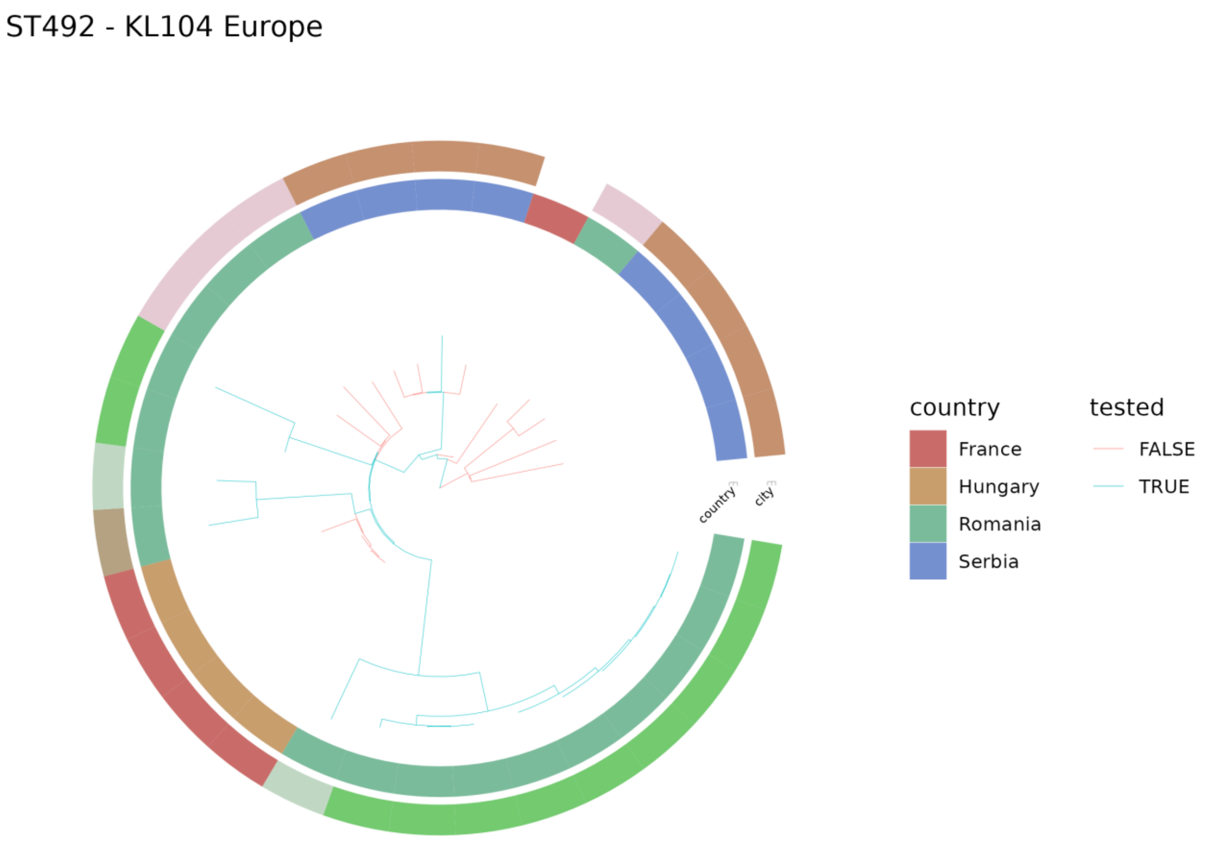


**Fig S10J. Phylogenetic tree of 31 European ST492-KL104 strains.** The phylogenetic tree was created using all 42 ST492 genomes from the 11,129 good quality (Methods, Filtering assemblies) *Acinetobacter baumannii* whole genome sequences included in this study. The ST492 tree was then filtered to ST492-KL104 European genomes for plotting.

ST636 - KL40 Europe

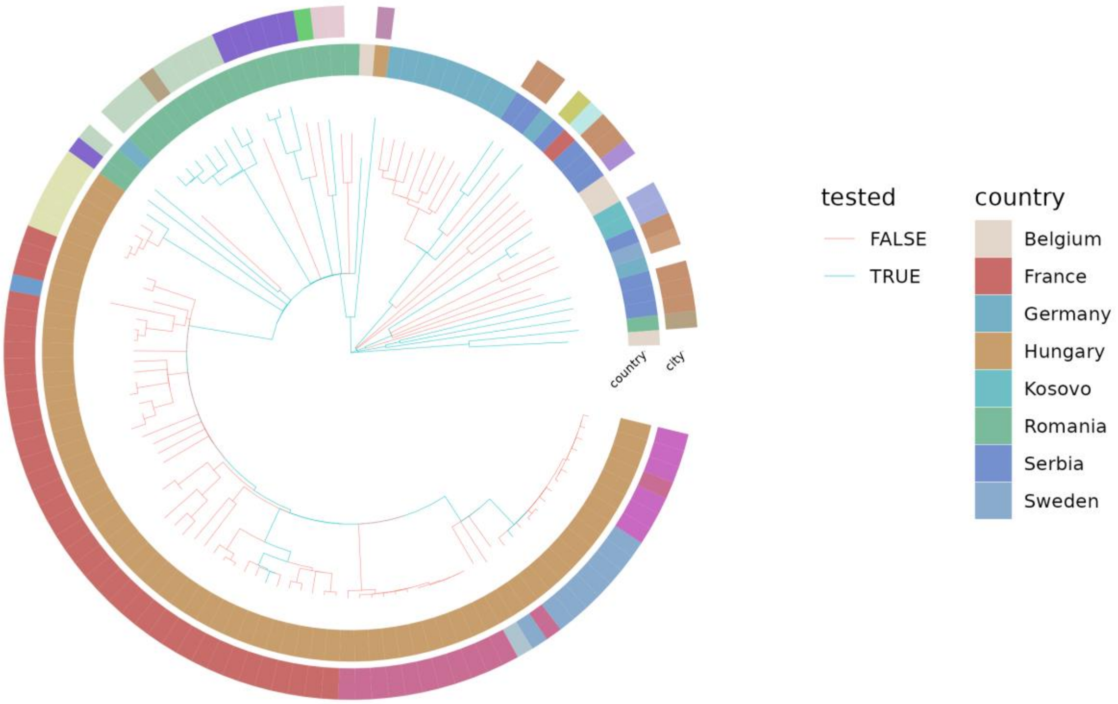


**Fig S10K. Phylogenetic tree of 123 European ST636-KL40 strains.** The phylogenetic tree was created using all 141 ST636 genomes from the 11,129 good quality (Methods, Filtering assemblies) *Acinetobacter baumannii* whole genome sequences included in this study. The ST636 tree was then filtered to ST636-KL40 European genomes for plotting. Please note that the 11,129 genomes do not include any other CPS types for ST636. Therefore, all the ST636 genomes that are not on this plot are also ST636-KL40 genomes but from another continent (Asia).

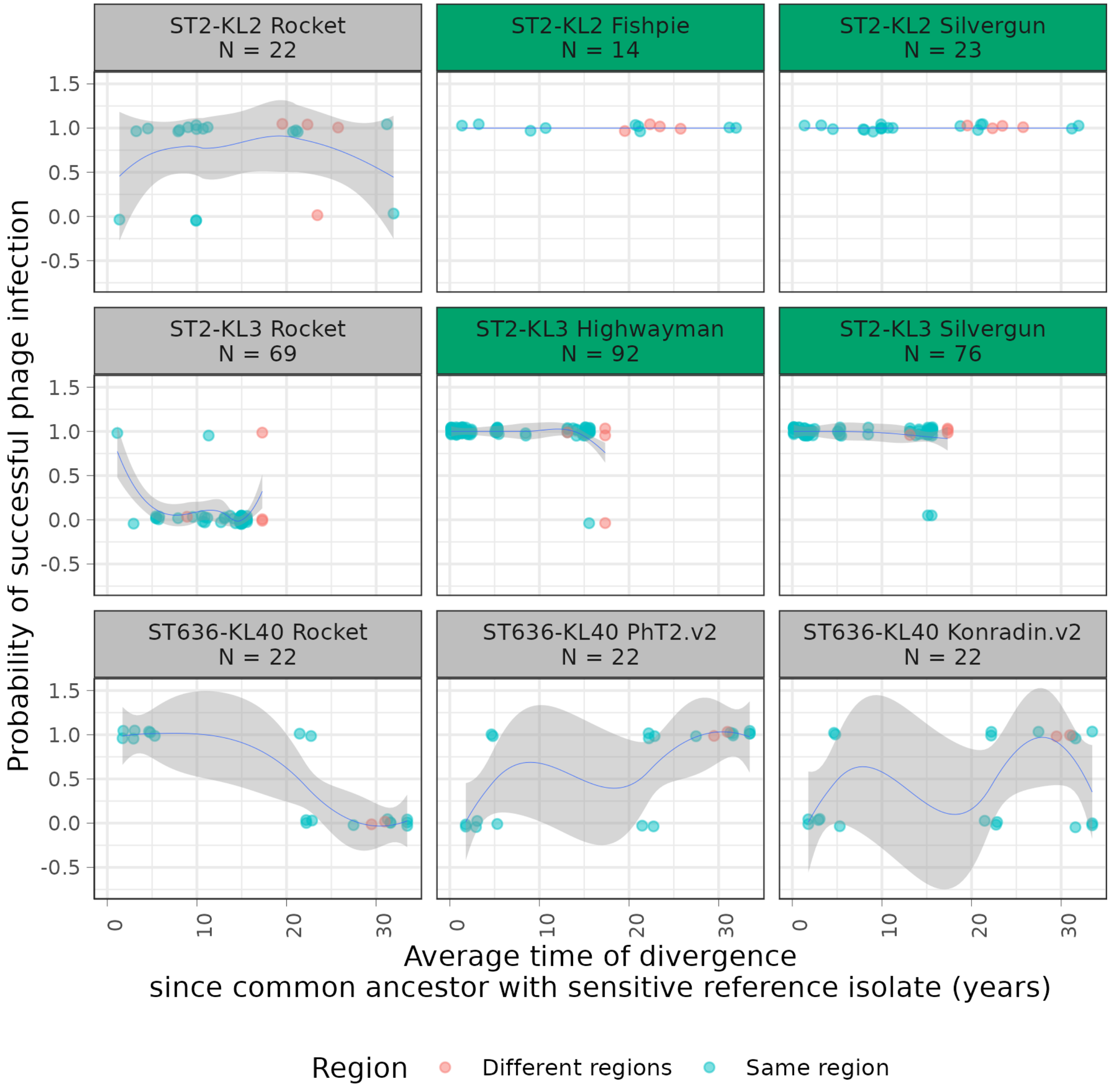


**Fig S11. Specific phages remain effective across most isolates of the same MLST-CPS type regardless of genetic divergence.** Each panel shows the effectiveness of a phage against *A. baumannii* isolates that belong to an MLST-CPS type, over various times of divergence. The x-axis shows the divergence between an isolate and a reference isolate, measured as the average distance from the most recent common ancestor of the isolate and the reference isolate, using the respective dated phylogenetic tree (Methods). The reference isolate was selected separately for each MLST-CPS type - phage combination and had the following properties: it (i) belonged to the same MLST-CPS type, (ii) was sensitive to the phage, and (iii) maximised the range of covered phylogenetic distances. The y-axis is 0 if the isolate was resistant to the phage and 1 if it was sensitive. N indicates the number of samples in each panel. Since one sample is always the reference, each panel contains N-1 data points. Data point colours indicate whether a sample was isolated in the same world region as the reference sample. The fitted line is smoothed by a loess method. Ideally, a successful phage should infect all isolates in an MLST-CPS category and the covered distance range should be large. Panels highlighted in green indicate promising MLST-CPS-phage combinations. For example, Silvergun could infect all studied ST2-KL2 isolates and these isolates span more than 30 years of genome divergence. Likewise, Highwayman could infect all but a few studied ST2-KL3 isolates and these isolates span 15-20 years. On the other hand, the effectiveness of studied phages against ST636-KL40 isolates was sporadic, indicating that we could not find an ideal phage against ST636-KL40.

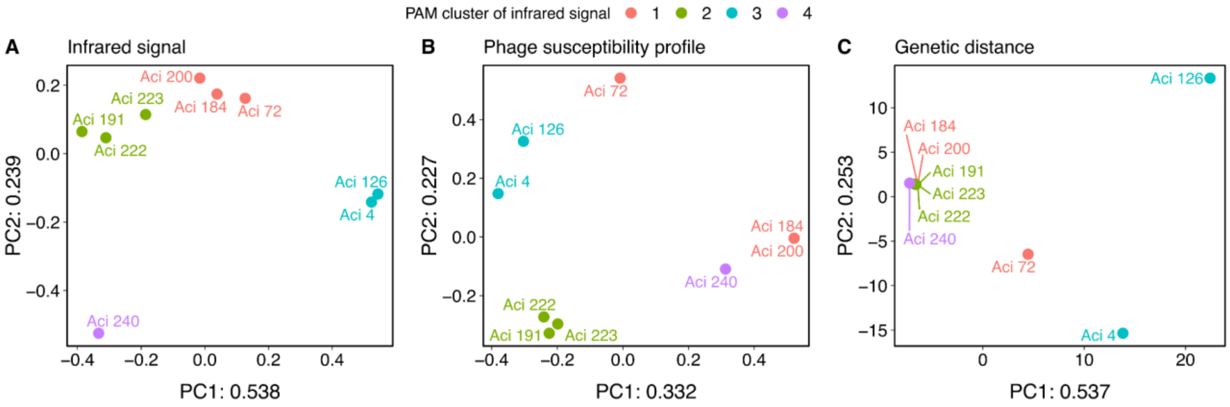


**Fig S12. Differences in cell surface properties between recently diverged isolates of ST636-KL40 strain correlate with phage susceptibility profile differences. A,** Principal coordinate analysis (PCoA) plot of nine ST636-KL40 isolates from Eastern and Southern Europe derived from the Fourier-transformed infrared measurement using the IR Biotyper instrument (Supplementary Table 23, Methods). Each isolate has at least one pair of isolates that are phylogenetically closely related (that is, diverged within two years). Principal coordinates 1 and 2 explain 53% and 24% of the variation. The isolates are coloured based on their Partitioning Around Medoids (PAM) clustering into four clusters. **B,** PCoA plot of the phage susceptibility profile differences calculated with the Jaccard index. Principal coordinates 1 and 2 explain 33% and 23% of the variation. The isolates are coloured based on the PAM clustering into four clusters of the samples in Fig S12A. Differences in the surface properties derived from the Fourier-transformed infrared signal correlate with phage susceptibility profile differences calculated with the Jaccard index (Mantel test, $r = 0.408$, $p$-value = 0.011, n = 9, number of permutations: 10,000, method: Pearson correlation). Measurements were carried out in three technical replicates. **C,** PCoA plot of the genetic distances for the isolates. Principal coordinates 1 and 2 explain 53% and 25% of the variation. The isolates are coloured based on the PAM clustering into four clusters of the samples in Fig S12A. Correlation cannot be observed between the Fourier-transformed infrared signal and the genetic distances for the isolates (Mantel test, $r = -0.025$, $p$-value = 0.499, n = 9, number of permutations: 10,000, method: Pearson correlation). The PCoA analyses were carried out with the ape R package.[3] The PAM clustering was calculated with the cluster R package.[4] The Jaccard distances were calculated using the ade4 R package.[5] The Mantel tests were carried out with the vegan R package.[6]

**A**

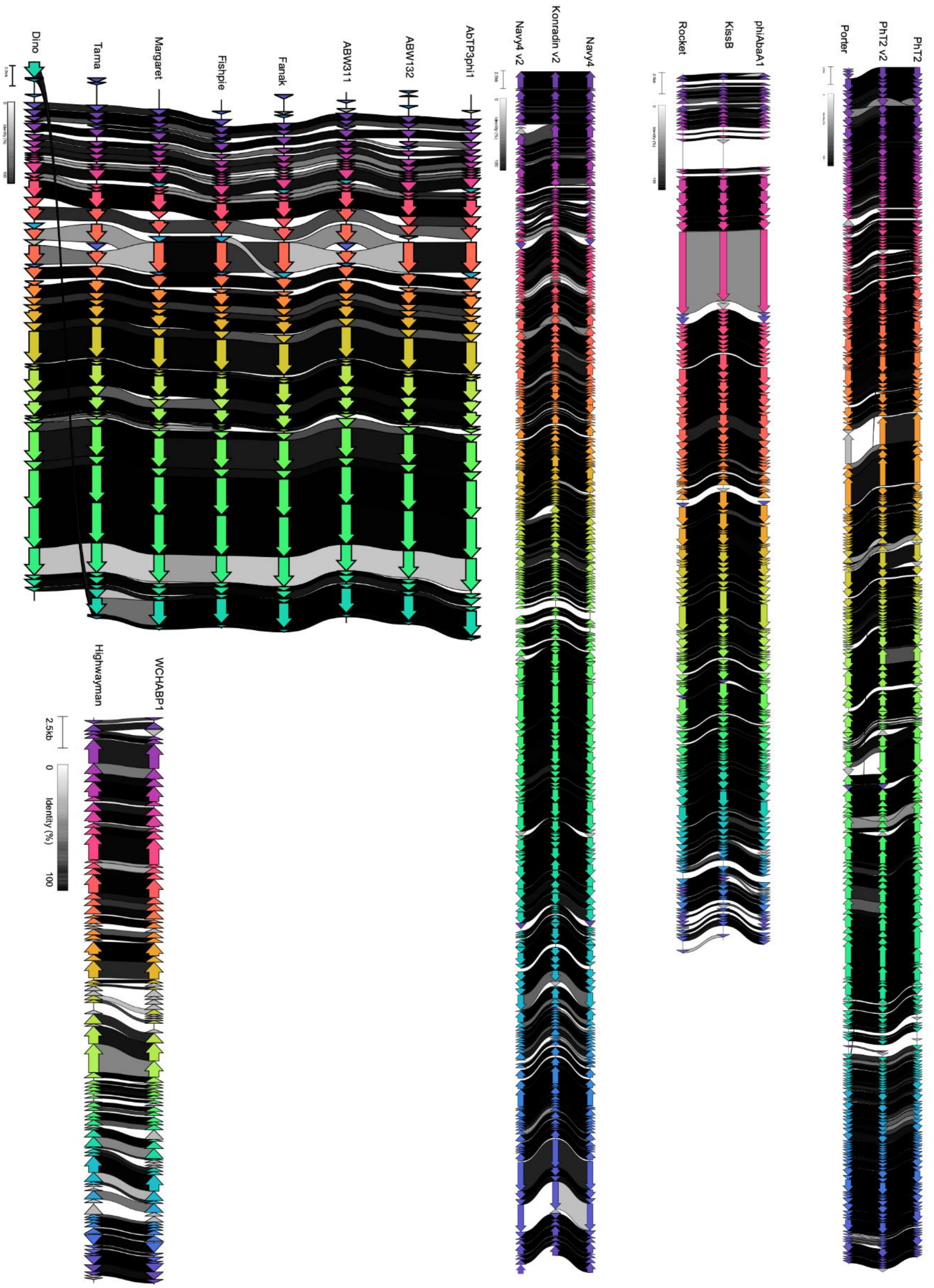
Dino
Tama
Margaret
Fishpie
Fanak
ABW311
ABW132
AbTP3phi1
Navy4 v2
Konradin v2
Navy4
Rocket
KissB
phiAbaA1
Porter
PhT2 v2
PhT2
Highwayman
WCHABP1
2.5kb
0
Identity (%)
100

1/2

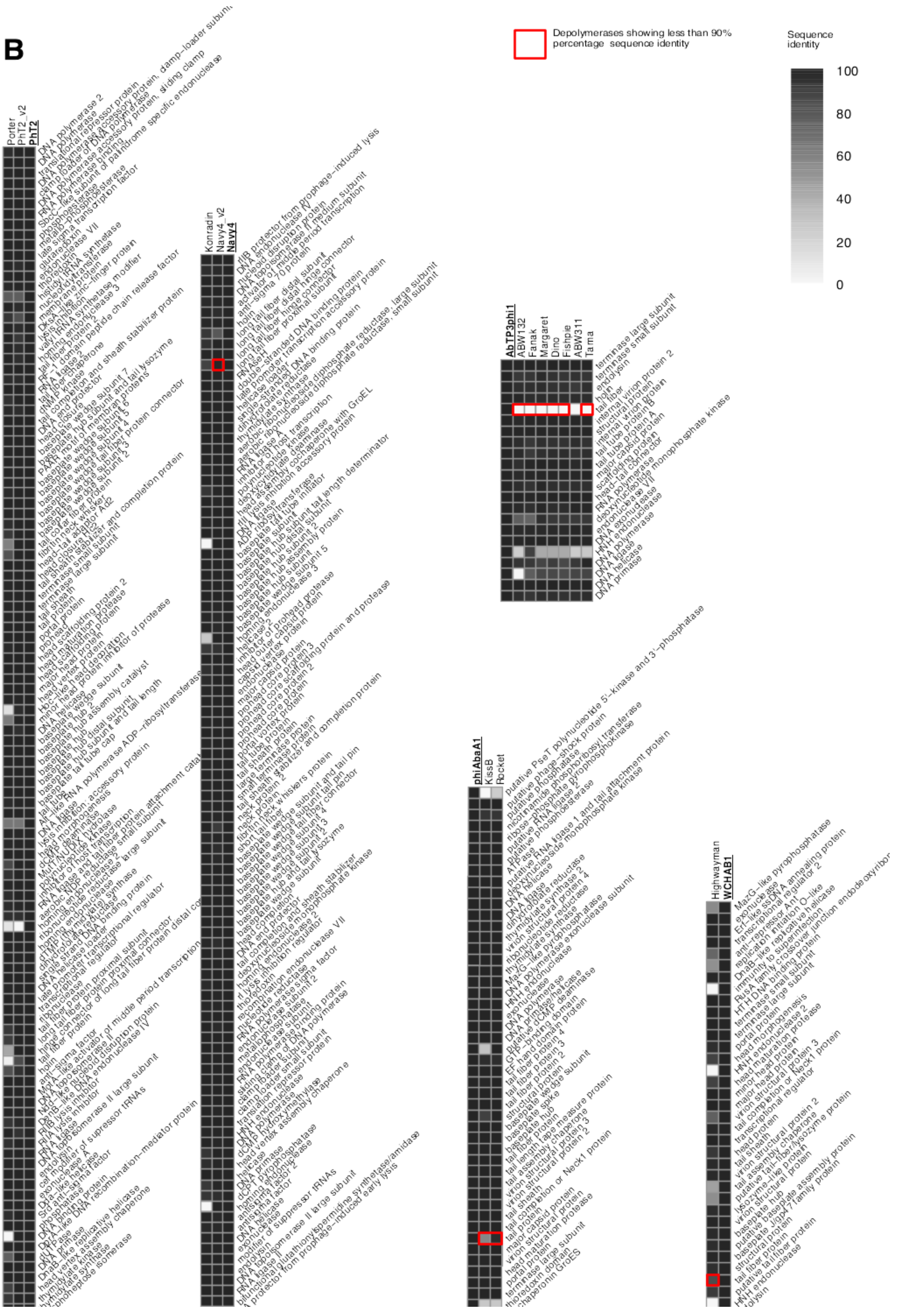
B
Depolymerases showing less than 90% percentage sequence identity
Sequence identity
100
80
60
40
20
0
Porter
PhT2_v2
PhT2
Konradin
Navy4_v2
Navy4
AbTP3phi1
ABW132
Fanak
Margaret
Dino
Fishpie
ABW311
Tama
phiAbaA1
KissB
Rocket
Highwayman
WCHAB1

**Fig S13. Comparing genome architectures and protein sequence identifies between the discovered 14 phages and their closest lytic or therapeutic relatives. A,** Genome architecture similarities as represented by Clinker plots for the 14 phages that show high similarity to other known lytic or therapeutic phages. These reference lytic or therapeutic phages are shown alongside the discovered phages for each group. The connecting lanes are shaded based on the degree of similarity between the ORFs from 0 to 100% percentage identity on the DNA level. As Silvergun does not have a known close relative, it is not shown on this figure **B,** Pairwise protein sequence identities of shared orthogroups for the 14 discovered phages and for their closest lytic or therapeutic relatives. The shading in the heatmap's cells represents the degree of similarity between the translated ORFs from 0 to 100% percentage identity.

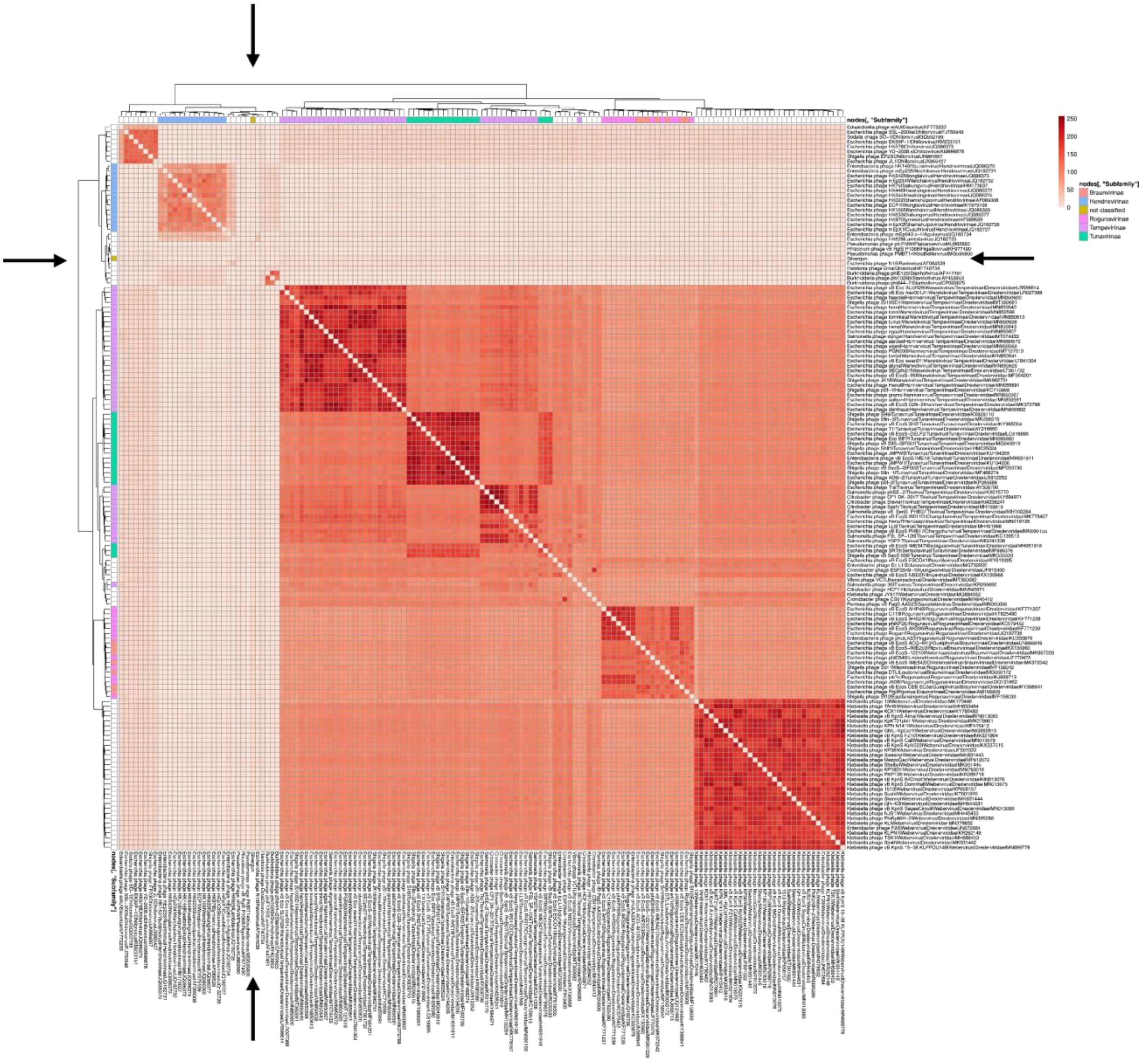


**Fig S14. Number of shared orthogroups between Silvergun and known phage species in its vContact network cluster.** Heatmap showing the Weight of all nodes connected to Silvergun in the vConTACT2 network (Fig 5B). This metric emphasises the number of shared orthogroups between two genomes - nodes in the network. Arrows indicate the position of Silvergun on the heatmap. Data is available in Supplementary Table 13-14.

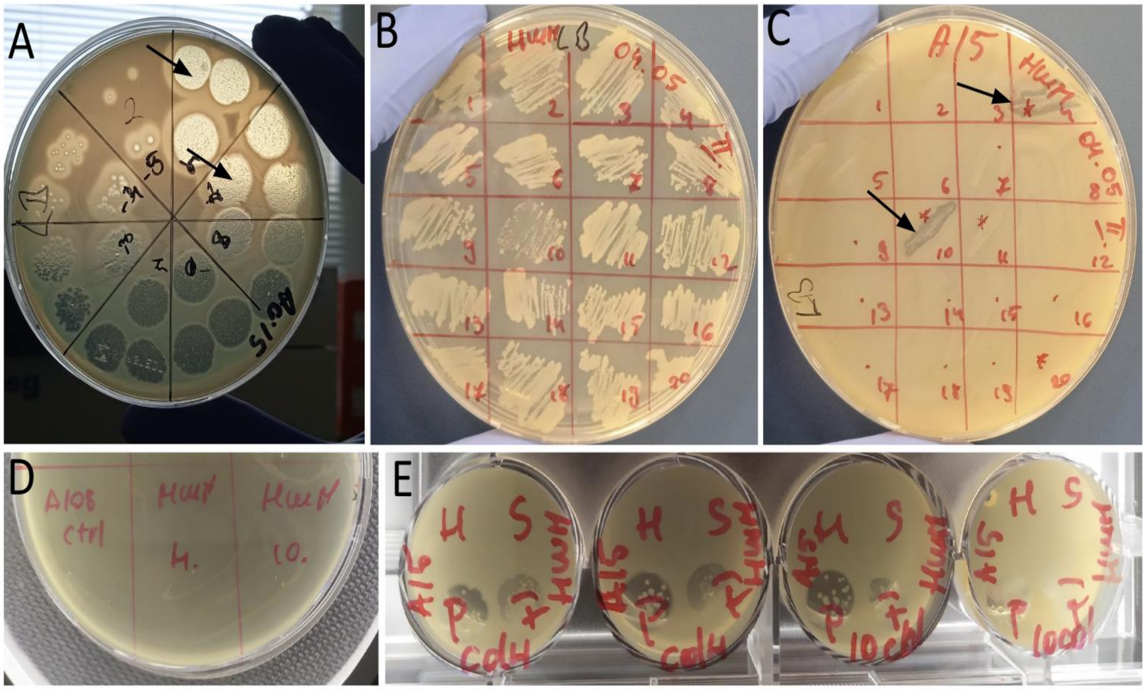


**Fig S15.1 Testing Highwayman for lysogenic lifecycle on one of the ST2-KL3 isolates (Aci15). A,** Spot assay with Highwayman in serial dilution after 24-hours incubation, spots are resulted from 10ul droplets, black arrows denote the location of mesas. **B-C**. Second round of patch assay. Identical colonies from the previous patch screen were plated on LB plate and LB with bacterial overlay. Arrows indicate the bacterial growth in the cells 4 and 10 with surrounding lysis, indicating the presence of the phage. **D**. Spot assay performed with the filtered supernatant from cultures of single colonies after the third round of purification of the colonies 4 and 10. As a control, we used the supernatant of a different bacteria. **E**. Phage susceptibility test, indicates the altered phage infection profile. Each test was done in duplicate. Cells show no plaques in the cases of H and S but they became sensitive to F and Ph.

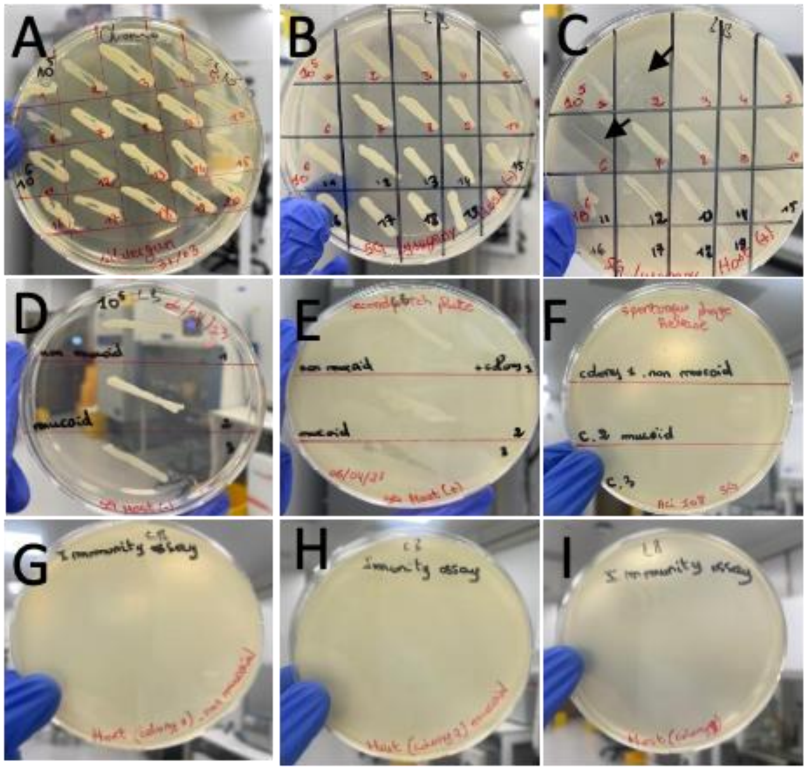


**Fig S15.2 Testing Silvergun for lysogenic lifecycle on one of the ST2-KL3 isolates (Aci108). A,** Streak plate of mesas formed at $10^5$ and $10^6$ PFU/ml phage droplets (10 colonies were screened for each mesas). **B-C,** First, patch using plates with and without lawns of the bacterial hosts. Black arrow (colony 2 and 6) show bacterial growth with surrounding lysis, indicating the presence of phage, possibly released from a lysogen. **D-E**, second patch plate, after three rounds of single colony purification. Colony 2 exhibited two different phenotypes mucoid and non-mucoid. Both of them were screened separately, without indication of phage presence when patched onto plates containing bacterial host lawns. However, for colony 6 (labelled 3 on the plate), there was a lysis zone surrounding it, probably due to phage carried over from the streak plate. **F,** Spot assay performed with the filtered supernatant from cultures of the putative lysogenic colonies, with no indication of spontaneous phage release. **G-I,** Immunity assay of colonies 1,2, and 3 indicates absence of lysis zones. The test demonstrates resistance to the phage, most likely due to a mechanism other than homoimmunity.

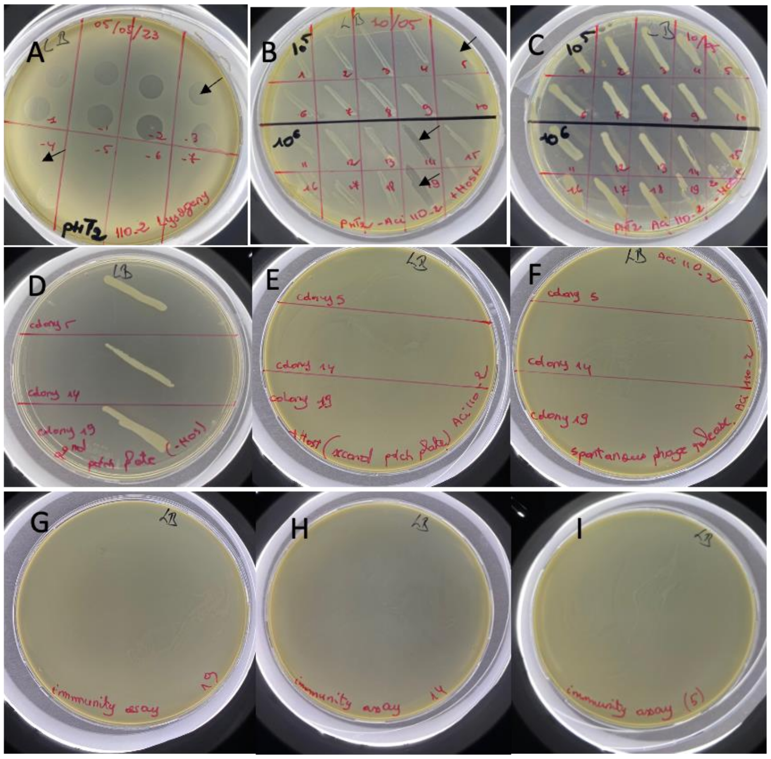


**Fig S15.3 Testing PhT2-v2 for lysogenic lifecycle on one of the ST2-KL3 derivative (A110-2) that is resistant to Highwayman and Silvergun. A,** Spot assay with PhT2-v2 in serial dilution after 24-hours incubation, spots are resulted from 10ul droplets, black arrows denote the location of mesas. **B-C**, First patch using plates with and without lawns of the bacterial hosts. Black arrows (colony 5,14, and 19) show bacterial growth with surrounding lysis, indicating the presence of phage, possibly released from a lysogen. **D-E,** second patch plate, after three rounds of single colony purification without indication of phage presence when patched onto plates containing bacterial host lawns. **F,** Spot assay performed with the filtered supernatant from cultures of the putative lysogenic colonies, with no indication of spontaneous phage release. **G-I,** Immunity assay of colonies 5,14, and 19 indicates absence of lysis zones. The test demonstrates resistance to the phage, most likely due to a mechanism other than homoimmunity.

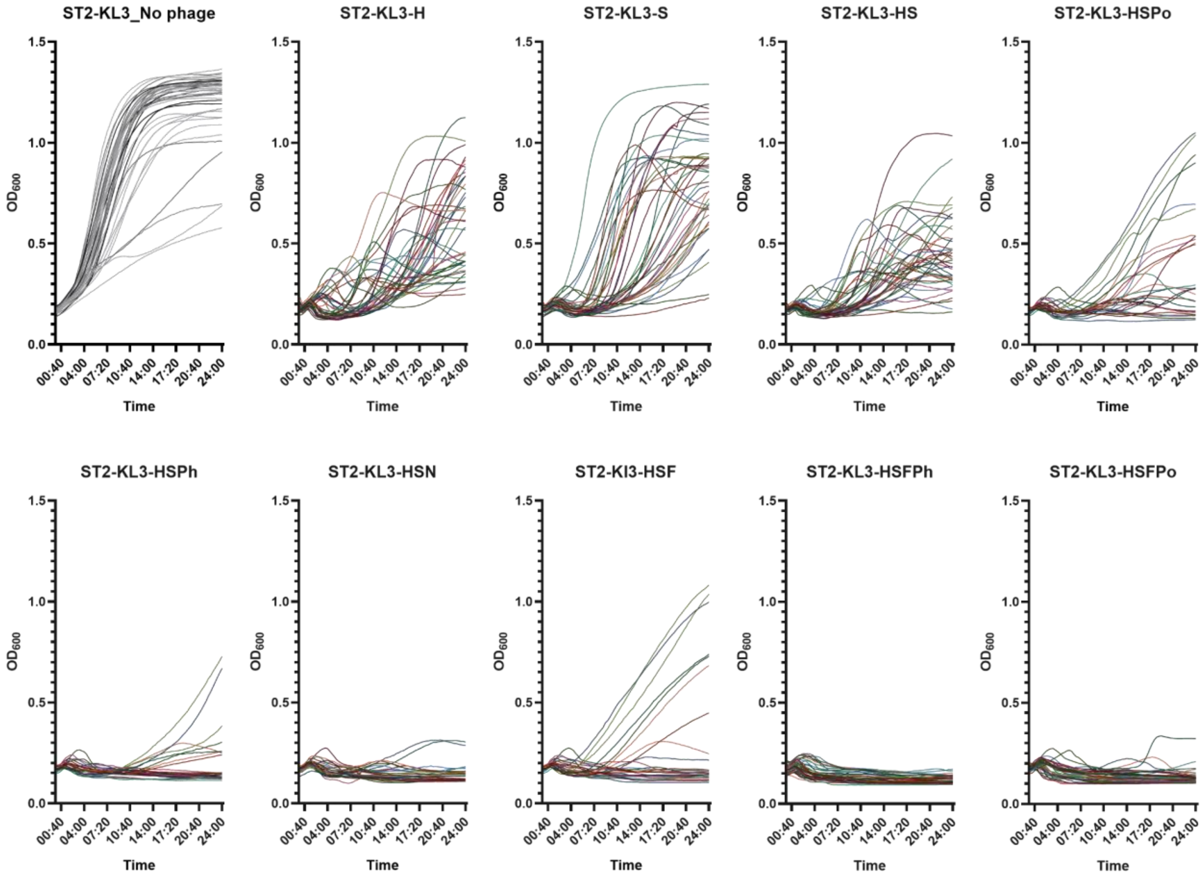


**Fig S16. The effect of different phages alone and in combination on the growth of ST2-KL3 CRAB isolates (n = 41, except the 3-phage combinations, where n = 27).** Growth curves were measured for 24 hours and represent individual strains**.** Area under the curve values are derived from these plots and are shown in Fig 6B. Each line represents the average growth ($OD_{600}$) in LB of 3 technical replicates. Each plot represents a different phage treatment. Abbreviation of phages: H – Highwayman, S – Silvergun, F – Fanak, Po – Porter, N - Navy-v2, Ph – PhT2-v2, $OD_{600}$ - optical density measured at 600 nm. Data is available in Supplementary Table 19.

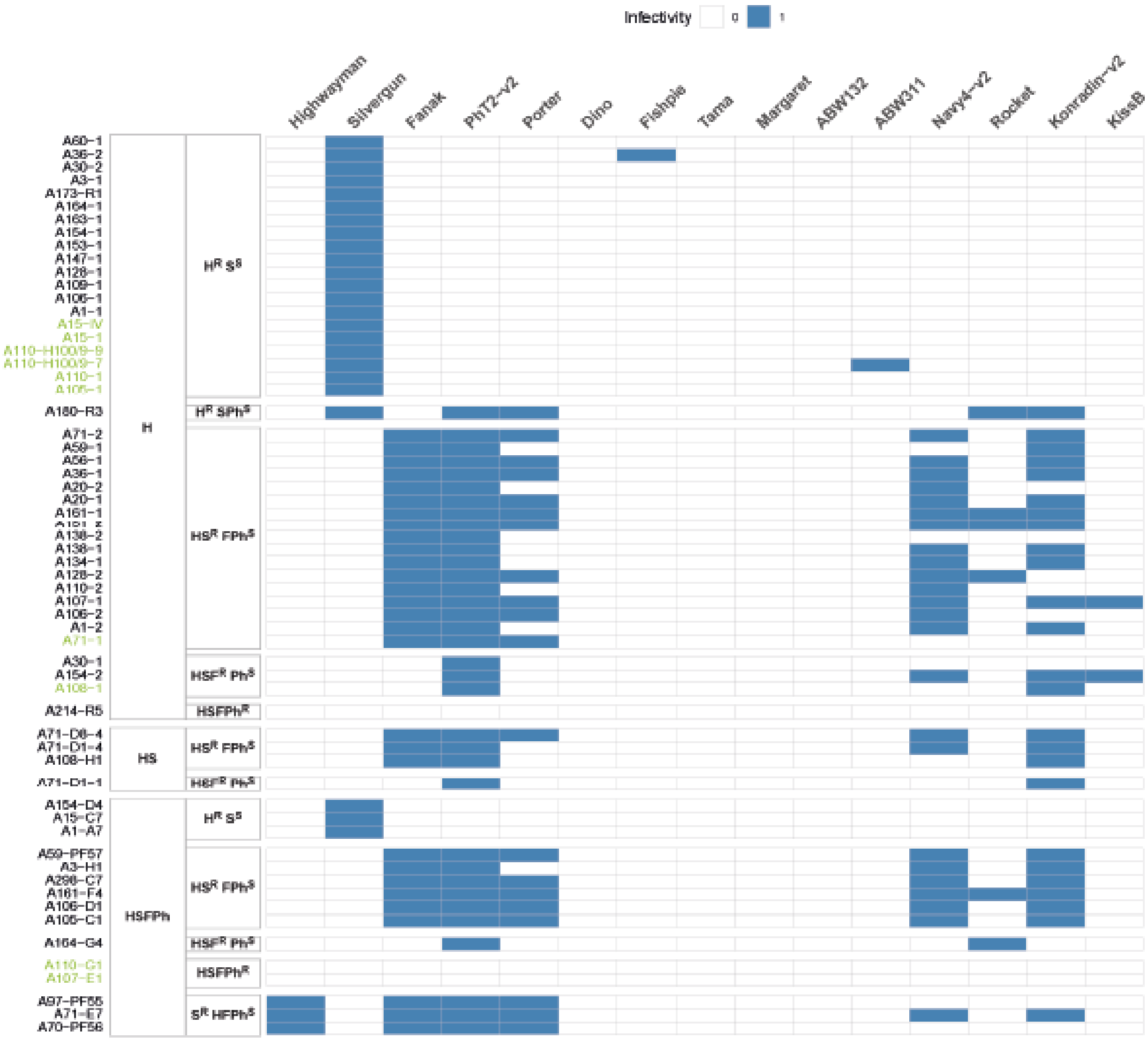

Infectivity
0
1
Highwayman
Silvergun
Fanak
PhT2-v2
Porter
Dino
Fishpie
Tama
Margaret
ABW132
ABW311
Navy4-v2
Rocket
Konradin-v2
KissB
H
HS
HSFPh

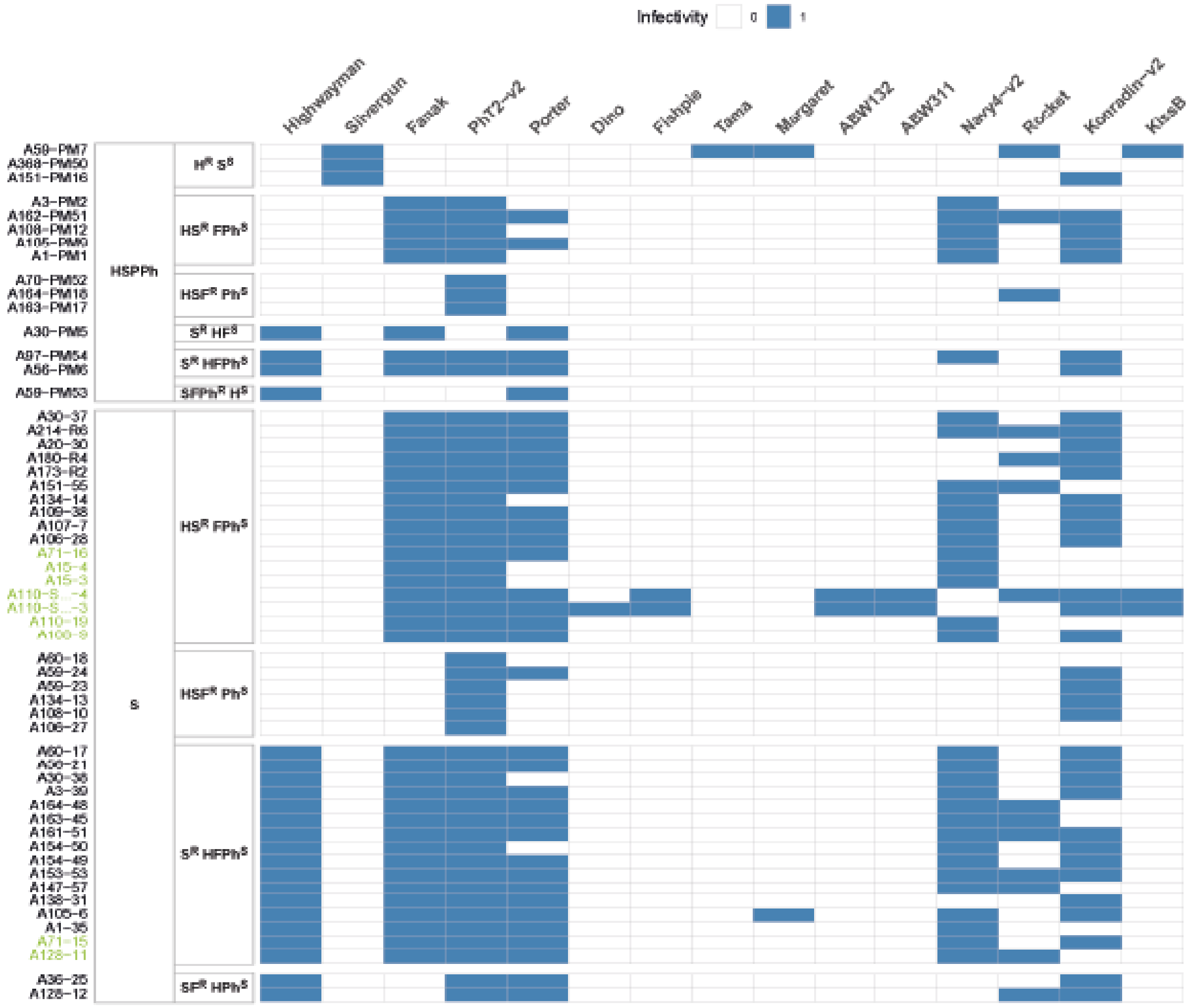


**Fig S17. Binary phage sensitivity profiles of the generated phage-resistant ST2-KL3 CRAB isolates (y axis, n = 149) against the 15 isolated phages (x axis).** First column in the heatmap shows the selection conditions, that is, individual phages or their combinations used for the co-incubation to induce phage resistance. The second column groups the isolated resistant ST2-KL3 CRAB cell lines according to their phage sensitivities to the phages in the HSFPh cocktail. The sensitivity profile was determined by spot assay. The colours blue and white indicate whether the phage can or cannot infect the tested isolate. Abbreviations of the phages: H – Highwayman, S – Silvergun, F – Fanak, Po – Porter, N - Navy-v2, Ph – PhT2-v2. The sequenced isolates are highlighted with green. Data is available in Supplementary Table 20.

Fig S18A

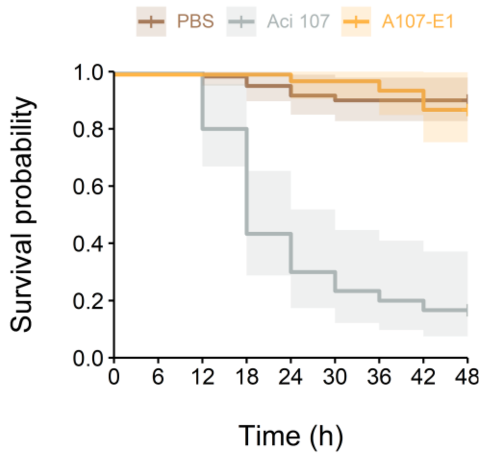


Fig S18B

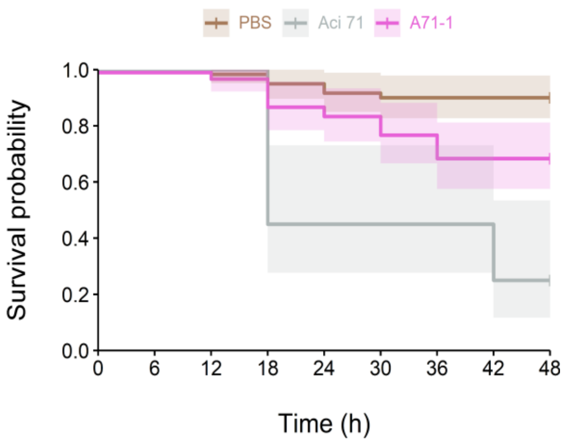


Fig S18C

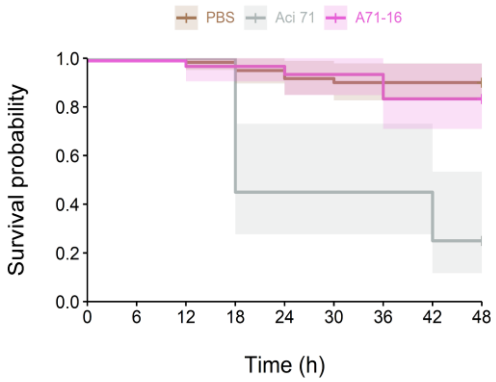


Fig S18D

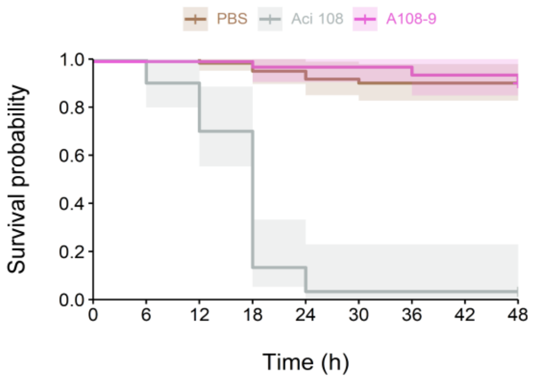

**Fig S18E**

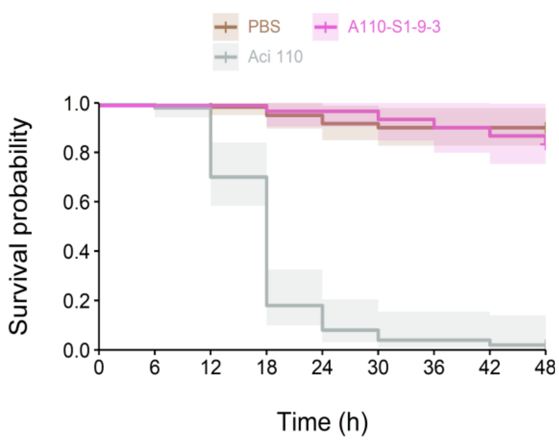


**Fig S18F**

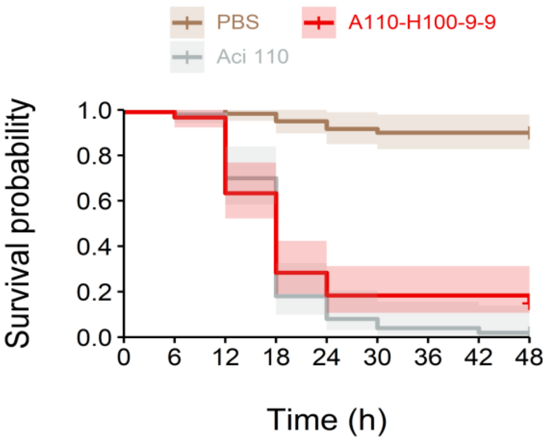


**Fig S18G**

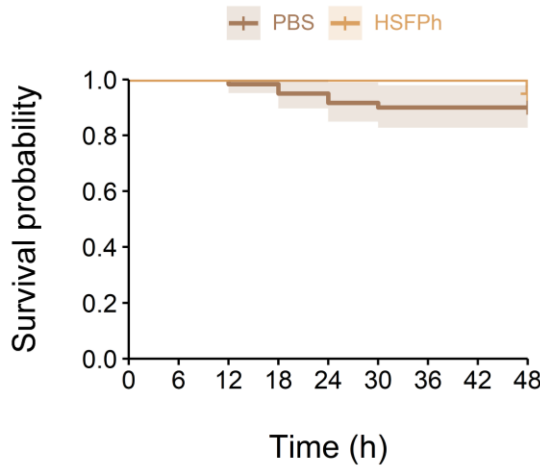


**Fig S18H**

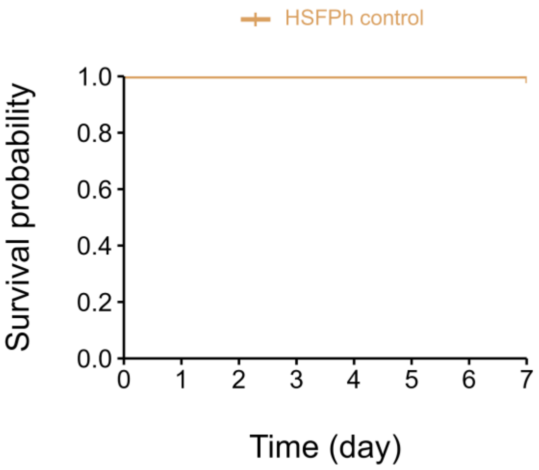


**Fig S18. A-F Kaplan-Meier curves showing the survival of *G. mellonella* larvae after infection with phage-resistant lines in comparison to their wild type counterparts.** HSFPh- (A, orange) and HS resistant lines (B-E, purple) have decreased virulence compared to their wild type counterparts (grey) ($p$ < 0.0001 from two-sided Log-rank test) (n = 10 larvae/group, ≥ 3 biological replicates/group, PBS – larvae injected only with PBS). H resistant line (F, red) is still virulent. **G-H** Kaplan-Meier curves showing the survival of *G. mellonella* larvae (G) (n = 10 larvae/group) and mice (H) (n = 5 mice/group) after the treatment with HSFPh cocktail without bacterial infection. The cocktail by itself does not have a negative effect on the survival of larvae or mice. Data is available in Supplementary Table 24.

log$_2$ change in MIC vs. wild type

-5 -4 -3 -2 -1 0

| Strain | MER | IMI | COL | LEV | T-S | Group |
|---|---|---|---|---|---|---|
| A151-PM16 | R | R | R to S | R | R | $H^R S^S$ |
| A15-IV | R | R | R to S | R | R | |
| A15-C7 | R | R | R to S | R | R | |
| A15-1 | R | R | R to S | R | R | |
| A110-H100/9-9 | R | R | R | R | R | |
| A110-1 | R to R | R | R to S | R | R | |
| A105-1 | R | R | S to S | R | R to R | |
| A1-A7 | R to I | R to R | S | R to R | R to R | |
| A71-E7 | R | R | R | R | R | $S^R HFPh^S$ |
| A71-15 | R to R | R to R | R | R | R | |
| A56-PM6 | R to R | R to R | R to R | R | R to R | |
| A128-11 | R | R | R to S | R | R to R | |
| A30-PM5 | R | R | S | R | R | $S^R HF^S$ |
| A71-16 | R to R | R to R | R to S | R | R | $HS^R FPh^S$ |
| A71-1 | R | R | R to S | R | R | |
| A3-H1 | R | R | S to S | R | R | |
| A298-C7 | R to R | R | S | R | R to R | |
| A161-F4 | R | R | S to S | R | R to R | |
| A15-3 | R | R | R to S | R | R | |
| A110-S1/9-3 | R | R | R to S | R | R | |
| A110-2 | R to R | R | R to S | R | R | |
| A110-19 | R | R | R to S | R | R | |
| A108-PM12 | R | R to R | S to S | R | R | |
| A108-9 | R to R | R | S | R | R | |
| A106-D1 | R | R | S to S | R | R | |
| A105-PM9 | R to R | R | R to S | R | R | |
| A105-C1 | R | R to R | R to S | R | R | |
| A1-PM1 | R | R | S to S | R | R | |
| A164-PM18 | R | R | S | R | R | $HSF^R Ph^S$ |
| A164-G4 | R | R | S to S | R | R | |
| A163-PM17 | R to R | R to R | R to S | R | R to R | |
| A108-1 | R to R | R | S to S | R | R to R | |
| A110-G1 | R to I | R to R | R | R to R | R to R | $HSFPh^R$ |
| A107-E1 | R to I | R to R | R to R | R to R | R to R | |

**Fig S19. Antibiotic sensitivity profile of phage-resistant strains (n = 34)**. Isolates are grouped based on their phage sensitivity profile, when only the phages from the HSFPh cocktail were taken into consideration for the grouping. The intensity of the colour green indicates the median ($n \geq 3$) of the $\log_2$ reduction in minimum inhibitory concentration (MIC) of the phage-resistant strains, compared to their wild type ancestor strain (the higher the reduction, the darker the intensity). Antibiotic susceptibility changes are also highlighted based on the MIC breakpoints from EUCAST data[7]. R - resistant, S - susceptible, I - intermediate (susceptible, increased exposure). In the case of colistin, 44% of the isolates transitioned from low-level resistant state to sensitive state (R to S), while 23% of the isolates became even more sensitive (S to S). The MIC of 5 antibiotics with clinical potential: MER-meropenem, IMI-Imipenem, COL-colistin, LEV-levofloxacin, TRM-Trimethoprim and Sulfamethoxazole was measured using the microbroth dilution method. Abbreviations of the phages: H – Highwayman, S – Silvergun, F – Fanak, N - Navy-v2, Ph – PhT2-v2. For details, see Methods and Supplementary Table 25.

**Supplementary tables**

**Supplementary Table 1.** *A. baumannii* strains
**Supplementary Table 2.** Top serotypes in each region between 2016-2022
**Supplementary Table 3.** Morisita dissimilarity indices between pairs of countries
**Supplementary Table 4.** Logistic regression results
**Supplementary Table 5.** Prevalence of MLS-CPS types in 2016-2022 compared with 2009-2015
**Supplementary Table 6.** Number of isolate pairs in genetic and geographic distance categories, global
**Supplementary Table 7.** Number of isolate pairs in genetic and geographic distance categories, regional
**Supplementary Table 8.** Number of isolate pairs in genetic and geographic distance categories, global with focus on European isolates
**Supplementary Table 9.** Strains from existing collections
**Supplementary Table 10.** *In vitro* phage sensitivity measurement results
**Supplementary Table 11.** Phylogenetic distances and phage profile distances between pairs of strains
**Supplementary Table 12.** List of previously characterised *A. baumannii* bacteriophages
**Supplementary Table 13.** Discovered phages
**Supplementary Table 14.** Metadata corresponding to the phages, created by the vConTACT2 tool that provides the underlying structure of the orthogroup networks in Fig 5.
**Supplementary Table 15.** The data structure of the vConTACT2 tool used in Fig 5.
**Supplementary Table 16.** Matrix of Average Nucleotide Identities (ANI) of phages in Fig 5E
**Supplementary Table 17.** A matrix of the identities of our depolymerase containing proteins compared to those in reference phages
**Supplementary Table 18.** Orthogroups mapping to the proteins of Silvergun in vConTACT2
**Supplementary Table 19.** Growth curves data
**Supplementary Table 20.** Phage-resistant *A. baumannii* isolates
**Supplementary Table 21.** Mutational profile of the ST2-KL3 phage-resistant isolates
**Supplementary Table 22.** Adsorption assay
**Supplementary Table 23.** Distance matrixes measured by the IR biotyper
**Supplementary Table 24.** Survival experiments
**Supplementary Table 25.** Antibiotic susceptibility profiles of the phage-resistant isolates
**Supplementary Table 26.** Antibiograms for *A. baumannii* isolates
**Supplementary Table 27.** Antibiotic resistance determinants and their statistical relationship with measured carbapenem resistance

**Supplementary Table 28.** Efficiency of plating (EOP) of Silvergun phage on ST2-KL2 and ST2-KL3 clinical isolates

**Supplementary Note S1.** Genomic and morphological analysis of the 15 isolated *A. baumannii* bacteriophages

**NOTES:**

Gene of interest: The presence or absence of anti-crisper, endolysin and lytic cycle related genes predicted using customary python script (see methods).

### identifier of hit [cognate organism of blastp hit] Hit [identifier of the Open Reading Frame (ORF)in pos; [position of the ORF on the related genome]

Predicted therapeutic suitability : The presence or absence of temperate markers, antimicrobial resistance (AMR) genes and virulence genes predicted using phageLeads online tool[8] (https://phageleads.dk/)

qcov: query coverage

Per.ident: percentage identity

n.d: not detected.

PFU: Plaque Forming Units

Host bacteria: clinical isolate used to enrich and isolate the phage.

Enzymatic activity: presence or absence of halo zone formed by the phages on bacterial lawn, which is indicative of presence of phage-derived enzyme leading to the destruction of bacterial cell wall causing cell death.

Figures: **A,** Transmission electron microscopy pictures of the phages stained with uranyl-acetate. **B,** Plaque morphology of the phages formed on the host bacterial lawn after overnight incubation. **C,** Genome maps were generated using SnapGene. Different coloured arrows represent predicted CDSs coding different functions. red: lysis. grey: hypothetical protein (hp), blue: DNA replication, transcription, packaging, and metabolism function. green: phage structure. The direction of arrows represents the direction of transcription.

#### vB_AbaM_Highwayman

Highwayman is a novel phage of the Myoviridae family, Obolenskvirus genus. Based on sequence similarity it shares with its closest predicted relative phage (Acinetobacter phage WCHABP1/ NCBI Accession: NC_041966.1; 81% coverage; 92,24% percentage identity- lytic phage). The novelty of its taxonomic placement is further established by the position of its terminase large subunit and capsid protein on phylogenetic tree that has been constructed using the homologous proteins of its closest relatives (Fig 4-5) Highwayman has a dsDNA genome with a 37,95 GC% and 116 predicted ORFs of which 32 (~ 27,59 %) could be assigned to putative functions with high confidentiality based on domain prediction and blastp analysis. Silvergun has been predicted to possess temperate lifestyle by PhageAI, albeit with a low confidence value (61,54 %) with the software denoting it doesn't have enough information in its database by which it could provide a sufficiently reliable prediction about the lifestyle of Highwayman. Despite this however, Highwayman showed strictly lytic behaviour, with further experimental results we generated supporting its non-lysogenic activity on the *A. baumannii* strains applied in present study (See supplementary figure 15). Furthermore, Highwayman lacks virulence or antibiotic resistance-related genes that would undermine its eligibility for therapeutic applications. Based on gene synteny and blastp similarity match we were able to identify the tail fiber region of Highwayman which, surprisingly, didn't carry any domain regions that could be predicted using InterProscan. It did however contain a hydrolase domain identified by the Phyre2 web service, corresponding to PDB domain 6TGF Aside from the endolysin protein, we were able to identify depolymerase domains on the tail tape measure protein (IPR023346). We didn't manage to identify further tail fiber/spike regions on Highwayman. However, Highwayman was able to effectively target and propagate on 98% of the studied ST2 KL3 isolates. Given that the tail fiber of Silvergun and Highwayman shares very low similarity with each other it is possible that Highwayman can achieve infection on the ST2 KL3 strains that is different from that of Silvergun, and which domains also exhibit a level of novelty that makes them difficult to detect. As such, the exact receptor of Highwayman remains unknown. Lastly, TEM confirmed the bioinformatic prediction of Highwayman’s morphology, that it clearly belongs to the myovirus category.

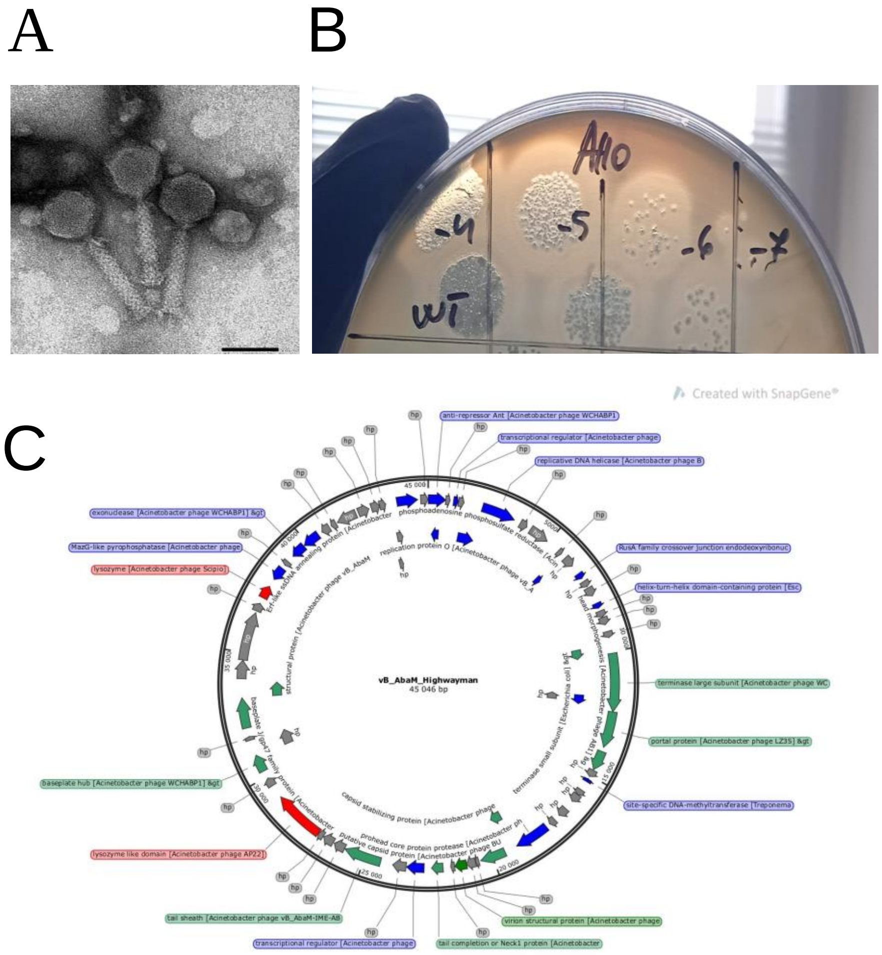


**Closest blastn hit**: Acinetobacter phage WCHABP1.

qcov: 81%, per.ident: 92,24 %

**Predicted lifestyle** (PhageAI): Temperate 61.54%. (NOTE prediction is not correct due to small sample for teaching up the algorithm, is written on the phageAI page)

**Predicted taxonomy**: Myoviridae; Obolenskvirus.

**Predicted therapeutic suitability:** no harmful genes detected

**Genes of interest:**

- Anti-CRISPR genes: n.d

- Endolysin genes:

- lysozyme [Acinetobacter phage Scipio] Hit_29 in pos 37258-37770
- lysozyme like domain [Acinetobacter phage AP22] > emb|CCH57752.1|
- lysozyme like domain [Acinetobacter phage AP22]    Hit_90 in pos 27015-29045

- Lytic cycle related genes:

- HNH endonuclease [Acinetobacter phage vB_AbaM_BP10]    Hit_75 in pos 11799-12218.
- site-specific DNA-methyltransferase [Treponema sp.]    Hit_132 in pos; reverse 15270-15148.

**-Host bacteria**: Aci 3 (ST2-KL3)

**-Genome size**:  45046bp

**-Enzymatic activity:** yes

#### vB_AbaS_Silvergun

**Closest blastn hit:** Siphoviridae environmental samples clone NHS-Seq1

qcov: 21%; per.ident: 75,70%

**Predicted lifestyle (phageAI)**: virulent 70,71%

**Predicted taxonomy**:  Siphoviridae; Jerseyvirus

**Predicted therapeutic suitability:** no harmful genes detected

**Genes of interest:**

- Anti-CRISPR genes:  n.d

- Endolysin genes:

- endolysin [Stenotrophomonas phage CUB19]    Hit_57 in pos 15836-16477
- CD225/dispanin family protein [Lysobacter oculi] Hit_194 in pos; reverse 8370-8008
- D-alanyl-D-alanine carboxypeptidase family protein [Phaeobacter sp. LSS9] Hit_197 in pos; reverse 2856-2710

- Lytic cycle related genes:

- HNH endonuclease [Aquamicrobium zhengzhouense] Hit_108 in pos 12897-13268
- HNH endonuclease [Stenotrophomonas sp. AG209] > gb|RIA19692.1| AP2 domain-containing protein [Stenotrophomonas sp. AG209]    Hit_137 in pos; reverse 44749-44204

- **Host bacteria:**  Aci 108 (ST2-KL3)

**-Genome size:** 46864 bp

**-Enzymatic activity:** yes

One of the several isolated phages in this study, nicknamed Silvergun, exhibited a unique and novel genetic makeup in comparison to the existing phage genomes in our database and the NCBI RefSeq database. It shares low similarity with its closest predicted relative, a phage belonging to the Siphoviridae family, Jerseyvirus genus (21% coverage; 75,70% percentage identity) and so most probably represents a member of a novel family, since it shares low similarity with both members of *Drexler*- and *Siphoviridae* phages. Investigating its orthogroups using a custom script employing blastp and MMseqs, we have discovered that it has a similar minor tail protein, peptidase, and tail assembly protein as the Drexlerviridae phages, and on top of that also shares 6 hypothetical proteins a DUF2213 a Fibronectin III type and a DUF2184 containing domain with the suspected Siphoviridae phage, along with the following proteins: Prophospohatase, holin, capsid assembly protein, tail protein, endolysin and gene product 10. Altogether it shares 30% of its orthogroups with the Siphoviridae isolate and less than 10% with the Drexlerviridae phages, when the cut-off for orthogroup clustering was 50% coverage and 50% identity between the two proteins. Its double-stranded DNA is 46,864 bp in length with 52,66 % GC content and predicted to contain 137putative ORFs. Silvergun is predicted to have a lytic life cycle based on PhageAI prediction and lacks any kind of obvious lysogeny related genes such as integrases or repressors. It also seems to be devoid of any virulence or antibiotic resistance conferring genes which would exclude it from therapeutic applications. Importantly, Silvergun can infect two distinct *A.baumannii* strains of our collection: ST2 KL2 and ST2 KL3 (figure 2A,2B). and shows an EOP $\geq 0.1$

on two of the most widely and rapidly spreading variants of pathogenic *A. baumannii* strains which is therapeutically acceptable as described in other studies.[9,10,11] The phage was inefficient toward only one ST2-KL2 strain (Aci 320) (Fig 2B). Though the tail fiber region of this phage couldn't be identified based solely on RefSeq similarity match, we were able to reveal the genetic region encoding this protein by parsing through the domain repertoires of the individual ORFs. Silvergun's tail fiber contains an immunoglobulin (Ig)-like domain -a fibronectin type III (Fn3) domain in particular (IPR003961)- along with a domain of unknown function (IPR015406), a carbohydrate-binding domain (IPR036573) and a Tip attachment protein J (IPR032876) known to facilitate attachment to host receptors. Its tail spike protein carries a depolymerase domain, a pectin lyase (IPR012334) that likely plays an important role in degrading the capsule of the host bacteria of Silvergun. Since there were no other detectable depolymerase domains on the identified phage tail proteins, it stands to reason the Silvergun can infect strains of differing capsule types due to its potentially promiscuous IPR012334 domain. Other than these observations, the tail fiber region of Silvergun didn't show similarity with phages whose host specificity or target receptor is known. Unfortunately, we didn't manage to pinpoint the capsid proteins of Silvergun neither by relying on similarity matches, nor by examining the domains of its proteome, further implying its taxonomically novel affiliation. As stated before, Silvergun has been predicted to have a siphovirus morphology. However, transmission electron microscopic analysis contradicts this result, as Silvergun displays a characteristic myovirus morphology (Fig 2).

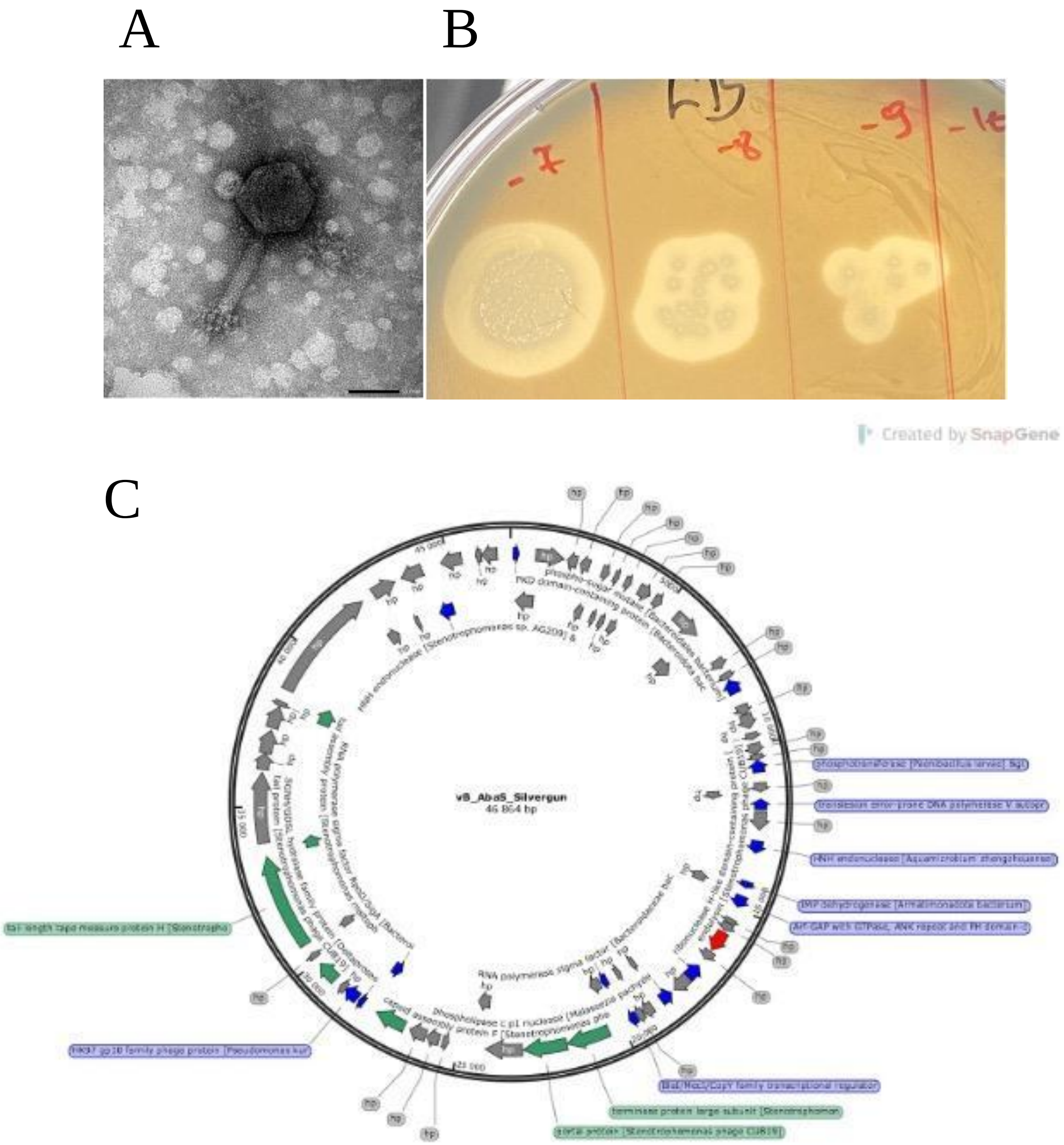
A
B
-7
-8
-9
Created by SnapGene
C
vB_AbaS_Silvergun
46 864 bp

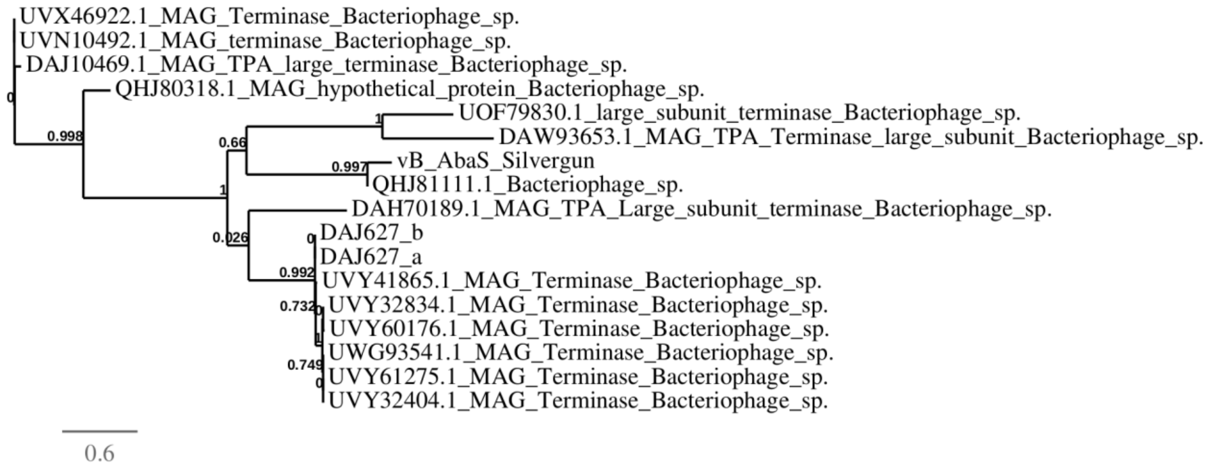


**Figure 1**. Position of Silvergun on a phylogenetic tree based on the amino acid sequence of the terminase large subunit.

A

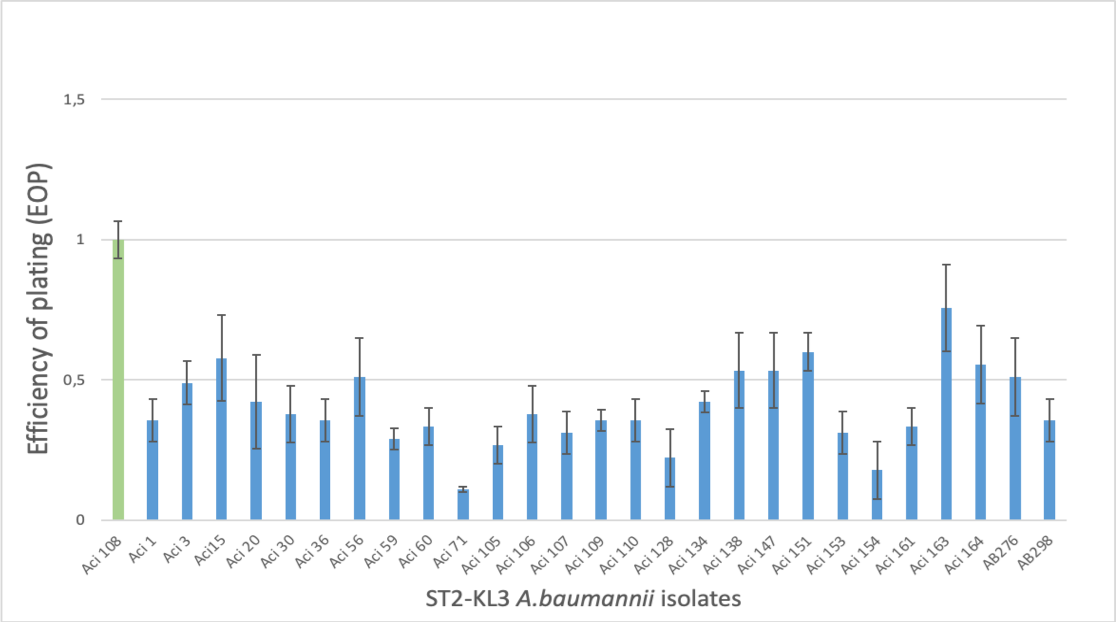


B

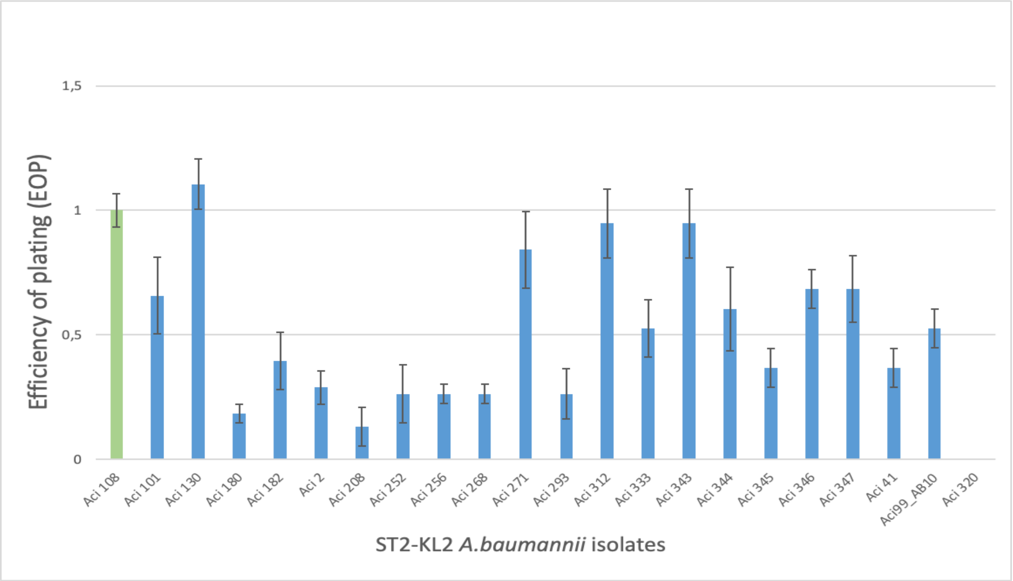


**Figure 2**. Efficiency of plating (EOP) of Silvergun phage on 28 ST2-KL3 (A) and 20 ST2-KL2 (B) isolates (for details, see Methods). Aci 108 indicated with green is the reference isolate on which Silvergun phage was isolated. Error bars represent standard deviation (n = 3). Data are shown in Supplementary Table S28.

#### vB_AbaP_Fishpie

**Closest blastn hit**: Acinetobacter phage vB_AbaP_ APK44

qcov: 90%, per.ident: 93,99%

**Predicted lifestyle** (phageAI): Virulent 100%

**Predicted taxonomy**: Autographiviridae; Friunavirus

**Predicted therapeutic suitability:** no harmful genes detected.

**Genes of interest:**

- Anti-CRISPR genes: n.d

- Endolysin genes:

  - endolysin [Acinetobacter phage IME-200] Hit_9 in pos 27079-27636
  - holin/anti-holin [Acinetobacter phage IME-200] Hit_56 in pos 26757-27092

**-** Lytic cycle related genes:

  - endonuclease VII [Acinetobacter phage phiAB1] Hit_2 in pos 5983-6423
  - HNH homing endonuclease [Acinetobacter phage vB_AbaP_APK87] Hit_37 in pos 90-536
  - tailspike protein [Acinetobacter phage vB_AbaP_APK2] Hit_55 in pos 24663-26744

**Host bacteria:** Aci 7 (ST2-KL2)

**Genome size:** 40736 bp

**Enzymatic activity**: Yes

A B

C

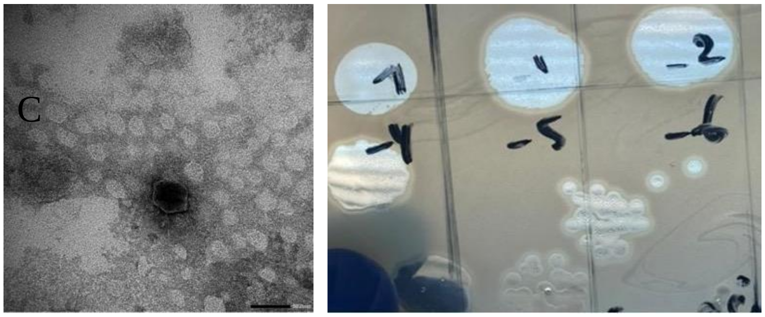

C

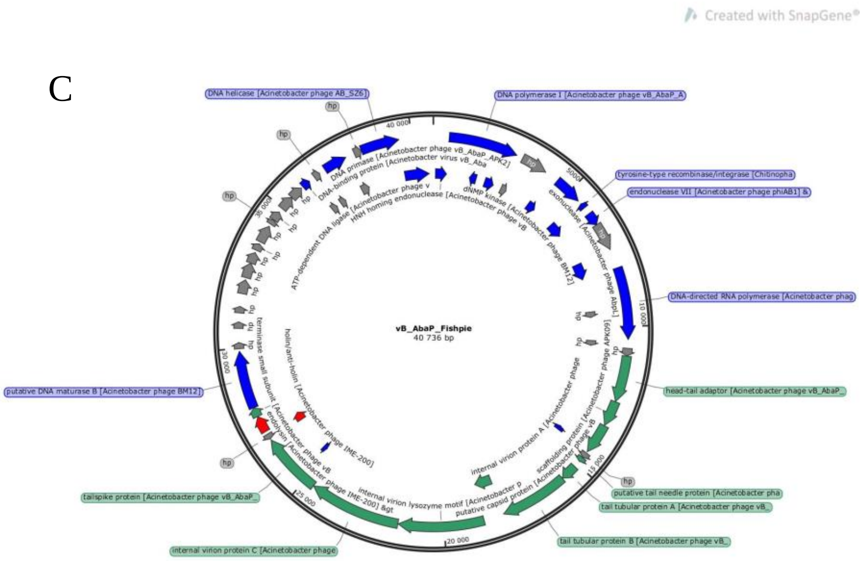
Created with SnapGene®
vB_AbaP_Fishpie
40 736 bp
DNA helicase [Acinetobacter phage AB_SZ6]
DNA polymerase I [Acinetobacter phage vB_AbaP_A
tyrosine-type recombinase/integrase [Chitinopha
endonuclease VII [Acinetobacter phage phiAB1] &
DNA-directed RNA polymerase [Acinetobacter phag
head-tail adaptor [Acinetobacter phage vB_AbaP_
putative tail needle protein [Acinetobacter pha
tail tubular protein A [Acinetobacter phage vB_
tail tubular protein B [Acinetobacter phage vB_
internal virion protein C [Acinetobacter phage
tailspike protein [Acinetobacter phage vB_AbaP_
putative DNA maturase B [Acinetobacter phage BM12]
DNA primase [Acinetobacter phage vB_AbaP_APK2]
internal virion lysozyme motif [Acinetobacter phage IME-200] &gt

#### vB_AbaP_ABW311

**Closest blastn hit**: Acinetobacter virus vB_AbaP_AGC01

qcov: 90%, per.ident: 95,97%

**Predicted lifestyle** (phageAI): Lytic 99,68%

**Predicted taxonomy**: Autographiviridae; Friunavirus

**Predicted therapeutic suitability:** no harmful genes detected.

**Genes of interest:**

- Anti-CRISPR genes: n.d

- Endolysin genes:

  - endolysin [Acinetobacter phage vB_AbaP_APK32] Hit_21 in pos 37201-37758
  - holin/anti-holin [Acinetobacter phage SH-Ab 15519] Hit_66 in pos 36879-37214

- Lytic cycle related genes:

  - putative HNH homing endonuclease [Acinetobacter phage vB_AbaP_ABWU2101] Hit_9 in pos 11647-12174
  - exonuclease [Acinetobacter phage vB_ApiP_P2] > gb|AZU99224.1| Hit_58 in pos 14517-15470
  - DNA endonuclease VII [Acinetobacter phage vB_AbaP_APK44] Hit_59 in pos 16023-16466
  - tailspike protein [Acinetobacter phage vB_AbaP_APK32] Hit_65 in pos 34833-36869

**Host bacteria:** Aci 311 (ST2-KL32)

**Genome size:** 40892 bp

**Enzymatic activity**: yes

A

B

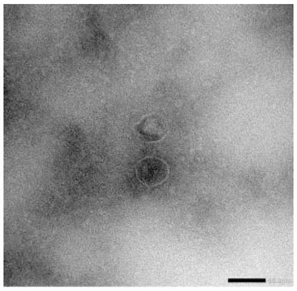

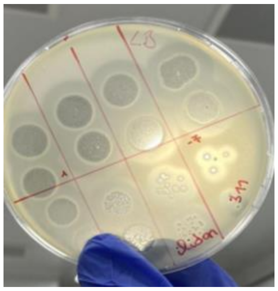

C

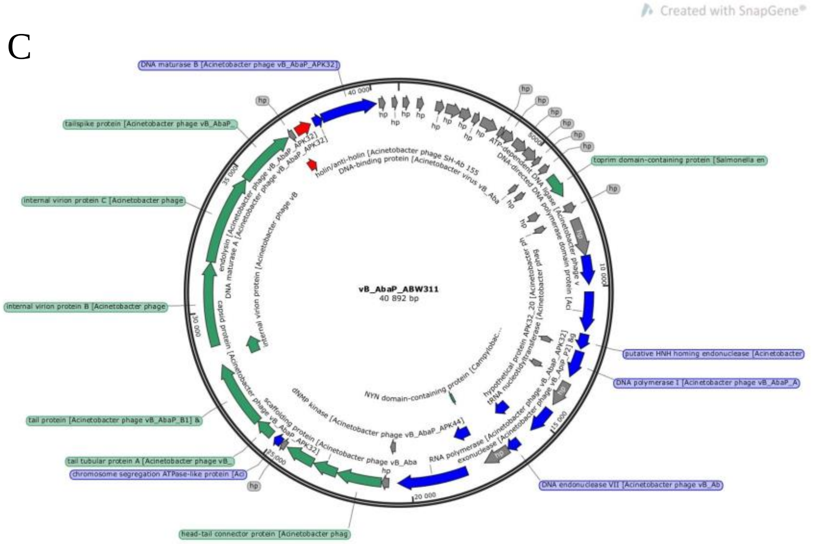
Created with SnapGene®
vB_AbaP_ABW311
40 892 bp
DNA maturase B [Acinetobacter phage vB_AbaP_APK32]
tailspike protein [Acinetobacter phage vB_AbaP_
internal virion protein C [Acinetobacter phage
internal virion protein B [Acinetobacter phage
tail protein [Acinetobacter phage vB_AbaP_B1] &
tail tubular protein A [Acinetobacter phage vB_
chromosome segregation ATPase-like protein [Ac
head-tail connector protein [Acinetobacter phag
toprim domain-containing protein [Salmonella en
putative HNH homing endonuclease [Acinetobacter
DNA polymerase I [Acinetobacter phage vB_AbaP_A
DNA endonuclease VII [Acinetobacter phage vB_Ab
40 000
35 000
30 000
25 000
20 000
15 000
10 000
5000

#### vB_AbaP_Tama

**Closest blastn hit**: Acinetobacter virus vB_AbaP_AGC01

qcov: 89%, per.ident: 95,58%

**Predicted lifestyle** (phageAI): Virulent 99,61%

**Predicted taxonomy**: Autographiviridae; Friunavirus

**Predicted therapeutic suitability:** no harmful genes detected

**Genes of interest:**

- Anti-CRISPR genes: n.d

- Endolysin genes:

  - holin/anti-holin [Acinetobacter phage SH-Ab 15519] Hit_25 in pos 39661-39996

- Lytic cycle related genes:

  - endonuclease VII [Acinetobacter phage phiAB1] Hit_16 in pos 18658-19098
  - tail fiber protein [Acinetobacter phage vB_AbaP_B1] Hit_24 in pos 37369-39651
  - putative HNH homing endonuclease [Acinetobacter phage APK20] Hit_59 in pos 14307-14834

**Host bacteria:** Aci 144 (ST2-KL9)

**Genome size:** 41641 bp

**Enzymatic activity**: Yes

A B

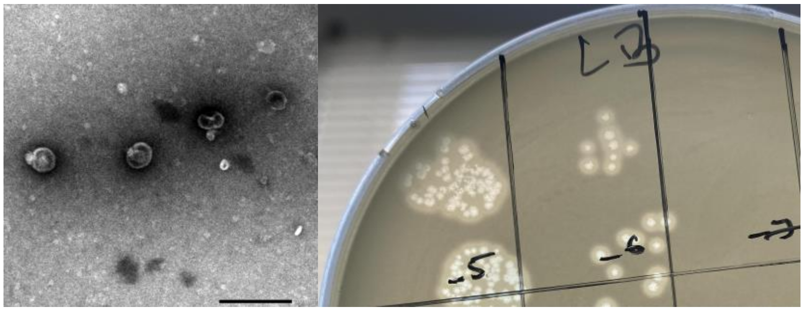


C

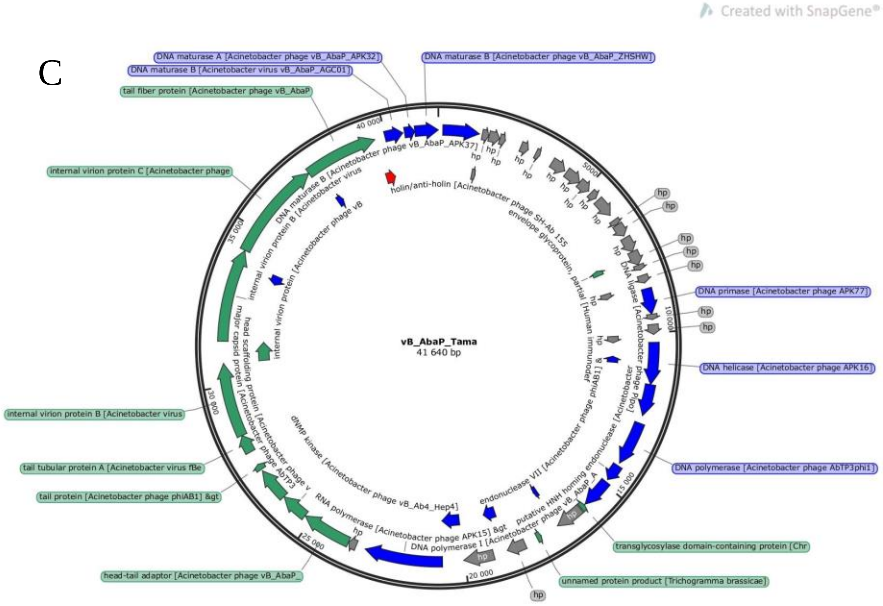

#### vB_AbaP_Fanak

**Closest blastn hit**: Acinetobacter phage APK09.

qcov: 94%, per.ident: 93,54%

**Predicted lifestyle** (phageAI): virulent 99,60%

**Predicted taxonomy**: Autographiviridae; Friunavirus

**Predicted therapeutic suitability:** no harmful genes detected

**Genes of interest:**

- Anti-CRISPR genes: n.d

- Endolysin genes:

  - holin/anti-holin [Acinetobacter phage SH-Ab 15519] Hit_6 in pos 16252-16587
  - internal virion protein with endolysin domain [Acinetobacter phage Fri1] Hit_30 in pos 10859-13957

- Lytic cycle related genes:

  - tail fiber protein [Acinetobacter phage vB_AbaP_B1] Hit_5 in pos 13960-16242
  - HNH endonuclease [Acinetobacter phage vB_AbaP_APK128] Hit_43 in pos 33449-33871
  - endonuclease VII [Acinetobacter phage vB_AbaP_B1] Hit_48 in pos 36770-37210

**Host bacteria:** A110-19 (ST2-KL3)

**Genome size:** 41388 bp

**Enzymatic activity**: Yes

A

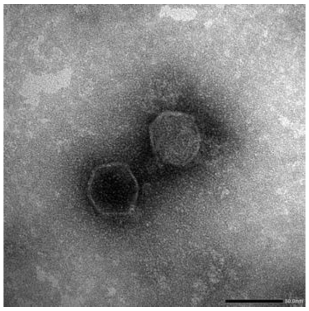

B

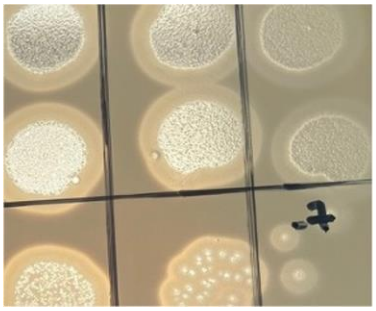

C

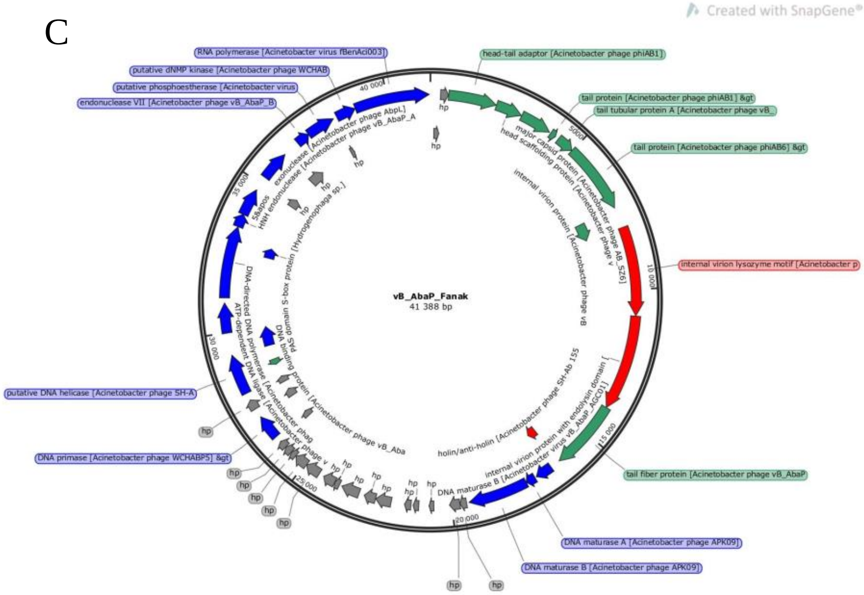
Created with SnapGene®
vB_AbaP_Fanak
41 388 bp
RNA polymerase [Acinetobacter virus fBenAci003]
putative dNMP kinase [Acinetobacter phage WCHAB
putative phosphoesterase [Acinetobacter virus
endonuclease VII [Acinetobacter phage vB_AbaP_B
head-tail adaptor [Acinetobacter phage phiAB1]
tail protein [Acinetobacter phage phiAB1] &gt
tail tubular protein A [Acinetobacter phage vB_
tail protein [Acinetobacter phage phiAB6] &gt
internal virion lysozyme motif [Acinetobacter p
tail fiber protein [Acinetobacter phage vB_AbaP
DNA maturase A [Acinetobacter phage APK09]
DNA maturase B [Acinetobacter phage APK09]
putative DNA helicase [Acinetobacter phage SH-A
DNA primase [Acinetobacter phage WCHABP5] &gt
hp
5000
10 000
15 000
20 000
25 000
30 000
35 000
40 000

#### vB_AbaP_Margaret

**Closest blastn hit**: Acinetobacter virus vB_AbaP_AGC01

qcov: 90%, per.ident: 97,62%

**Predicted lifestyle** (phageAI): Virulent 99,64%

**Predicted taxonomy**: Autographiviridae; Friunavirus

**Predicted therapeutic suitability:** no harmful genes detected

**Genes of interest:**

- Anti-CRISPR genes: n.d

- Endolysin genes:

- putative endolysin [Acinetobacter phage vB_AbaP_PD-AB9] Hit_39 in pos; reverse 27207-26650
- holin/anti-holin [Acinetobacter phage Abp1] Hit_86 in pos; reverse 27529-27194

- Lytic cycle related genes:

- HNH endonuclease [Acinetobacter phage vB_ApiP_P2] Hit_55 in pos; reverse 12558-12112
- exonuclease [Acinetobacter phage vB_ApiP_P2] Hit_60 in pos; reverse 8478-7525
- endonuclease VII [Acinetobacter phage phiAB1] Hit_61 in pos; reverse 6972-6532
- tail fiber protein [Acinetobacter phage SWH-Ab-3] Hit_70 in pos; reverse 29759-27660

**Host bacteria:** Aci 7 (ST2-KL2)

**Genome size:** 41685 bp

**Enzymatic activity**: yes

A B

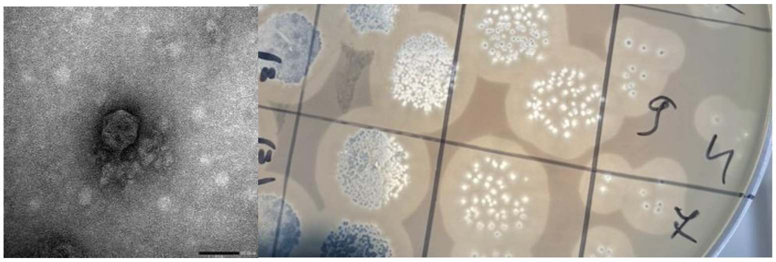

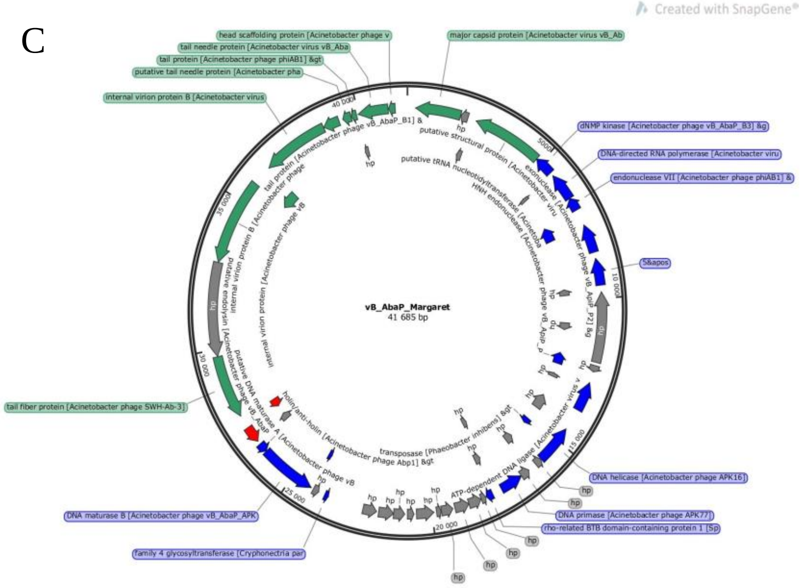
Created with SnapGene®
head scaffolding protein [Acinetobacter phage v
tail needle protein [Acinetobacter virus vB_Aba
tail protein [Acinetobacter phage phiAB1] &gt
putative tail needle protein [Acinetobacter pha
internal virion protein B [Acinetobacter virus
major capsid protein [Acinetobacter virus vB_Ab
dNMP kinase [Acinetobacter phage vB_AbaP_B3] &g
DNA-directed RNA polymerase [Acinetobacter viru
endonuclease VII [Acinetobacter phage phiAB1] &
5&apos
vB_AbaP_Margaret
41 685 bp
tail fiber protein [Acinetobacter phage SWH-Ab-3]
DNA helicase [Acinetobacter phage APK16]
DNA primase [Acinetobacter phage APK77]
rho-related BTB domain-containing protein 1 [Sp
DNA maturase B [Acinetobacter phage vB_AbaP_APK
family 4 glycosyltransferase [Cryphonectria par
transposase [Phaeobacter inhibens] &gt
holin/anti-holin [Acinetobacter phage Abp1] &gt
putative structural protein [Acinetobacter viru
putative tRNA nucleotidyltransferase [Acinetoba
HNH endonuclease [Acinetobacter pha
ATP-dependent DNA ligase [Acinetobacter virus v
putative DNA maturase A [Acinetobacter phage vB
putative endolysin [Acinetobacter phage vB_AbaP
internal virion protein [Acinetobacter phage vB
40 000
35 000
30 000
25 000
20 000
15 000
10 000
5000
hp

#### vB_AbaM_Porter

**Closest blastn hit**: Acinetobacter phage Morttis

qcov: 93%, per.ident: 96,87%

**Predicted lifestyle** (phageAI): Lytic 64.41%

**Predicted taxonomy**:  Myoviridae; Tegunavirus

**Predicted therapeutic suitability:** no harmful genes detected

(The phage was predicted to contain an integrase which potentially can enable access to the temperate lifestyle cycle, but there was no evidence on its functionality as the Blastp hits results showed similarity to Hypothetical proteins).
(vB_AbaM_Porter_CDS_1_164_119855_120862)

**Genes of interest:**

- Anti-CRISPR genes:

- type I-C CRISPR-associated protein Cas8c/Csd1 [Deltaproteobacteria bacterium] Hit_473 in pos; reverse 7444-7256

- Endolysin genes:

- lysis inhibition: accessory protein [Acinetobacter phage AbTZA1]   Hit_31 in pos 51151-51393
- Chain A, Endolysin [Acinetobacter phage AbTZA1]    Hit_110 in pos 8414-8977
- lysis inhibition [Acinetobacter phage AbTZA1]    Hit_172 in pos 112526-112864
- nardilysin-like isoform X1 [Varroa destructor] Hit_254 in pos 64314-64436
- lytic transglycosylase domain-containing protein [Caulobacter sp. UNC358MFTsu5.1]    Hit_332 in pos; reverse 154400-154224
- M3 family oligoendopeptidase [Phycisphaerales bacterium] Hit_338 in pos; reverse 128288-128145
- S8 family serine peptidase [Noviherbaspirillum humi] Hit_339 in pos; reverse 125486-125331
- baseplate hub and tail lysozyme [Acinetobacter phage AB-Navy71]    Hit_350 in pos; reverse 100649-98865

- holin [Acinetobacter phage AbTZA1] Hit_383 in po Mainzs; reverse 21680-20958
- hemolysin III family protein [Holdemania filiformis] Hit_466 in pos; reverse 28729-28454

- Lytic cycle related genes:

- recombination endonuclease subunit [Acinetobacter phage vB_AbaM_PhT2] Hit_71 in pos 124777-125790
- DNA endonuclease IV [Acinetobacter phage Maestro] Hit_119 in pos 17105-17707
- RNA ligase and tail fiber protein attachment catalyst [Acinetobacter phage AbTZA1] Hit_137 in pos 45302-46450
- tail fiber protein [Acinetobacter phage AbTZA1] Hit_166 in pos 106475-107041
- homing endonuclease [Escherichia phage IME08] Hit_261 in pos 71337-72005
- homing endonuclease [Acinetobacter phage AbTZA1] Hit_280 in pos 110427-110858
- endonuclease VII [Acinetobacter phage AbTZA1] Hit_289 in pos 122160-122642
- SbcC-like subunit of palindrome specific endonuclease [Acinetobacter phage AbTZA1] Hit_294 in pos 126387-128069
- baseplate wedge tail fiber connector [Acinetobacter phage Maestro] Hit_353 in pos; reverse 92018-91152
- short tail fiber [Acinetobacter phage Maestro] Hit_355 in pos; reverse 88676-87150
- long tail fiber proximal subunit [Acinetobacter phage Maestro] Hit_381 in pos; reverse 32216-28338
- homing endonuclease [Acinetobacter phage AB-Navy4] Hit_398 in pos; reverse 160390-159452
- homing endonuclease [Acinetobacter phage AbTZA1] Hit_459 in pos; reverse 40822-40568
- long tail fiber distal hinge connector [Acinetobacter phage Maestro] Hit_467 in pos; reverse 27139-26435

- tail fiber protein [Acinetobacter phage AbTZA1]    Hit_468 in pos; reverse 26422-22289
- homing endonuclease [Acinetobacter phage vB_AbaM_PhT2]    Hit_535 in pos; reverse 40272-39487

**Host bacteria:**  Aci 136 (ST2-KL12)

**Genome size:** 167177 bp

**Enzymatic activity**: No

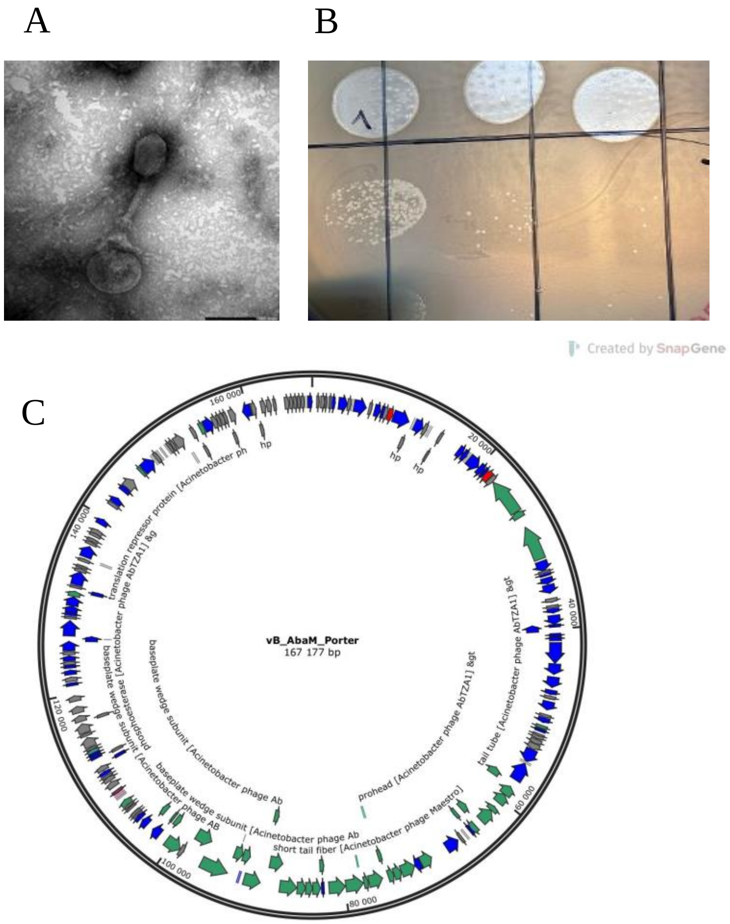

#### vB_AbaP_Dino

**Closest blastn hit**: Acinetobacter phage BM12.

qcov: 96%, per.ident: 98,63%

**Predicted lifestyle** (phageAI): Virulent 100%.

**Predicted taxonomy**: Autographiviridae; Friunavirus.

**Predicted therapeutic suitability:** no harmful genes detected

**Genes of interest**:

- Anti-CRISPR genes: n.d.

- Endolysin genes:

  - endolysin [Acinetobacter phage IME-200] Hit_56 in pos; reverse 1661-1104
  - holin/anti-holin [Acinetobacter phage IME-200] Hit_105 in pos; reverse 1983-1648

- Lytic cycle related genes:

  - tail fiber protein [Acinetobacter phage BM12] Hit_52 in pos; reverse 13055-10764
  - HNH endonuclease [Acinetobacter phage phiAB1] Hit_92 in pos; reverse 29091-28645
  - HNH family endonuclease [Acinetobacter phage Fri1] Hit_93 in pos; reverse 27855-27367
  - endonuclease VII [Acinetobacter phage vB_AbaP_B1] Hit_95 in pos; reverse 22797-22357
  - tailspike protein [Acinetobacter phage vB_AbaP_APK2] Hit_104 in pos; reverse 4077-1996

**Host bacteria:** Aci 101 (ST2-KL2)

**Genome size:** 41561 bp

**Enzymatic activity**: yes

A

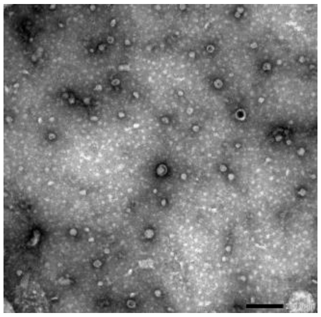

B

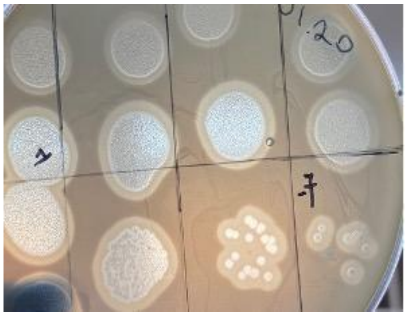

C

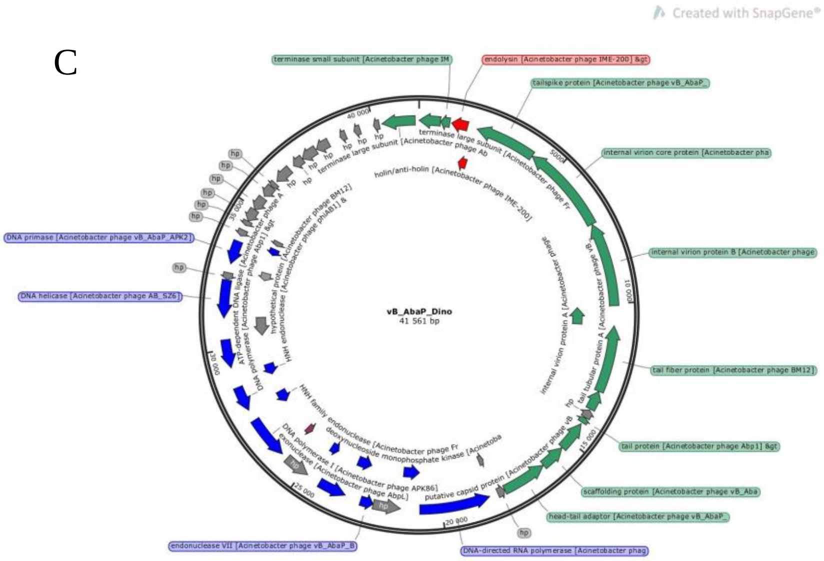
Created with SnapGene®
terminase small subunit [Acinetobacter phage IM
endolysin [Acinetobacter phage IME-200] &gt
tailspike protein [Acinetobacter phage vB_AbaP_
internal virion core protein [Acinetobacter pha
internal virion protein B [Acinetobacter phage
tail fiber protein [Acinetobacter phage BM12]
tail protein [Acinetobacter phage Abp1] &gt
scaffolding protein [Acinetobacter phage vB_Aba
head-tail adaptor [Acinetobacter phage vB_AbaP_
hp
DNA-directed RNA polymerase [Acinetobacter phag
endonuclease VII [Acinetobacter phage vB_AbaP_B
DNA helicase [Acinetobacter phage AB_SZ6]
DNA primase [Acinetobacter phage vB_AbaP_APK2]
terminase large subunit [Acinetobacter phage Ab
holin/anti-holin [Acinetobacter phage IME-200]
terminase large subunit [Acinetobacter phage Fr
internal virion protein A [Acinetobacter phage
tail tubular protein A [Acinetobacter phage vB
putative capsid protein [Acinetobacter phage vB
HNH family endonuclease [Acinetobacter phage Fr
deoxynucleoside monophosphate kinase [Acinetoba
DNA polymerase I [Acinetobacter phage APK86]
exonuclease [Acinetobacter phage AbpL]
ATP-dependent DNA ligase [Acinetobacter phage A
DNA polymerase [Acinetobacter phage Abp1] &gt
hypothetical protein [Acinetobacter phage BM12]
HNH endonuclease [Acinetobacter phage phiAB1] &
vB_AbaP_Dino
41 561 bp
5000
10 000
15 000
20 000
25 000
30 000
35 000
40 000

#### vB_AbaM_PhT2-v2

**Closest blastn hit**: Acinetobacter phage vB_AbaM_PhT2

qcov: 96%, per.ident: 99,62%

**Predicted lifestyle** (phageAI): Virulent 64,45%

**Predicted taxonomy**: Myoviridae; Tegunavirus

**Predicted therapeutic suitability:** no harmful genes detected

**Genes of interest:**

- Anti-CRISPR genes:

- type I-C CRISPR-associated protein Cas8c/Csd1 [Deltaproteobacteria bacterium] Hit_103 in pos 70901-71089

- Endolysin genes:

- holin [Acinetobacter phage vB_AbaM_PhT2] Hit_29 in pos 56659-57387
- baseplate hub subunit and tail lysozyme [Acinetobacter phage AbTZA1] Hit_66 in pos 144781-146568
- M3 family oligoendopeptidase [Phycisphaerales bacterium] Hit_121 in pos 117248-117391
- S8 family serine peptidase [Noviherbaspirillum humi] Hit_122 in pos 120050-120205
- nardilysin-like isoform X1 [Varroa destructor] Hit_328 in pos; reverse 15063-14941
- M20 family metallopeptidase [Bradyrhizobium sp. CCBAU 45394] Hit_361 in pos; reverse 131261-131127
- Chain A, Endolysin [Acinetobacter phage AbTZA1] Hit_517 in pos; reverse 69931-69368

- Lytic cycle related genes:

- homing endonuclease [Acinetobacter phage vB_ApiM_fHyAci03] Hit_5 in pos 12058-12849
- homing endonuclease [Acinetobacter phage vB_AbaM_PhT2] Hit_17 in pos 38836-39621

- baseplate hub subunit and tail lysozyme [Acinetobacter phage AbTZA1]    Hit_66 in pos 144781-146568
- baseplate wedge tail fiber protein connector [Acinetobacter phage vB_AbaM_PhT2]    Hit_69 in pos 153415-154281
- tail collar fiber protein [Acinetobacter phage vB_AbaM_PhT2]    Hit_71 in pos 156757-158295
- homing endonuclease [Acinetobacter phage Maestro] Hit_86 in pos 12605-12757
- homing endonuclease [Acinetobacter phage AbTZA1] Hit_96 in pos 37805-38539
- tail fiber protein proximal subunit [Acinetobacter phage vB_AbaM_PhT2]    Hit_183 in pos 46893-50771
- long tail fiber protein proximal connector [Acinetobacter phage vB_AbaM_PhT2]    Hit_184 in pos 50781-51914
- hinge connector of long tail fiber protein distal connector    Hit_185 in pos 51972-52676
- tail fiber protein [Acinetobacter phage vB_AbaM_PhT2]    Hit_186 in pos 52686-56441
- tail fiber protein [Acinetobacter phage AbTZA1]    Hit_243 in pos; reverse 138957-138391
- metallo-phosphoesterase [Acinetobacter phage vB_AbaM_PhT2]   Hit_259 in pos; reverse 120759-119746
- SegB-like homing endonuclease [Acinetobacter phage Mokit] Hit_356 in pos; reverse 135113-134682
- homing endonuclease [Escherichia phage IME08] Hit_440 in pos; reverse 5249-4572
- Recombination endonuclease VII [Acinetobacter baumannii] Hit_467 in pos; reverse 123376-122894
- SbcC-like subunit of palindrome specific endonuclease [Acinetobacter phage vB_AbaM_PhT2]    Hit_472 in pos; reverse 119149-117467

**Host bacteria:** A72-R (ST636-KL40)

**Genome size:** 169554 bp

**Enzymatic activity**: No

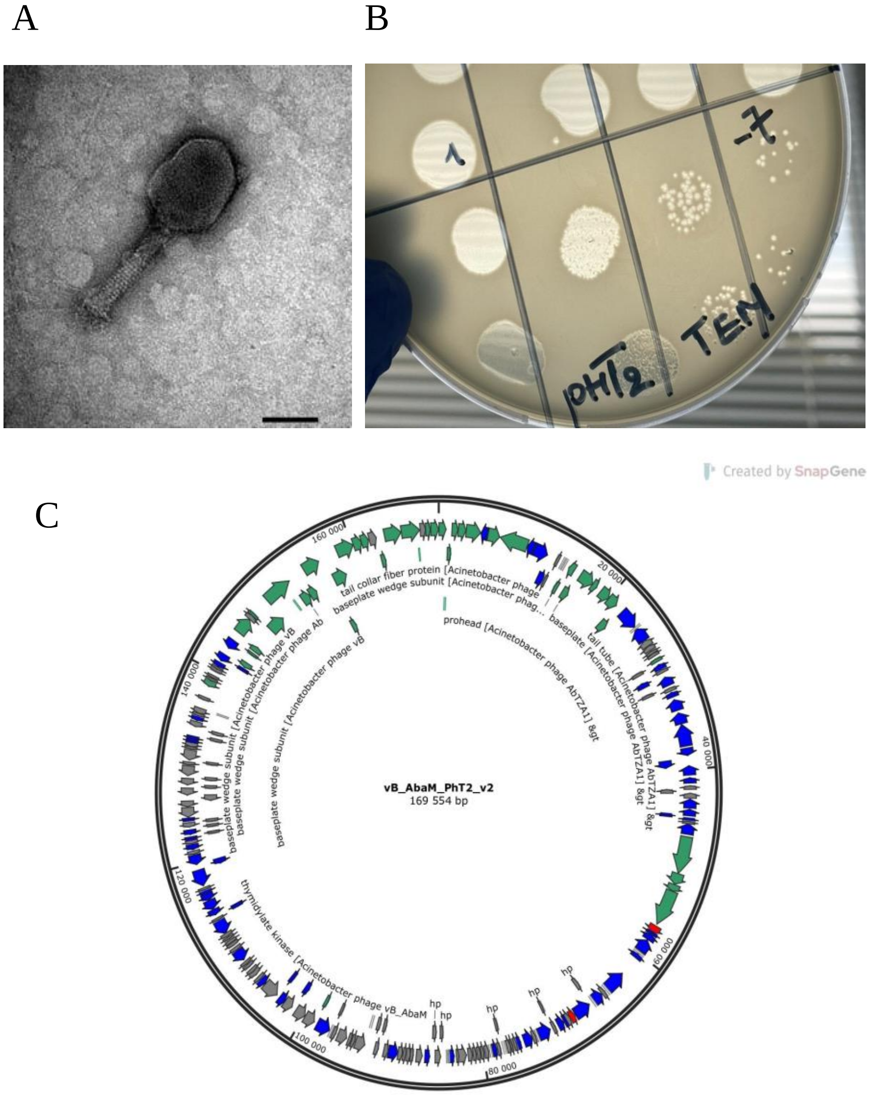

#### vB_AbaP_ABW132

**Closest blastn hit**: Acinetobacter phage AbpL

qcov: 90%, per.ident: 94,70%

**Predicted lifestyle** (phageAI): Lytic 99,71%

**Predicted taxonomy**: Autographiviridae; Friunavirus

**Predicted therapeutic suitability:** no harmful genes detected

**Genes of interest**:

- Anti-CRISPR genes: n.d

- Endolysin genes:

  - endolysin [Acinetobacter phage APK16] Hit_18 in pos 26800-27357
  - M48 family metallopeptidase [Rhizomicrobium electricum] Hit_33 in pos 7448-7600
  - holin/anti-holin [Acinetobacter phage Fri1] Hit_59 in pos 26478-26813

- Lytic cycle related genes:

  - endonuclease [Acinetobacter phage AbpL] Hit_31 in pos 5807-6283

**Host bacteria:** Aci 132 (ST492-KL104)

**Genome size:** 41103bp

**Enzymatic activity**: No

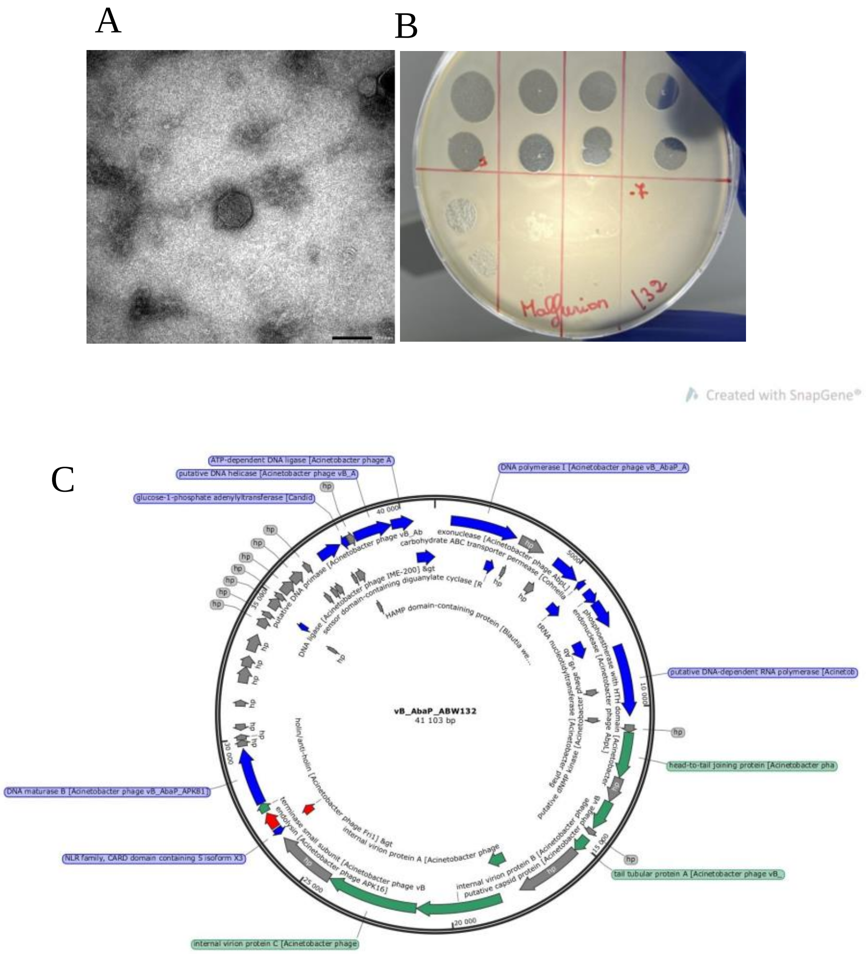
A
B
Malfurion 132
-7
Created with SnapGene®
C
vB_AbaP_ABW132
41 103 bp
ATP-dependent DNA ligase [Acinetobacter phage A
putative DNA helicase [Acinetobacter phage vB_A
glucose-1-phosphate adenylyltransferase [Candid
DNA polymerase I [Acinetobacter phage vB_AbaP_A
putative DNA-dependent RNA polymerase [Acinetob
head-to-tail joining protein [Acinetobacter pha
tail tubular protein A [Acinetobacter phage vB_
internal virion protein C [Acinetobacter phage
NLR family, CARD domain containing 5 isoform X3
DNA maturase B [Acinetobacter phage vB_AbaP_APK81]
exonuclease [Acinetobacter phage AbpL]
carbohydrate ABC transporter permease [Cohnella
putative DNA primase [Acinetobacter phage vB_Ab
DNA ligase [Acinetobacter phage IME-200] &gt
sensor domain-containing diguanylate cyclase [R
HAMP domain-containing protein [Blautia we...
phosphoesterase with HTH domain [Acinetobacter
endonuclease [Acinetobacter phage AbpL]
tRNA nucleotidyltransferase [Acinetobacter phag
putative dNMP kinase [Acinetobacter phage vB
holin/anti-holin [Acinetobacter phage Fri1] &gt
internal virion protein A [Acinetobacter phage
internal virion protein B [Acinetobacter phage
putative capsid protein [Acinetobacter phage vB
terminase small subunit [Acinetobacter phage vB
endolysin [Acinetobacter phage APK16]
hp
5000
10 000
15 000
20 000
25 000
30 000
35 000
40 000

#### vB_AbAM_phiAbaA1_Rocket

**Closest blastn hit**: Acinetobacter phage vB_AbaM_phiAbaA1.

qcov: 98%, perident: 98,44%

**Predicted lifestyle** (phageAI): virulent 89,48%.

**Predicted taxonomy**:  Myoviridae; Saclayvirus

**Genes of interest**:

- Anti-CRISPR genes: n.d.

- Endolysin genes: n.d.

- Lytic cycle related genes:

- mobile endonuclease [Acinetobacter phage vB_AbaM_CP14]    Hit_23 in pos 27538-27867
- baseplate spike [Acinetobacter phage vB_AbaM_phiAbaP1]    Hit_51 in pos 71947-72750
- tail fiber protein [Acinetobacter phage vB_AbaM_phiAbaP1]    Hit_53 in pos 76078-76938
- tail fiber protein [Acinetobacter phage vB_AbaM_phiAbaP1]    Hit_55 in pos 78763-79134
- tail fiber protein [Acinetobacter phage vB_AbaM_phiAbaP1]    Hit_125 in pos 76931-78013
- tail fiber protein [Acinetobacter phage vB_AbaM_phiAbaP1]   Hit_178 in pos 69450-70388
- HNH endonuclease [Acinetobacter phage vB_AbaM_phiAbaP1]    Hit_190 in pos 92721-93341

**Host bacteria:** A127-54 (ST636-KL40)

**Genome size:** 104232 bp

**Enzymatic activity**: No

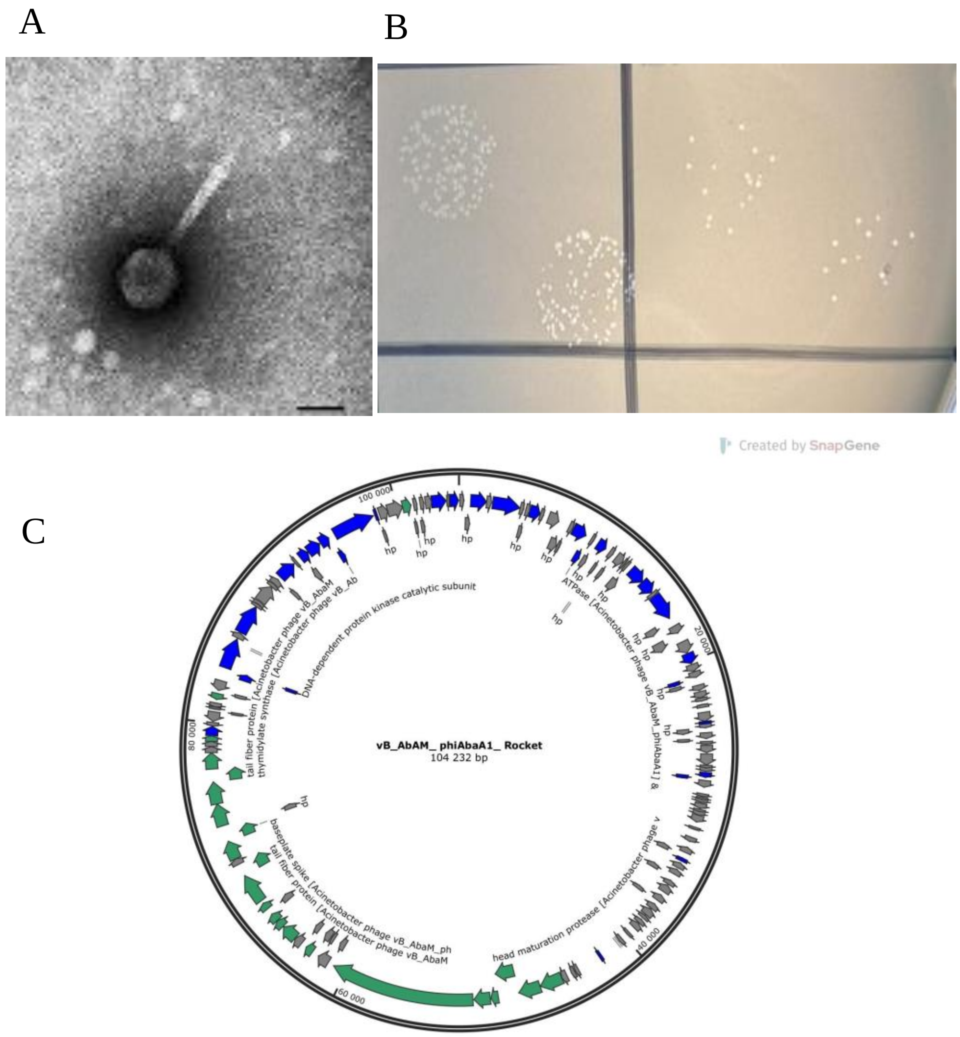

A
B
C
Created by SnapGene
vB_AbAM_ phiAbaA1_ Rocket
104 232 bp
100 000
20 000
40 000
60 000
80 000
hp
DNA-dependent protein kinase catalytic subunit
tail fiber protein [Acinetobacter phage vB_AbaM
thymidylate synthase [Acinetobacter phage vB_Ab
ATPase [Acinetobacter phage vB_AbaM_phiAbaA1] &
head maturation protease [Acinetobacter phage v
baseplate spike [Acinetobacter phage vB_AbaM_ph
tail fiber protein [Acinetobacter phage vB_AbaM

#### vB_AbaM_KissB

**Closest blastn hit**: Acinetobacter phage vB_AbaM_phiAbaA1.

qcov: 92%; perident: 97,78%

**Predicted lifestyle** (phageAI): Lytic 92,12%

**Predicted taxonomy**: Myoviridae; Saclayvirus

**Predicted therapeutic suitability:** no harmful genes detected

**Genes of interest**:

- Anti-CRISPR genes: n.d.

- Endolysin genes: n.d.

- Lytic cycle related genes:

- tail fiber protein [Acinetobacter phage vB_AbaM_phiAbaP1] Hit_164 in pos; reverse 62369-61287
- mobile endonuclease [Acinetobacter phage vB_AbaM_CP14] Hit_202 in pos; reverse 7025-6897
- tail fiber protein [Acinetobacter phage vB_AbaM_phiAbaP1] Hit_216 in pos; reverse 69847-68909
- HNH endonuclease [Acinetobacter phage vB_AbaM_phiAbaP1] Hit_226 in pos; reverse 46618-45998
- HNH homing endonuclease [Acinetobacter phage AM24] Hit_271 in pos; reverse 97242-96826
- baseplate spike [Acinetobacter phage vB_AbaM_phiAbaP1] Hit_284 in pos; reverse 67317-66547
- tail fiber protein [Acinetobacter phage vB_AbaM_phiAbaP1] Hit_286 in pos; reverse 63222-62362
- tail fiber protein [Acinetobacter phage vB_AbaM_phiAbaP1] Hit_288 in pos; reverse 60537-60166

**Host bacteria:** Aci 131(ST2-KL77)

**Genome size:** 102626 bp

**Enzymatic activity:** No

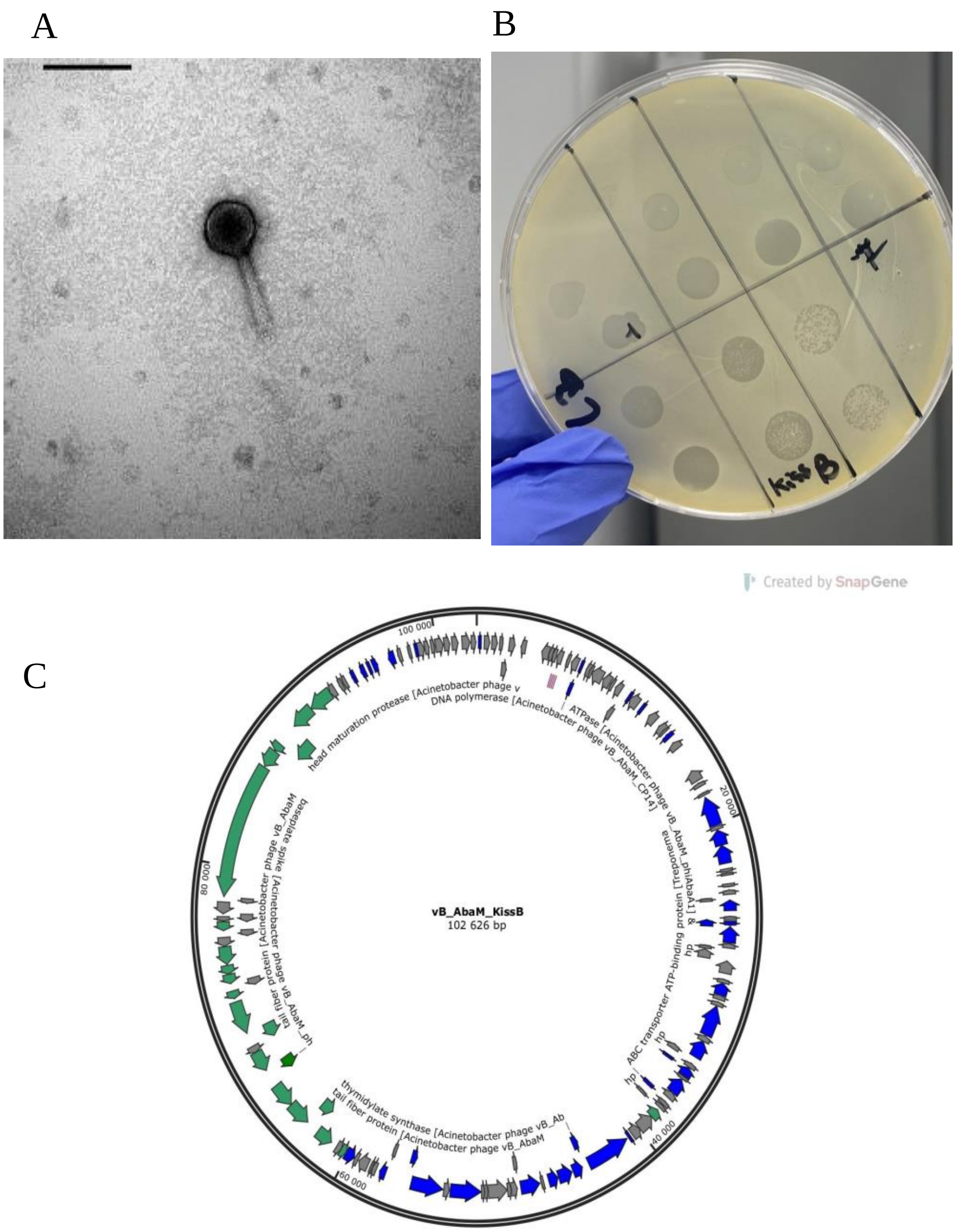

#### vB_AbaM_Konradin-v2

**Closest blastn hit**: Acinetobacter phage Konradin

qcov: 98%, perident: 98,57%

**Predicted lifestyle** (phageAI): Virulent 67.20%

**Predicted taxonomy**: Myoviridae; Tegunavirus

**Predicted therapeutic suitability:** no harmful genes detected

**Genes of interest**:

- Anti-CRISPR genes: n.d.

- Endolysin genes:

- M28 family metallopeptidase [Rhodanobacter glycinis] Hit_6 in pos 4933-5160
- peptidase S41 [Flavobacteriales bacterium] Hit_78 in pos 100315-100443
- putative peptidase inhibitor [Acinetobacter phage vB_AbaM_Konradin] Hit_154 in pos 67040-67315
- putative lytic murein transglycosylase [Acinetobacter phage vB_AbaM_Konradin] Hit_255 in pos 55614-56135
- putative peptidase inhibitor [Acinetobacter phage Stupor] Hit_269 in pos 66450-66761
- endolysin [Acinetobacter phage vB_AbaM_Apostate] Hit_313 in pos 116538-117104
- putative membrane-bound inhibitor of lysozyme [Acinetobacter phage vB_AbaM_Konradin] Hit_318 in pos 122052-122336
- putative glutathionylspermidine synthetase/amidase [Acinetobacter phage AM101] Hit_319 in pos 122400-123668
- aminopeptidase P family protein [Clostridia bacterium] Hit_380 in pos; reverse 101162-101013
- baseplate wedge subunit [Acinetobacter phage vB_ApiM_fHyAci03] Hit_417 in pos; reverse 35942-34125
- holin [Acinetobacter phage vB_AbaM_Apostate] Hit_453 in pos; reverse 132232-131501

- baseplate hub subunit and tail lysozyme [Acinetobacter phage vB_AbaM_Berthold] Hit_490 in pos; reverse 45436-43652
- S8 family serine peptidase [Shewanella algidipiscicola] Hit_552 in pos; reverse 92682-92557

- Lytic cycle related genes:

- endonuclease VII [Acinetobacter phage Abraxas] Hit_45 in pos 67348-67830
- SbcD-like subunit of palindrome specific endonuclease [Acinetobacter phage vB_AbaM_Konradin] Hit_48 in pos 69568-70578
- SbcC-like subunit of palindrome specific endonuclease [Acinetobacter phage vB_AbaM_Konradin] Hit_49 in pos 71440-73125
- DNA binding protein [Acinetobacter phage KARL-1] Hit_88 in pos 107866-108339
- DenB-like DNA endonuclease IV [Acinetobacter phage vB_AbaM_Konradin] Hit_97 in pos 127741-128394
- RNA ligase and tail fiber protein attachment catalyst [Acinetobacter phage vB_AbaM_Konradin] Hit_112 in pos 154231-155376
- ribosomal RNA small subunit methyltransferase A [Spirochaeta sp.] Hit_132 in pos 36629-36814
- putative endonuclease subunit [Acinetobacter phage vB_AbaM_Apostate] Hit_186 in pos 108899-109243
- homing endonuclease [Acinetobacter phage AC4] Hit_231 in pos 15417-16844
- tail fiber chaperone [Acinetobacter phage vB_ApiM_fHyAci03] Hit_247 in pos 48891-49130
- tail fiber protein [Acinetobacter phage vB_AbaM_Berthold] Hit_249 in pos 51327-51893
- methyltransferase [Sorangium cellulosum] Hit_250 in pos 52917-53207
- putative Seg-like homing endonuclease [Acinetobacter phage vB_AbaM_Konradin] Hit_253 in pos 54735-54995
- homing endonuclease [Acinetobacter phage KARL-1] Hit_254 in pos 55050-55487

- HNH endonuclease signature motif containing protein [Calothrix elsteri] Hit_448 in pos; reverse 145270-145133
- tail fiber protein proximal subunit [Acinetobacter phage KARL-1] Hit_450 in pos; reverse 142537-138617
- long tail fiber protein proximal connector [Acinetobacter phage KARL-1] Hit_451 in pos; reverse 138607-137477
- N-6 DNA methylase [Nitrobacter hamburgensis] Hit_458 in pos; reverse 128032-127901
- baseplate wedge tail fiber protein connector [Acinetobacter phage vB_ApiM_fHyAci03] Hit_493 in pos; reverse 36808-35942
- tail collar fiber protein [Acinetobacter phage vB_AbaM_Lazarus] Hit_495 in pos; reverse 33466-31940
- hinge connector of long tail fiber protein distal connector [Acinetobacter phage KARL-1] Hit_529 in pos; reverse 137553-136708
- tail fiber protein [Acinetobacter phage KARL-1] Hit_530 in pos; reverse 136695-132844
- HNH endonuclease [Acinetobacter phage vB_AbaM_Berthold] Hit_556 in pos; reverse 81702-80578
- tRNA (guanosine(37)-N1)-methyltransferase TrmD [Anaerolineales bacterium] Hit_569 in pos; reverse 58149-57991

**Host bacteria:** Aci 222 (ST636-KL40)

**Genome size:** 164573 bp

**Enzymatic activity**: No

A B

C

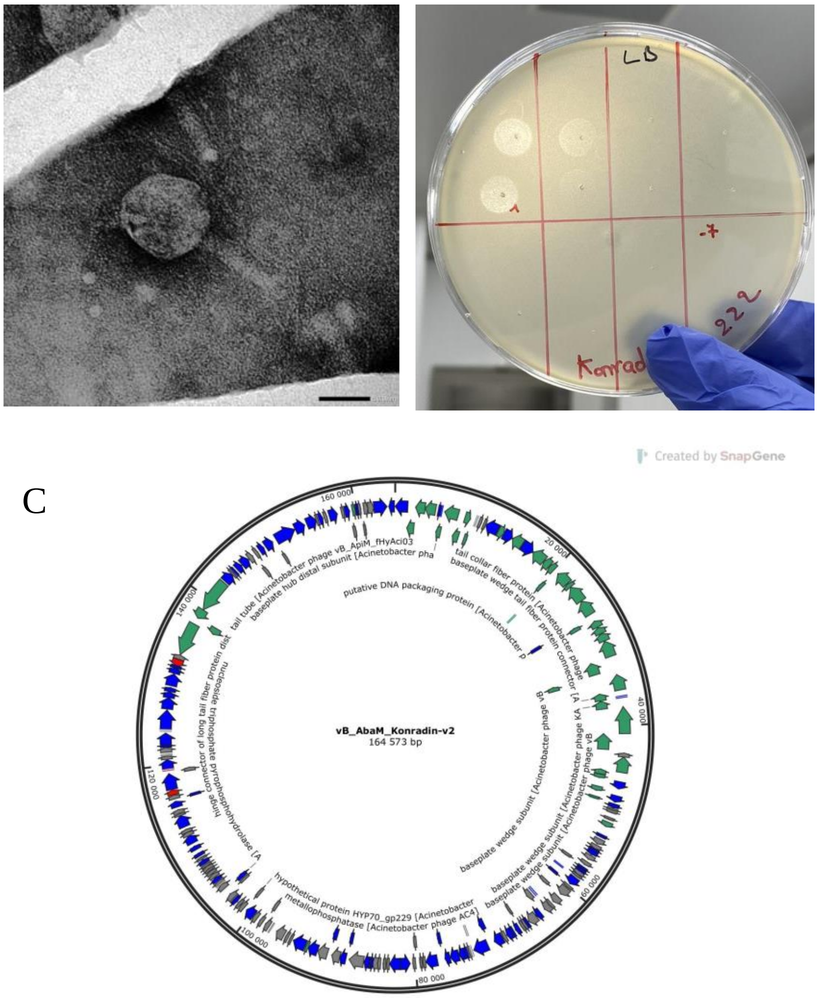
LD
-7
Konrad
222
Created by SnapGene
vB_AbaM_Konradin-v2
164 573 bp
160 000
20 000
40 000
60 000
80 000
100 000
120 000
140 000
tail tube [Acinetobacter phage vB_ApiM_fHyAci03
baseplate hub distal subunit [Acinetobacter pha
tail collar fiber protein [Acinetobacter phage
baseplate wedge tail fiber protein connector [A
putative DNA packaging protein [Acinetobacter p
baseplate wedge subunit [Acinetobacter phage vB
baseplate wedge subunit [Acinetobacter phage KA
baseplate wedge subunit [Acinetobacter phage vB
hypothetical protein HYP70_gp229 [Acinetobacter
metallophosphatase [Acinetobacter phage AC4]
nucleoside triphosphate pyrophosphohydrolase [A
hinge connector of long tail fiber protein dist

#### vB_AbaM_Navy4-v2

**Closest blastn hit**: Acinetobacter phage AB-Navy4

qcov: 97%, per.ident: 98,20%

**Predicted lifestyle** (phageAI): Virulent 74,08%

**Predicted taxonomy**: Myoviridae; Tegunavirus

**Predicted therapeutic suitability:** no harmful genes detected

**Genes of interest:**

- Anti-CRISPR genes: n.d.

- Endolysin genes:

- putative peptidase inhibitor [Acinetobacter phage vB_AbaM_Berthold] Hit_1 in pos 862-1161
- endolysin [Acinetobacter phage vB_ApiM_fHyAci03] Hit_39 in pos 51718-52284
- putative peptidase inhibitor [Acinetobacter phage vB_AbaM_Berthold] Hit_115 in pos 1139-1414
- putative membrane-bound inhibitor of lysozyme [Acinetobacter phage vB_AbaM_Berthold] Hit_173 in pos 58280-58564
- glutathionylspermidine synthase [Acinetobacter phage vB_AbaM_Berthold] Hit_174 in pos 58628-59896
- M28 family metallopeptidase [Rhodanobacter glycinis] Hit_209 in pos 104927-105154
- baseplate wedge subunit [Acinetobacter phage vB_ApiM_fHyAci03] Hit_210 in pos 108737-109135
- putative lytic murein transglycosylase [Acinetobacter phage vB_AbaM_Lazarus] Hit_242 in pos 157895-158416
- putative peptidase inhibitor [Acinetobacter phage Stupor] Hit_250 in pos 549-860
- holin [Acinetobacter phage vB_AbaM_Apostate] Hit_408 in pos; reverse 68458-67727

- aminopeptidase P family protein [Clostridia bacterium] Hit_508 in pos; reverse 34806-34657
- baseplate hub subunit and tail lysozyme [Acinetobacter phage vB_AbaM_Berthold] Hit_531 in pos; reverse 146831-145047

- Lytic cycle related genes:

- endonuclease VII [Acinetobacter phage vB_ApiM_fHyAci03]  Hit_2 in pos 1447-1929
- SbcD-like subunit of palindrome specific endonuclease [Acinetobacter phage vB_ApiM_fHyAci03]  Hit_5 in pos 3667-4677
- SbcC-like subunit of palindrome specific endonuclease [Acinetobacter phage vB_AbaM_Konradin]  Hit_6 in pos 5539-7224
- DNA binding protein [Acinetobacter phage vB_AbaM_Berthold] Hit_31 in pos 42559-43032
- putative endonuclease subunit [Acinetobacter phage vB_AbaM_Apostate] Hit_33 in pos 44083-44427
- homing endonuclease [Acinetobacter phage AM101]  Hit_83 in pos 116206-117636
- tail fiber chaperone [Acinetobacter phage vB_ApiM_fHyAci03]  Hit_100 in pos 151093-151332
- tail fiber protein [Acinetobacter phage vB_AbaM_Konradin] Hit_103 in pos 153556-154122
- putative HNH endonuclease [Acinetobacter phage Stupor] Hit_165 in pos 45755-46708
- RNA ligase and tail fiber protein attachment catalyst [Acinetobacter phage vB_AbaM_Kimel]  Hit_194 in pos 89843-90988
- MAG: peptide chain release factor N(5)-glutamine methyltransferase [Candidatus Omnitrophica bacterium]  Hit_220 in pos 117116-117301

- homing endonuclease [Acinetobacter phage vB_AbaM_Konradin] Hit_234 in pos 147911-148561
- putative Seg-like homing endonuclease [Acinetobacter phage vB_AbaM_Konradin] Hit_240 in pos 157016-157276

- homing endonuclease [Acinetobacter phage vB_AbaM_Berthold] Hit_241 in pos 157331-157768
- putative homing endonuclease [Acinetobacter phage Stupor] Hit_293 in pos 53892-54740
- DNA endonuclease IV [Acinetobacter phage AB-Navy4] Hit_298 in pos 63966-64619
- homing endonuclease [Acinetobacter phage vB_AbaM_Berthold] Hit_338 in pos 148233-148394
- homing endonuclease [Acinetobacter phage Maestro] Hit_387 in pos; reverse 110860-110708
- long tail fiber distal hinge connector [Acinetobacter phage AB-Navy97] Hit_406 in pos; reverse 73129-72284
- tail fiber protein [Acinetobacter phage vB_ApiM_fHyAci03] Hit_407 in pos; reverse 72271-68534
- homing endonuclease [Acinetobacter phage AB-Navy4] Hit_506 in pos; reverse 38847-37909
- baseplate wedge tail fiber protein connector [Acinetobacter phage vB_ApiM_fHyAci03] Hit_534 in pos; reverse 138203-137337
- short tail fibers protein [Acinetobacter phage Stupor] Hit_536 in pos; reverse 134861-133335
- homing endonuclease [Acinetobacter phage vB_ApiM_fHyAci03] Hit_554 in pos; reverse 111407-110613
- HNH endonuclease signature motif containing protein [Calothrix elsteri] Hit_567 in pos; reverse 80861-80724
- long tail fiber proximal subunit [Acinetobacter phage AB-Navy4] Hit_569 in pos; reverse 78137-74193
- long tail fiber hinge connector [Acinetobacter phage AB-Navy97] Hit_570 in pos; reverse 74183-73053
- mobile endonuclease [Acinetobacter phage Abraxas] Hit_594 in pos; reverse 15800-14676

**Host bacteria:** Aci 132 (ST492-KL104)

**Genome size:** 167167 bp

**Enzymatic activity**: No

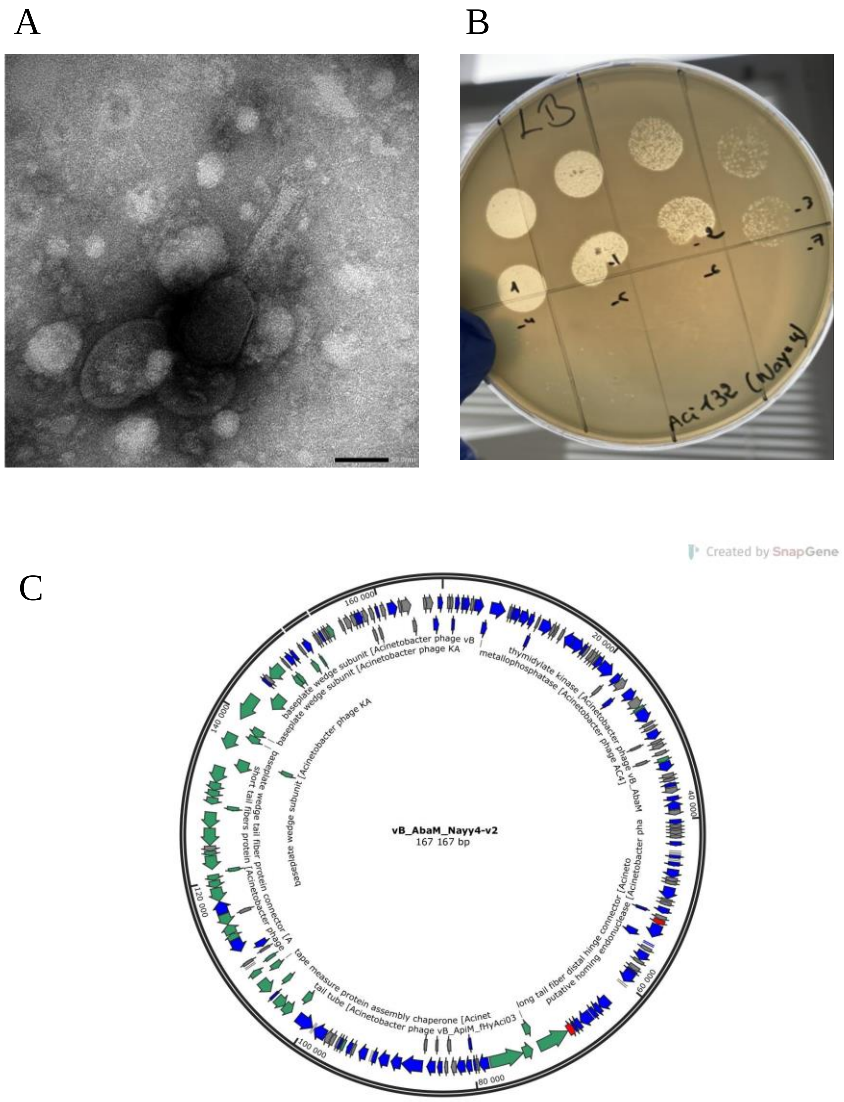
